# Efficient coding explains neural response homeostasis and stimulus-specific adaptation

**DOI:** 10.1101/2023.10.29.564616

**Authors:** Edward James Young, Yashar Ahmadian

## Abstract

In the absence of adaptation, the average firing rate of neurons would rise or drop when changes in the environment make their preferred stimuli more or less prevalent. However, by adjusting the responsiveness of neurons, adaptation can yield firing rate homeostasis and stabilise the average rates of neurons at fixed levels, despite changes in stimulus statistics. In sensory cortex, adaptation is typically also stimulus specific, in that neurons reduce their responsiveness to over-represented stimuli, but maintain or even increase their responsiveness to stimuli far from over-represented ones. Here, we present a normative explanation of firing rate homeostasis grounded in the efficient coding principle. Specifically, we show that this homeostasis can arise when neurons adapt their responsiveness to optimally mitigate the effect of neural noise on population coding fidelity, at minimal metabolic cost. Unlike previous efficient coding theories, we formulate the problem in a computation-agnostic manner, enabling our framework to apply far from the sensory periphery. We then apply this general framework to Distributed Distributional Codes, a specific computational theory of neural representations serving Bayesian inference. We demonstrate how homeostatic coding, combined with such Bayesian neural representations, provides a normative explanation for stimulus-specific adaptation, widely observed across the brain, and how this coding scheme can be accomplished by divisive normalisation with adaptive weights. Further, we develop a model within this combined framework, and, by fitting it to previously published experimental data, quantitatively account for measures of stimulus-specific and homeostatic adaption in the primary visual cortex.

## 1 Introduction

Neural population responses, and thus the computations on sensory inputs represented by them, are corrupted by noise. The extent of this corruption can, however, be modulated by changing neural gains. When the gain of a neuron increases, the strength of its response noise, or trial-to-trial variability, typically grows sublinearly and more slowly than its average response; for example, for Poisson-like firing, response noise grows as the square root of the mean response. The neuron can therefore increase its signal-to-noise ratio by increasing its gain or responsiveness. However, this increase in coding fidelity comes at the cost of an elevated average firing rate, and thus higher metabolic energy expenditure. From a normative perspective, coding fidelity and metabolic cost are thus two conflicting forces.

We start this study by asking what is the optimal way of adjusting neural gains in order to combat noise with minimal metabolic cost, *irrespective* of the computations represented by the neural population. As we will see, the answer depends on the prevailing stimulus statistics. We will thus address the following more specific question: given a neural population with *arbitrary* tuning curve shapes and general noise distribution, how should neurons optimally adjust their gains depending on the stimulus statistics prevailing in the environment? To address this question, we use efficient coding theory as our normative framework (Attneave, 1954; Nadal and Parga, 1999; Laughlin, 1981; Barlow, 2012; Linsker, 1988; Ganguli and Simoncelli, 2014; Wei and Stocker, 2015; Atick and Redlich, 1990). Efficient coding theories formalise the problem of optimal coding subject to biological constraints and costs, and by finding the optimal solution make predictions about the behavior of nervous systems (which are postulated to have been approximately optimised, *e*.*g*., via natural selection). Concretely, the Infomax Principle (Linsker, 1988; Atick and Redlich, 1990; Ganguli and Simoncelli, 2014; Wei and Stocker, 2015) states that sensory systems optimise the mutual information between a noisy neural representation and an external stimulus, subject to metabolic constraints, *e*.*g*., on the energy cost of spiking activity. To that end, the neural system should exploit the statistics of stimuli in its local environment (Simoncelli and Olshausen, 2001). Efficient coding theories can therefore predict how sensory systems should optimally *adapt* to changes in environmental stimulus statistics. There is, however, a key difference between the approach we will adopt and classic applications of the efficient coding principle. In most applications, this principle has been applied to systems located early in the sensory stream, and information transmission is taken to be the exclusive goal of the system. Accordingly, the entire input-output transformations performed by the system is assumed to have been optimised (under metabolic or anatomical constraints) purely for this goal. Based on this assumption, such theories typically make predictions about patterns of neural selectivity, including the shapes and arrangement of neural tuning curves. However, for general brain regions located deeper within the sensory-motor pathway, information transmission is far from an adequate charactrisation of the system’s computational goals. For example, many neural systems perform computations which require systematically discarding information contained in the stimulus input. The discarded information may correspond to nuisance variables, or be irrelevant to the behavioural tasks faced by the organism. Here, we make the key assumption that the shapes and arrangement of tuning curves are determined by computational goals — beyond mitigating the effect of noise on coding fidelity — to which our efficient coding framework remains agnostic. We therefore do not optimise these aspects of the population code and allow them to be arbitrary. On the other hand, we assume the neural gains (which set the scale of the tuning curves) are adapted and optimised purely for the purpose of fighting noise corruption at minimal metabolic cost, regardless of the system’s computational goals. Conceptually, this division of labour is akin to the separation, in digital computers, between error correction (which is indispensable in practice) and general computations, and the need for the former to be performed in a manner agnostic to the ongoing computations.

Suppose an environmental shift makes a stimulus feature more prevalent. Neurons sensitive to that feature will then see their preferred feature more often, and thus their average firing rate will initially increase. However, typically, those neurons gradually adapt by reducing their responsiveness or gain (Solomon and Kohn, 2014; Clifford et al., 2007; Benucci et al., 2013). Such gain adaptation can lead to *firing rate homeostasis* (Desai, 2003; Turrigiano and Nelson, 2004; Maffei and Turrigiano, 2008; Hengen et al., 2013), whereby adaptation brings the average firing rate back to the level prior to the environmental shift (Fig. 1A). Thus, under such homeostatic adaptation, neuronal populations maintain a constant stimulus-averaged firing rate in spite of changes to the environment (Benucci et al., 2013; Hengen et al., 2013; Maffei and Turrigiano, 2008).

**Figure 1:**
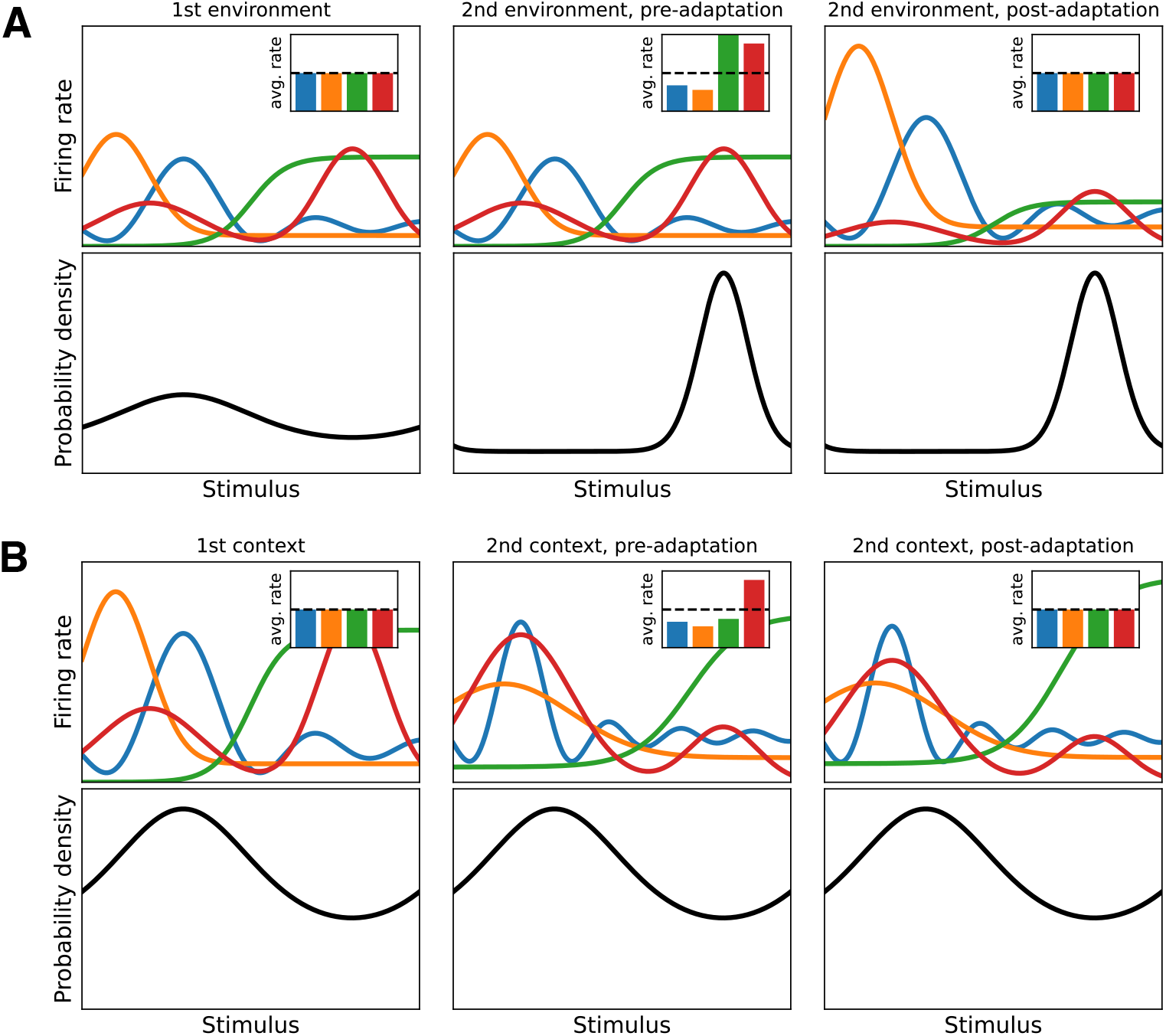
Firing rate homeostasis via gain adaptation (cartoon examples). (A) Gain adaptation after a shift in environmental statistics, without changes in tuning curve shapes. The bottom row shows the stimulus distributions in the first (left column) and the second(middle and right columns) environments. The top row shows the tuning curves of the same four neurons (main plots) and their mean firing rates (insets) when adapted to the first (left) and the second (right) environments, as well as immediately after the transition, but before adaptation to the new stimulus statistics (middle). Immediately after the transition the mean firing rates differ from their values in the first environment (middle and left insets). Neurons adapt to the new environment solely by independently adjusting their gains, *i*.*e*., by scaling their tuning curves up or down (correspondingly, the tuning curves of the same color in the middle and right columns — which belong to the same neuron — are scaled versions of each other and have identical shape), to a degree appropriate for returning their stimulus averaged firing rates to the level before the environmental shift (right and left insets). (B) Gain adaptation after a change in tuning curve shapes, due, *e*.*g*., to a change in the system’s computational goals. The plots have the same format as in (A). In this case, the stimulus distribution (bottom row) does not change between the two contexts, but the tuning curves have different shapes in the second context (top row, second and third columns vs. the first). Again, the average firing rates change immediately after the transition (middle vs. left insets), but the neurons adapt their gains to the right degree to return their rates to the level before the context changed (right inset).

There are multiple levels at which homeostatic adaptation can be observed. Firstly, there is population homeostasis, in which the stimulus-average firing rate of an entire population of neurons remains constant, without individual neurons necessarily holding their rates constant (Slomowitz et al., 2015; Sanzeni et al., 2023). Secondly, there is what we term *cluster homeostasis*. In this form of homeostasis, stimulus-average firing rate of groups or clusters of neurons with similar stimulus tuning remains stable under environmental shifts (Benucci et al., 2013), but the firing rate of individual neurons within a cluster can change. Lastly, in the strongest form, homeostasis can occur at the level of individual neurons, in which case the firing rate of each individual neuron is kept constant under changes in environment statistics (Marder and Prinz, 2003). Previous normative explanations for firing rate homeostasis often focus on the necessity of preventing the network from becoming hypo- or hyperactive (Turrigiano and Nelson, 2004; Maffei and Turrigiano, 2008; Keck et al., 2013; Hengen et al., 2013). Although this might explain homeostasis at the population level, it does not adequately explain homeostasis at a more fine-grained level. As we will show in this paper, the answer to the normative question that we posed above provides an explanation for firing rate homeostasis at such a finer level. Specifically, we will show that, for a wide variety of neural noise models (and under conditions that we argue do obtain in the visual cortex), optimal adjustment of neural gains to combat the effect of coding noise predicts firing rate homeostasis at the level of clusters of similarly-tuned neurons.

Having shown the optimality of homeostatic gain adaptation regardless of the computational goals of the population (which we have assumed determine its tuning curve shapes), we then apply our framework to *Distributed Distributional Codes (DDC)* (Vertes and Sahani, 2018), as a specific computational theory for neural implementation of Bayesian inference. Combining DDC theory (which determines tuning curve shapes) with our normative results (on optimal adjustments of neural gains) yields what we call *homeostatic DDC*. We will show that homeostatic DDCs explain the typical finding that sensory adaptations are stimulus specific (Kohn, 2007; Schwartz et al., 2007; Taaseh et al., 2011). In stimulus specific adaptation (SSA), the suppression of response is greater for test stimuli that are closer to an over-represented adaptor stimulus than for stimuli further away (to which the neuron may even respond more strongly after adaptation). In particular, this causes a repulsion of tuning curves away from the adaptor (Movshon and Lennie, 1979; Müller et al., 1999; Dragoi et al., 2000, 2002; Benucci et al., 2013). We will show that homeostatic DDC are able to account for SSA effects which cannot be fully accounted for in previous efficient coding frameworks (Wei and Stocker, 2015; Ganguli and Simoncelli, 2014; Snow et al., 2016), and we will use them to quantitatively model homeostatic SSA as observed in V1, by re-analysing previously published data (Benucci et al., 2013). This model relies on a special class of homeostatic DDC which we term *Bayes-ratio coding*. We show that Bayes-ratio coding has attractive computational properties: it can be propagated between populations without synaptic weight adjustments, and it can be achieved by divisive normalisation (which has been called a canonical neural operation) with adaptive weights (Carandini and Heeger, 2012; Westrick et al., 2016).

We start the next section by introducing our efficient coding framework for addressing the normative question posed above. We show numerically that our framework predicts that the firing rate of clusters of similarly tuned neurons should remain constant despite shifts in environmental stimulus statistics, with individual neurons free to shuffle their average rates. This result is shown to hold across a diverse set of noise models, demonstrating the robustness of the optimality of homeostasis to the precise nature of neural response noise. We additionally demonstrate that our framework can account for a wide distribution of stimulus-averaged single-neuron firing rates as observed in cortex. We then provide an analytic argument that demonstrates why, within a certain parameter regime, homeostasis arises from our optimisation problem, and show that cortical areas, in particular the primary visual cortex (V1), are likely to be within this parameter regime. We then numerically validate the quality of our analytic approximations to the optimal solution. Lastly, we apply our theory to DDC representational codes, showing how homeostatic DDC can account for stimulus specific adaptation effects observed experimentally.

## 2 Results

### 2.1 Theoretical framework

The central question we seek to address is how the presence of response noise affects optimal neural coding strategies. Specifically, we consider how a neural population should distribute activity in order to best mitigate the effect of neural noise on coding fidelity, subject to metabolic constraints, irrespective of the computations it performs on sensory inputs (which we do not optimise, but take as given). We consider a population of *K* neuronal units,^1^ responding to the (possibly high-dimensional) stimulus ***s*** according to their tuning curves ***h***(***s***). More precisely, we assume that our population engages in rate coding using time bins of a fixed duration, and denote the vector of joint population spike counts in a coding interval by ***n*** = (*n*_1_, …, *n*_*K*_). Single-trial neural responses, ***n***, are taken to be noisy emissions from the neural tuning curves, ***h***(***s***), according to a noise distributions ***n|s*** *~ P*_noise_(***n***|***s***), subject to the constraint that the trial-averaged response 𝔼[***n*** | ***s***] equals ***h***(***s***). We consider a number of different noise models, including correlated and power-law noise, showing that our main results are robust to specific assumptions about the nature of noise in neural systems. The sensory environment is characterised by a stimulus distribution *P* (***s***); accordingly, an “environmental shift” corresponds to a change in *P* (***s***), making certain stimuli more or less prevalent.

The central assumption of our efficient coding framework is that, while the shapes of neural tuning curves are determined by the computational goals of the circuit, the gains of those tuning curves are adapted purely for the purpose of fighting noise corruption at minimal metabolic cost, regardless of the computational goals. To this end, we adopt a shape-amplitude decomposition of the neural tuning curves. The tuning curve of the *a*-th unit, *h*_*a*_(***s***), is factorised into a *representational curve*, Ω_*a*_(***s***), and a *gain, g*_*a*_:

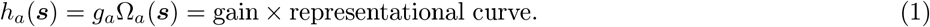

Fig. 3A demonstrates the effect of changing the gain while keeping the representational curve, and therefore the tuning curve shape, fixed.^2^ The representational curve can be thought of as a normalised tuning curve; in fact, to make the definition of gain unambiguous, we define Ω_*a*_(***s***) to be the tuning curve normalised by its peak value, *i*.*e*., Ω_*a*_(***s***) *≡ h*_*a*_(***s***)*/* max *h*_*a*_ (Fig. 3A inset). Correspondingly, the gain of a tuning curve is simply its maximum value, *g*_*a*_ = max_***s***_ *h*_*a*_(***s***).

We take the shape of the tuning curve, captured by Ω_*a*_, to encode the computations represented by the neural population, and thus to be determined by those computational goals. These computational goals, and hence the Ω_*a*_ may change across environments or contexts (*e*.*g*., corresponding to different tasks), but our efficient coding theory will not dictate how they should change (see Fig. 1B, and Section 2.7 for an example). Rather, we treat the representational curves, Ω_*a*_, as given, and do not optimise them within our efficient coding framework. In this respect, we deviate from the assumption of many classic studies based on the efficient coding principle, wherein information transmission is taken to be the sole computational goal of the system (usually located early in the sensory stream), and therefore the input-output transformation performed by the system is assumed to have been optimised purely for this purpose. General computations, however, often involve discarding information contained in the stimulus, that, *e*.*g*., constitute nuisance variables or may be irrelevant to behavioural tasks; this is the rationale for taking the Ω_*a*_ as given and not optimising them purely based on information transmission criteria. In line with this rationale, we do not make any prior assumptions about or put any constraints on the shapes of the representational curves: the Ω_*a*_ can be any complex (*e*.*g*., multi-modal, discontinuous, or heterogeneous) set of functions of the possibly high-dimensional stimulus, and can thus represent any computation. This makes our treatment more generally applicable than most other efficient coding frameworks which place tight constraints on the shapes and configuration of the tuning curves (see the Discussion for a more detailed comparison of our approach with those studies). In particular, this generality enables our theory to apply to populations located deep in the processing pathway, and not just to primary sensory neurons. For example, the representational curves, Ω_*a*_, could map the normalised response of face-detecting neurons in the inferotemporal cortex as a function of the stimulus, ***s***, which could be taken as the high-dimensional raster of RGB values in an image shown to the animal.

Our framework, relating environmental stimuli to neural tuning curves and finally to noisy neural responses, and the respective computational roles of the gains and representational curves, is summarised by the diagram in Fig. 2.

**Figure 2:**
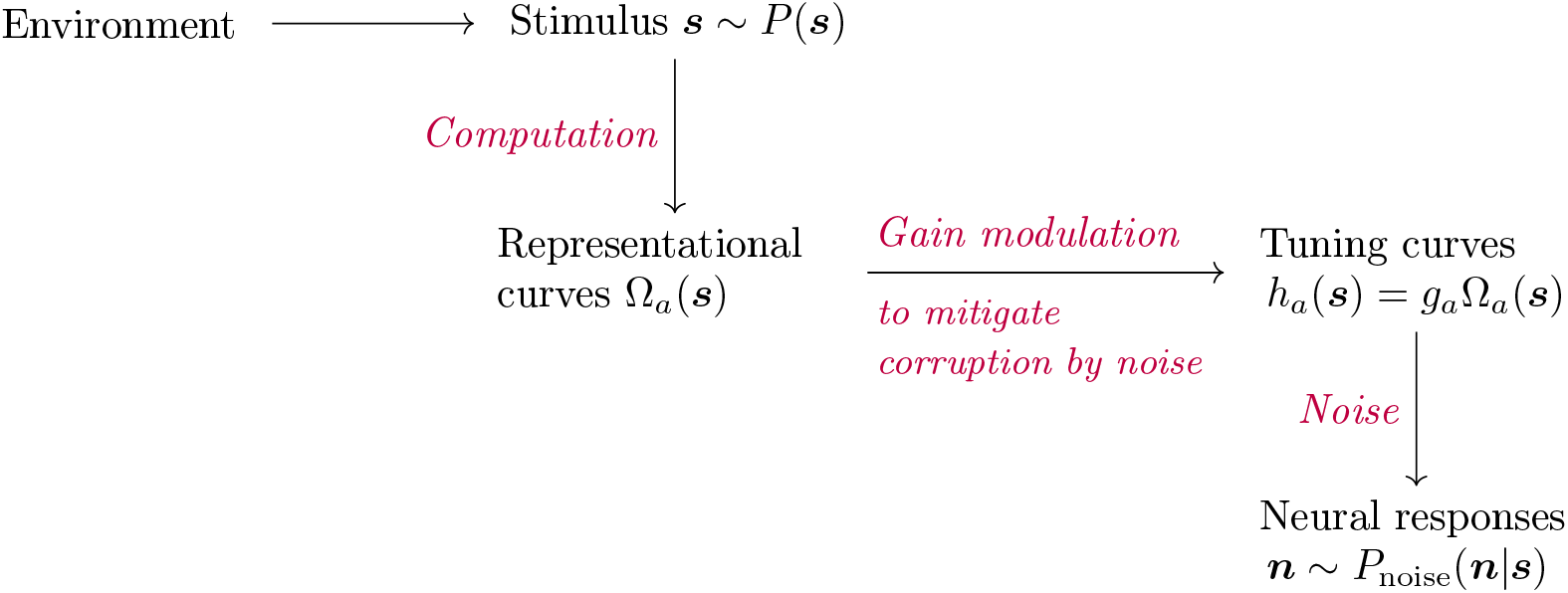
The assumptions and the generative model underlying the theoretical framework. The environment gives rise to a stimulus distribution *P* from which the stimulus ***s*** is drawn. Conceptually, the brain performs some computation on ***s***, the result of which are represented by neurons via their *normalised* tuning curves, Ω_*a*_(***s***), which we thus refer to as representational curves. These are multiplied by adaptive gains, *g*_*a*_, to yields the actual tuning curves *h*_*a*_(***s***). The single-trial neural responses are noisy emissions based on *h*_*a*_(***s***). We assume the only function of gain modulation is to mitigate the eventual corruption of computational results by neural noise, subject to metabolic energy constraints.

We theorise that units adapt their gains to maximise an objective function that trades off the metabolic cost of neural activity with the information conveyed by the responses (Levy and Baxter, 1996; Ganguli and Simoncelli, 2014). Informally

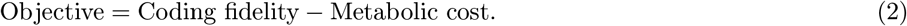

As is common in efficient coding frameworks (see *e*.*g*., Atick and Redlich (1990); Ganguli and Simoncelli (2014)) we assume the main contributor to metabolic cost is the energy cost of emitting action potentials. Thus we take the metabolic cost term in the objective to be proportional to 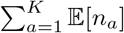, the total average spike count fired by the population. On the other hand, a natural and common choice (up to approximations) for the coding fidelity is the mutual information between the stimulus and response, *I*(***n***; ***s***). However, since in general the analytic maximisation of mutual information is intractable, in the tradition of efficient coding theory (Brunel and Nadal, 1998; Ganguli and Simoncelli, 2014; Linsker, 1988), we will instead optimise an approximate surrogate for mutual information. We decompose the mutual information as *I*(***s***; ***n***) = *H*[***n***] − *H*[***n*** | ***s***]. The marginal entropy term *H*[***n***] can be upper bounded by the entropy of a Gaussian with the same covariance,

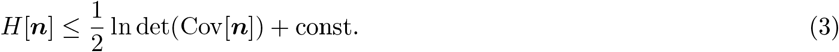

Substituting the right hand side for the marginal entropy term in the mutual information, gives us an upper bound for the latter. Using this upper bound on mutual information as our quantification of “coding fidelity”, yields the objective function

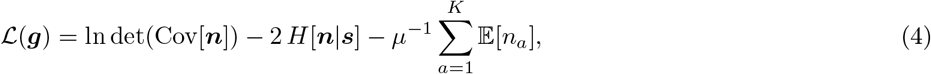

where the constant *µ >* 0 controls the information-energy trade-off. This trade-off is illustrated in Fig. 3B. To summarise, we assume that the optimal gains maximise the objective Eq. (4)

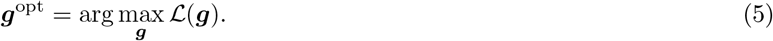

**Figure 3:**
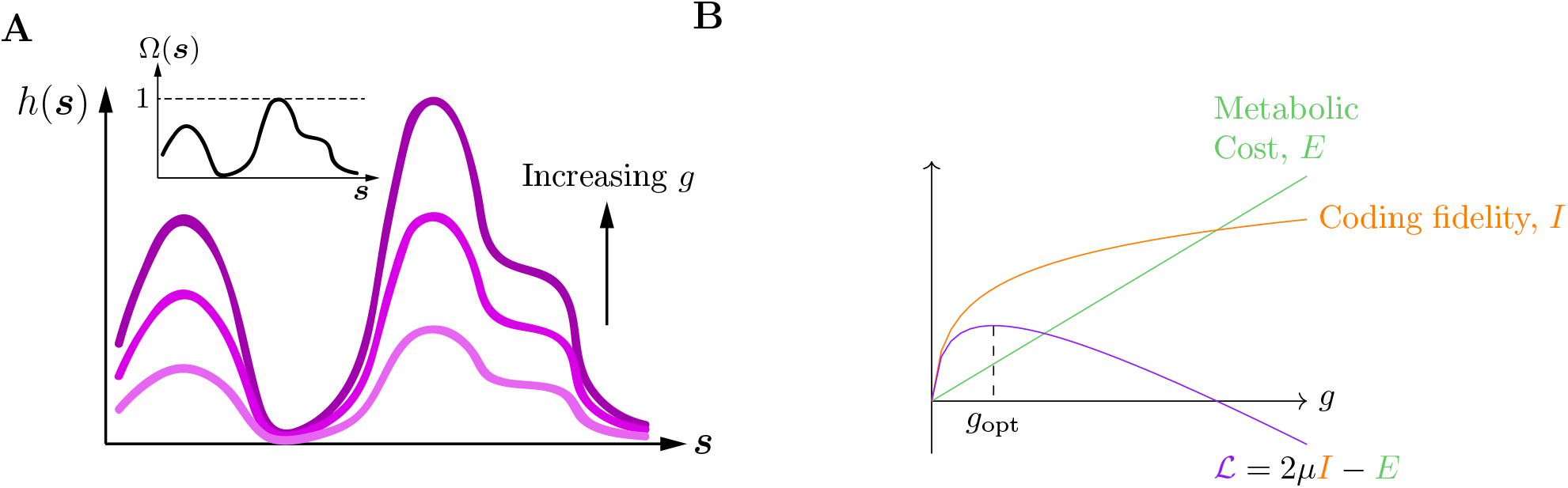
The effect of changing the gain on tuning curves and different components of the objective function. (A) As the gain, *g*, of a neuron is increased, the shape of its tuning curve remains the same, but all firing rates are scaled up. The inset shows the gain-independent representational curve, which we define as the tuning curve normalised to have a peak value of 1. (This cartoon shows a one-dimensional example, but our theory applies to general tuning curve shapes and joint configurations of population tuning curves, on high-dimensional stimulus spaces.) (B) Cartoon representation of the efficient coding objective function. Optimal neural gains maximise an objective function, ℒ, which is the weighted difference between mutual information, *I*, capturing coding fidelity, and metabolic cost, *E*, given by average population firing rate.

Note that because the Ω_*a*_ depend deterministically on the stimulus ***s***, and in turn fully determine the statistics of *n*_*a*_, we have the mutual information identity *I*(***n***; ***s***) = *I*(***n***; **Ω**) (this identity also holds for the upper bound on the mutual information that we are using). This means that the coding fidelity term in the objective function can equivalently be interpreted, without any reference to the low-level stimulus, as (an upper bound on) the mutual information between the population’s noisy output and its ideal, noise-free outputs, **Ω**, as specified by the computational goals of the circuit.

### 2.2 Optimal adaptation of gains leads to rate homeostasis across a variety of noise models

We now investigate the consequences of optimal gain adaptation, according to Eqs. (4)–(5), for the statistics of population response. As our hypothesis is that gains adapt in order to optimally mitigate the effect of neural noise on coding fidelity (with minimal energy expenditure), ideally we would like our conclusions to be robust to the model of noise. We thus consider a broad family of noise models which allow for general noise correlation structure, as well as different (sub- or super-Poissonian) scalings of noise strength with mean response. Specifically, we assume that conditioned on the stimulus, the noisy population response, ***n***, has a normal distribution, with mean ***h***(***s***), and stimulus-dependent covariance matrix

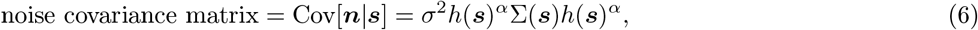

where *h*(***s***) is the diagonal matrix with the components of the tuning curve vector, ***h***(***s***), on its diagonal (more generally, we adopt the convention that the non-bold and index-free version of a bold symbol, representing a vector, denotes a diagonal matrix made from that vector), the noise strength *σ* scales the overall strength of noise, and 0 *< α <* 1 is a parameter that controls the power-law scaling of noise power with mean response ***h***(***s***). Finally, Σ(***s***) is the noise correlation matrix which can depend on the stimulus, but is independent of the gains. The noise model Eq. (6) can equivalently be characterised by (1) the noise variance of spike count *n*_*a*_ is given by *σ*^2^*h*_*a*_(***s***)^2*α*^, and (2) the noise correlation coefficient between *n*_*a*_ and *n*_*b*_ is given by Σ_*ab*_(***s***), independently of the neural gains. Thus the strength (standard deviation) of noise scales sublinearly, as gain to the power *α*. Note that the case *σ* = 2*α* = 1 corresponds to Poisson-like noise with variance of *n*_*a*_ equal to its mean; in App. B.2, we show that, under mild conditions, a true Poisson noise model (with no noise correlations) leads to the same results as we derive below for the Gaussian noise model, Eq. (6), with *σ* = 2*α* = 1 and Σ(***s***) = *I*.

In addition to the gains, the objective ℒ (***g***), Eq. (4), also depends tacitly on (*i*.*e*., is a functional of) the stimulus distribution *P*, the representational curves **Ω**, and the stimulus-dependent noise correlation matrix Σ. As shown in App. B.1, for our class of noise models, Eq. (6), ℒ depends on *P*, **Ω**, and Σ only through the following collection of signal and noise statistics. (By “signal” we refer to the noise-free trial-averaged population response, ***h***(***s***); under variations of stimulus, ***h***(***s***) can be thought of as a multivariate random variable with associated statistics.)

1. The units’ normalised average responses

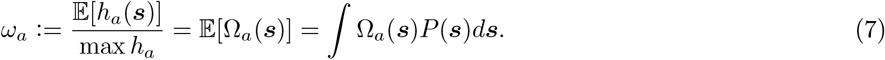

These are related to the units’ mean responses, *r*_*a*_, via *r*_*a*_ := 𝔼[*n*_*a*_] = 𝔼[*h*_*a*_(***s***)] = *g*_*a*_*ω*_*a*_.
2. The relative size of stimulus-dependent signal variations, as quantified by the coefficients of variation of the units’ trial-average responses, CV_*a*_ := SD[*h*_*a*_(***s***)]*/*𝔼[*h*_*a*_(***s***)].
3. Signal correlation structure, as quantified by the matrix, *ρ*, of correlation coefficients of the trial-averaged responses, *ρ*_*ab*_ := Cov[*h*_*a*_(***s***), *h*_*b*_(***s***)] */* (SD[*h*_*a*_(***s***)]SD[*h*_*a*_(***s***)]).
4. Noise correlation structure, as captured by the average normalised noise covariance matrix^3^, *W*, defined by

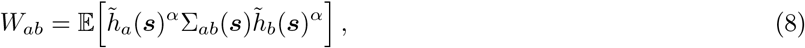

where 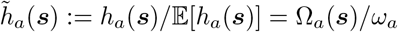 denotes the tuning curve normalised by its stimulus average.

Note that all these statistics are invariant with respect to arbitrary rescalings of the tuning curves, and are thus independent of the gains — in particular, in all the definitions above, the *h*_*a*_ could be replaced with the Ω_*a*_. In terms of these quantities, the objective function, Eq. (4), is given by

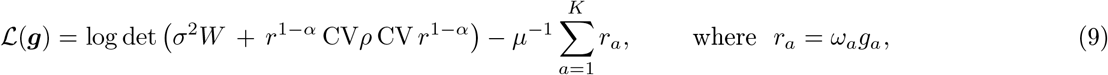

and — in line with the convention introduced above — CV and *r* denote *K × K* diagonal matrices with CV_*a*_ and *r*_*a*_ = *ω*_*a*_*g*_*a*_ on their diagonals, respectively. Thus the objective depends on the gains, ***g***, only via the average responses ***r***. In addition, it also depends on the gain-independent quantities, CV, *ρ*, and *W*, which capture aspects of signal and noise statistics.

We now present the results of numerical optimisation of the objective Eq. (9) for different noise distributions within the general family of noise models, Eq. (6), for a population of *K* = 10, 000 units. To capture the notion of adaptation to changes in the environment or context, we simulated a family of environment models smoothly parameterised by a single parameter *ν* ∈ [0, 1]. As we have just seen, the objective depends only on *ω, ρ, W*, and CV. Changes in these quantities can arise due to changes in the stimulus density, *P* (***s***), the representational curves of the population, Ω(***s***), or both. We specify environment models only through the parameters on which the objective depends (*ω, ρ, W*, and CV), remaining agnostic to the underlying cause of changes to these quantities.

For simplicity, we fixed the response coefficient of variation CV_*a*_ = 1, the noise scale *σ*^2^ = 1, and the information-energy trade-off parameter *µ* = 5*/*(1 − *α*) in all environments. (Below, in Sec. 2.5, we will discuss the biological basis for these parameter choices.) On the other hand, the normalised average response, *ω*_*a*_, and the signal correlation matrix, *ρ*, depended smoothly on the environment parameter *ν*, as follows. For each unit *a* = 1, …, *K*, the normalised average responses in the extreme environments, *ω*_*a*_(*ν* = 0) and *ω*_*a*_(*ν* = 1), were drawn independently from a wide distribution (specifically the beta distribution Beta(6, 1)). The normalised average response at intermediate values of *ν, ω*_*a*_(*ν*), was then obtained by linear interpolation between the independently sampled boundary values. This method for sampling environments is illustrated in Fig. 4. Similarly, to construct the signal correlation matrix *ρ*(*ν*), we first constructed the boundary values *ρ*(*ν* = 0) and *ρ*(*ν* = 1) with independent random structures and smoothly interpolated between them (see Sec. 4.1 for the details). We did this in such a way that, for all *ν*, the correlation matrices, *ρ*(*ν*), were derived from covariance matrices with a 1*/n* power-law eigenspectrum (*i*.*e*., the ranked eigenvalues of the covariance matrix fall off inversely with their rank), in line with the findings of Stringer et al. (2019) in the primary visual cortex. Finally, the stimulus-averaged noise correlation matrix, *W*, is determined by our choice of the noise model subfamily used in each simulation. We now present simulation results for three 1-parameter subfamilies of the general family of noise models, Eq. (6), in turn.

**Figure 4:**
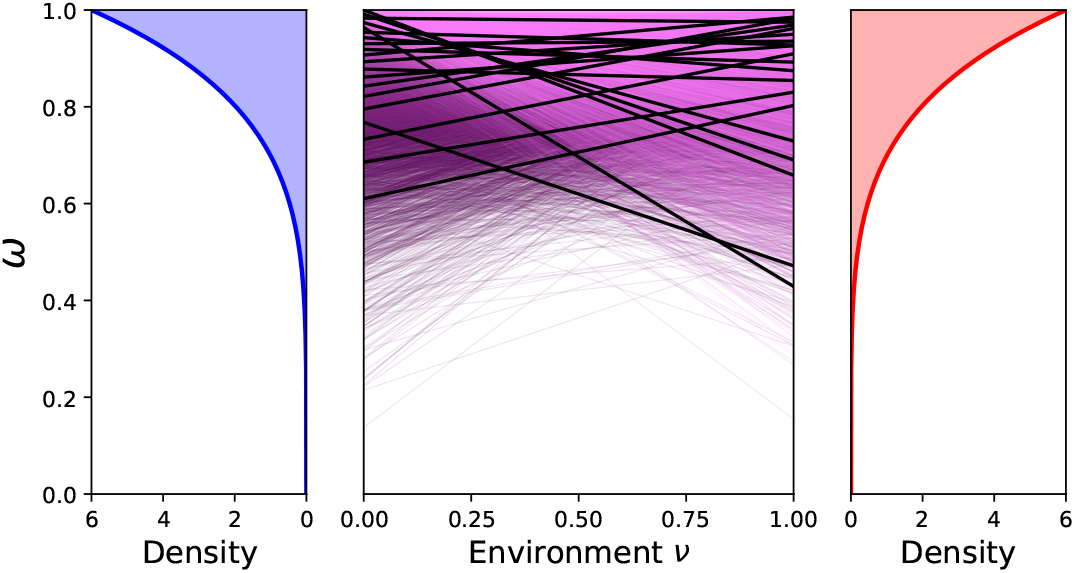
The structure of pre-modulated average population responses in the simulated environments. We simulated a one-parameter family of environments parameterised by *ν* ∈ [0, 1]. In the environments on the two extremes, for each unit, the gain-independent average normalised responses *ω*_*a*_(*ν* = 0) and *ω*_*a*_(*ν* = 1) are randomly and independently sampled from a Beta(1, 6) distribution, with density shown on the left and right panels. For intermediate environments (*i*.*e*., for 0 *< ν <* 1), *ω*_*a*_(*ν*) is then obtained by linearly interpolating these values (middle panel). The middle panel shows *K* = 10000 such interpolations, with every 500th interpolation (ordered by initial value) coloured black.

#### Uncorrelated power-law noise

Consider first the case of no average noise correlations, in the sense that *W* = *I*.^4^ We investigated the adaptation behavior of optimally gain-modulated population responses as a function of the noise scaling exponent *α*. Fig. 5A shows the scaling of spike count variance as a function of mean spike count for different values of the noise scaling exponent *α*. By construction, in the absence of adaptation, an environmental shift (*ν* = 0 *→ ν* = 1) results in a wide range of average firing rates (Fig. 5B, middle panel). But following adaptation, the firing rates become tightly concentrated around a single value (Fig. 5B, bottom panel), which matches the value they were concentrated around in the original environment (Fig. 5B, top panel). Thus, adaptation results in (1) *uniformisation* of average firing rates across the population in a fixed environment, and (2) firing rate *homeostasis*, where following adaptation the mean firing rates of the units returns to their value before the environmental shift. These same uniformisation and homeostatic effects are seen for other values of *α* (Fig. 5B and Supp. Fig. 14). In Fig. 5C we quantify deviation from perfect homeostasis as the average % difference in the units’ mean firing rates between the initial environment (*ν* = 0) and environments at increasing *ν*. As shown, the faster the spike count variance grows with mean spike count (*i*.*e*., the greater the noise scaling exponent *α* is) the greater the deviation from firing rate homeostasis. But even for super-poissonian noise with *α* = 0.75 the average relative change in firing rates between the environments never exceeds 3% (Fig. 5C).

**Figure 5:**
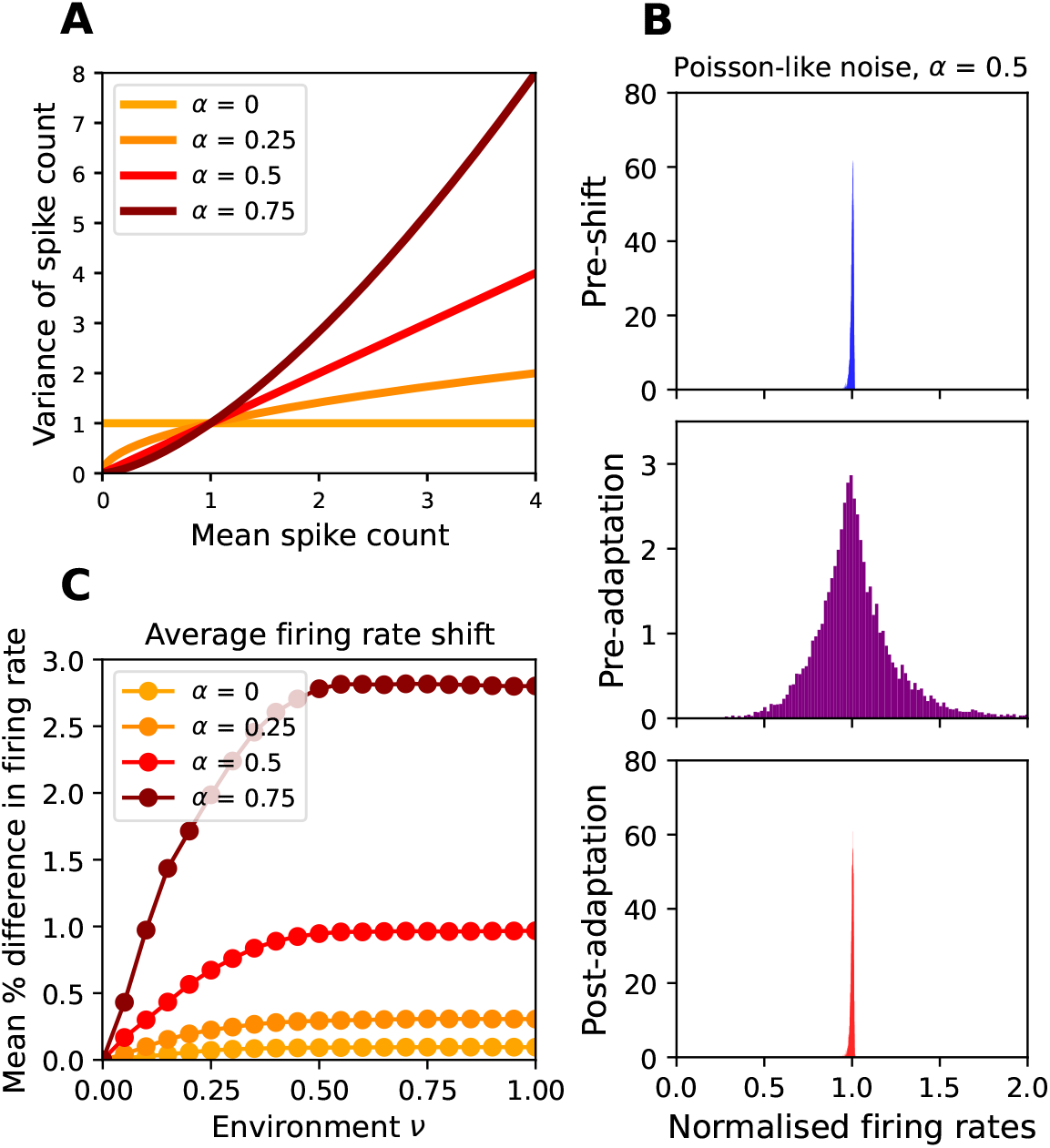
Optimal gains under uncorrelated power-law noise lead to homeostasis of rates. (A) Illustration of power-law noise. The trial-to-trial variance of spike counts, as a function of the trial-averaged spike count, is shown for different values of the noise scaling parameter *α*. (B) Distribution of average firing rates before and after a discrete environmental shift from *ν* = 0 to *ν* = 1 for the case *α* = 1*/*2. Top: the histogram of firing rates adapted to the original environment (*ν* = 0) before the shift, as determined by the optimal gains Eq. (5) in this environment, 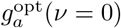. Middle: the histogram of firing rates immediately following the environmental shift to *ν* = 1, but before gain adaptation; these are proportional to 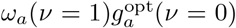. Bottom: the histogram of firing rates after adaptation to the new environment, proportional to 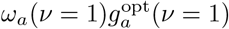. We have normalised firing rates such that the pre-adaptation firing rate distribution has mean 1. (C) Deviation from firing rate homeostasis for different values of *α* as a function of environmental shift. For each environment (parametrised by *ν*), we compute the optimal gains and find the adapted firing rate under these gains. The average relative deviation of the (post-adaptation) firing rates from their value before the environmental shift, 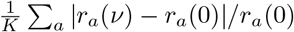, is plotted as a function of *ν*.

Even under super-poissonian noise (*α* = 0.75 in Fig. 5C), the average relative change in firing rates between environments never gets above 3%. Fig. 5C additionally shows that the slower the spike count variance grows with mean spike count (*i*.*e*., the smaller the noise scaling exponent *α* is) the more precise the firing rate homeostasis.

#### Uniform noise correlations

In the second sub-family of noise models we allow for noise correlations, but consider only Poisson-like scaling of noise (*i*.*e*., a noise scaling exponent *α* = 1*/*2). More precisely, we assume that the effective noise correlation coefficients between all pairs of units (corresponding to off-diagonal elements of the matrix *W*, Eq. (8)) are the same, equal to *p* (Fig. 6A).^5^ As shown in Fig. 6B, optimal gain adaptation leads to uniformisation and homeostasis of mean firing rates in this noise model as well. In fact, in the presence of uniform noise correlations, homeostasis is even stronger compared to the previous noise model with zero noise correlations; in this case, mean relative deviations in average firing rates never exceed 1% (Fig. 6C). Further, as noise correlations get stronger (*i*.*e*., as *p* increases) we see tighter homeostasis (see also Supp. Fig. 15).

**Figure 6:**
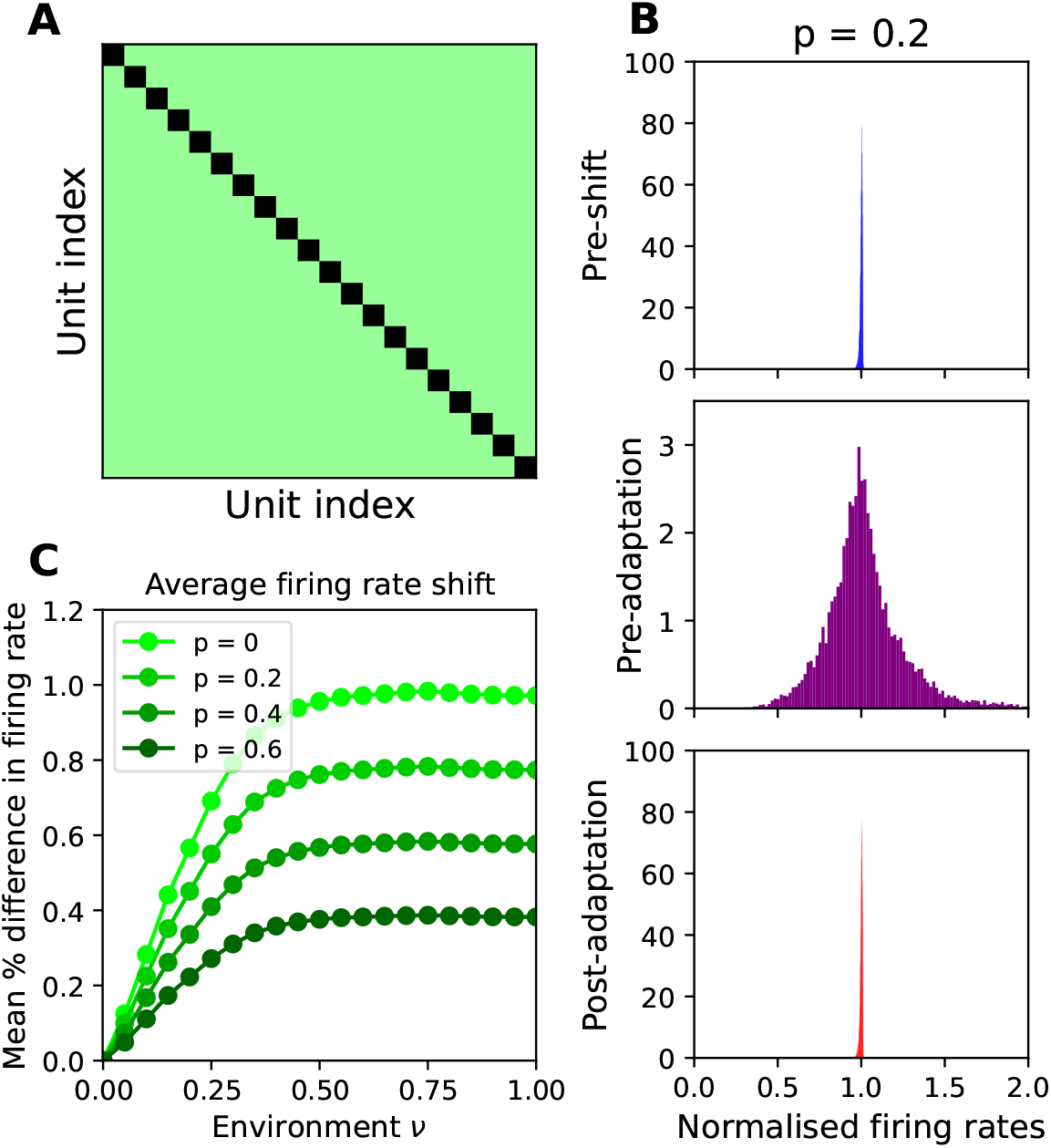
Firing rate homeostasis under Poissonian noise with uniform noise correlations. (A) Illustration of an effective noise correlation matrix with uniform correlation coefficients; the (green) off-diagonal elements have the same value, *p <* 1, while the (black) diagonal elements are 1. Panels B and C have the same format as in Fig. 5.

#### Aligned correlated noise

In the final noise model we considered, we once again fixed *α* = 1*/*2 (Poisson-like scaling), but allowed for heterogeneous effective noise correlation coefficients. Specifically, we took the effective noise correlation matrix *W* to have approximately the same eigenbasis as *ρ*, the signal correlation matrix — hence “aligned noise”—, but with a different eigenvalue spectrum. As described above, for all noise sub-families, the signal correlation structure *ρ*(*ν*) was obtained by normalising a covariance matrix with 1*/n* spectrum (corresponding to the findings of Stringer et al. (2019) in V1); in the current case, we obtained the noise correlation structure *W* (*ν*) by normalising a covariance matrix with the same eigenbasis but with a 1*/n*^*γ*^ spectrum (*i*.*e*., with the *n*-th largest eigenvalue of *W* scaling as 1*/n*^*γ*^); see Fig. 7A, and Sec. 4.1 for further details. Fig. 7B demonstrates that, as with the other noise sub-families, optimal gain adaptation in the presence of aligned correlated noise also leads to homeostasis and uniformisation of mean firing rates (see Supp. Fig. 16 for rate histograms for other values of *γ*). Across different values of *γ*, homeostasis is again tighter compared to the zero noise correlation case, with the average firing rate shift never exceeding 1% (Fig. 5C). Further, with the exception of *γ* = 1, we see more homeostasis for higher values of *γ*, corresponding to lower dimensional noise. The special case of *γ* = 1 corresponds to so-called information-limiting noise (Moreno-Bote et al., 2014; Kanitscheider et al., 2015), where noise and signal correlation structures are fully aligned: *W* = *ρ*. In this case, homeostasis is perfect, and the mean relative error in firing rates is 0. We provide an analytic proof that in this case we have perfect homeostasis and uniformisation in App. B.5, and discuss this situation further in Sec. 2.5 and Sec. 2.6.

**Figure 7:**
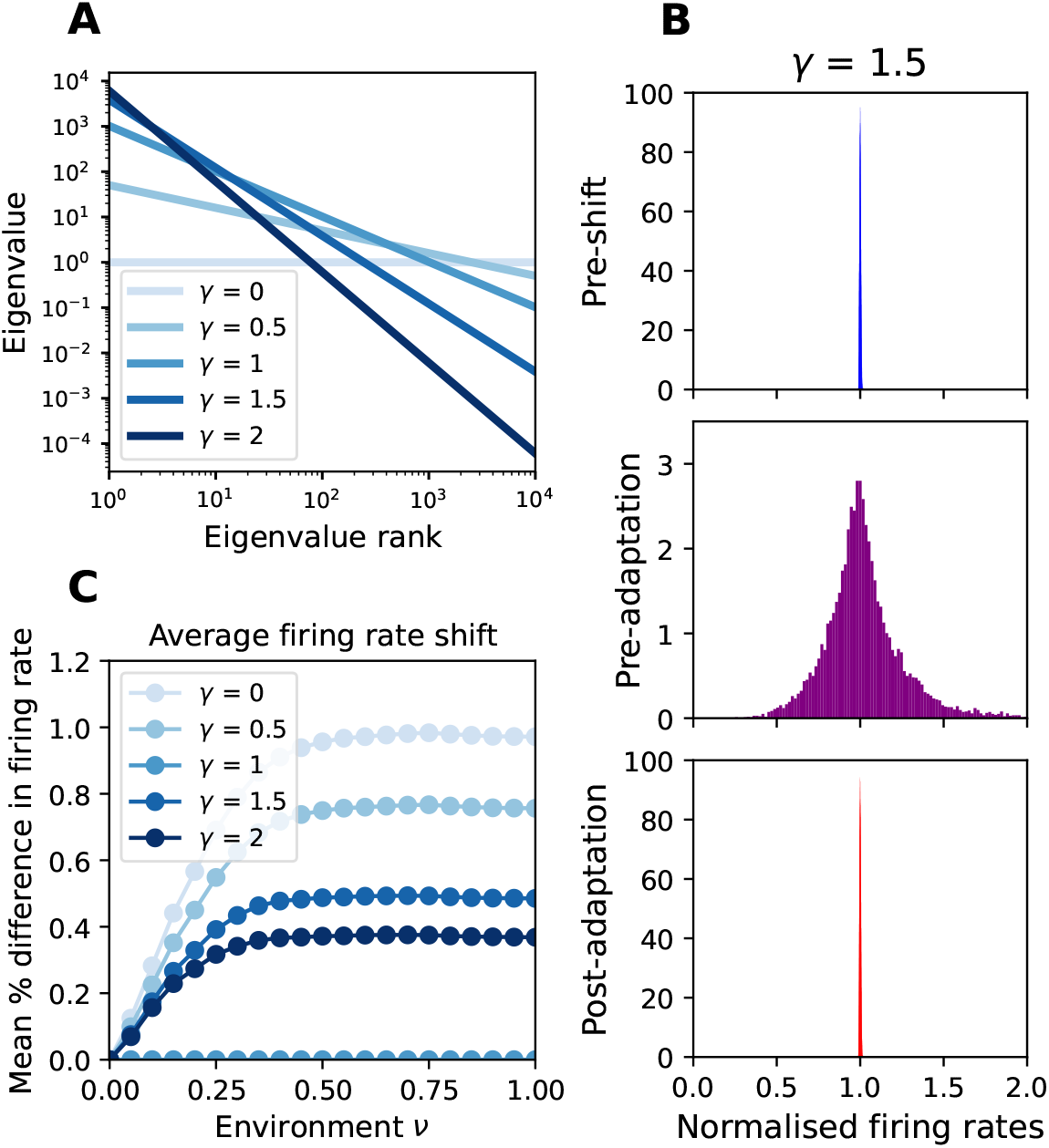
Firing rate homeostasis under signal-aligned noise. (A) Eigenspectra of noise covariance matrix for the aligned noise model for various values of *γ*; in this model, the *n*-th eigenvalue of the average noise correlation matrix, *W*, is proportional to 1*/n*^*γ*^. Panels B and C have the same format as in Fig. 5.

We have shown that, according to our efficient coding framework, optimal adaptation of response gains (with the goal of combating neural noise at minimal metabolic cost) can robustly account for the firing rate homeostasis of individual units within a population, despite environmental shifts or changes in neural representations (tuning curve shapes), under a diverse family of noise models. To shed light on these results, in Sec. 2.4, we analytically investigate sufficient conditions on the parameters, Eq. (7)–(8), of our model under which we expect homeostasis to emerge. Following that, in Sec. 2.5, we will show that biological estimates for these parameters obtained in cortex (which are determined by natural stimulus statistics, as well as the stimulus dependence of high-dimensional population responses) are consistent with these conditions.

### 2.3 A clustered population accounts for the diversity of firing rates seen in cortex

In addition to homeostasis, the above simulation results also predict uniformisation of average firing rates across the population, in any given environment. By contrast, cortical neurons display a wide range of average firing rates spanning a few orders of magnitude (Buzsáki and Mizuseki, 2014). But a cortical neuron can have very similar response tuning to other neurons in the same area, and neurons can therefore be grouped based on the similarity of their tuning curve shapes. We can thus interpret the units in our preceding numerical experiments not as single neurons, but as clusters of similarly tuned neurons. In this interpretation, uniformisation occurs at the cluster level, with potentially a wide distribution of single-cell average firing rates within each cluster. We found that our model is indeed consistent with such an interpretation. In particular, in the case of uncorrelated noise with Poisson-like mean-variance scaling, we show that a simple extension of our model gives rise to both diversity of firing rates as seen in cortex, as well as homeostasis and uniformisation at the level of clusters of similarly tuned neurons.

To gain intuition and allow for analytic solutions, we study a toy model in which the *N* neurons in the population are sorted into *K* such clusters. In this toy model, neurons within a cluster have very similar stimulus tuning, but neurons in different clusters are tuned differently (Fig. 8A); the model thus exaggerates a more realistic situation in which there is a more gradual transition from similar to dissimilar tuning curves within the entire population. A perturbation strength parameter, *ϵ ≥* 0, controls how much the tuning curve shapes of single neurons in a cluster deviate from the stereotype tuning curve shape for that cluster (see Eq. (25) in Methods). When *ϵ* = 0, we have perfect clusters, *i*.*e*., neurons in a cluster have identical representational curves. As the perturbation strength *ϵ* increases, within-cluster deviations become stronger.

**Figure 8:**
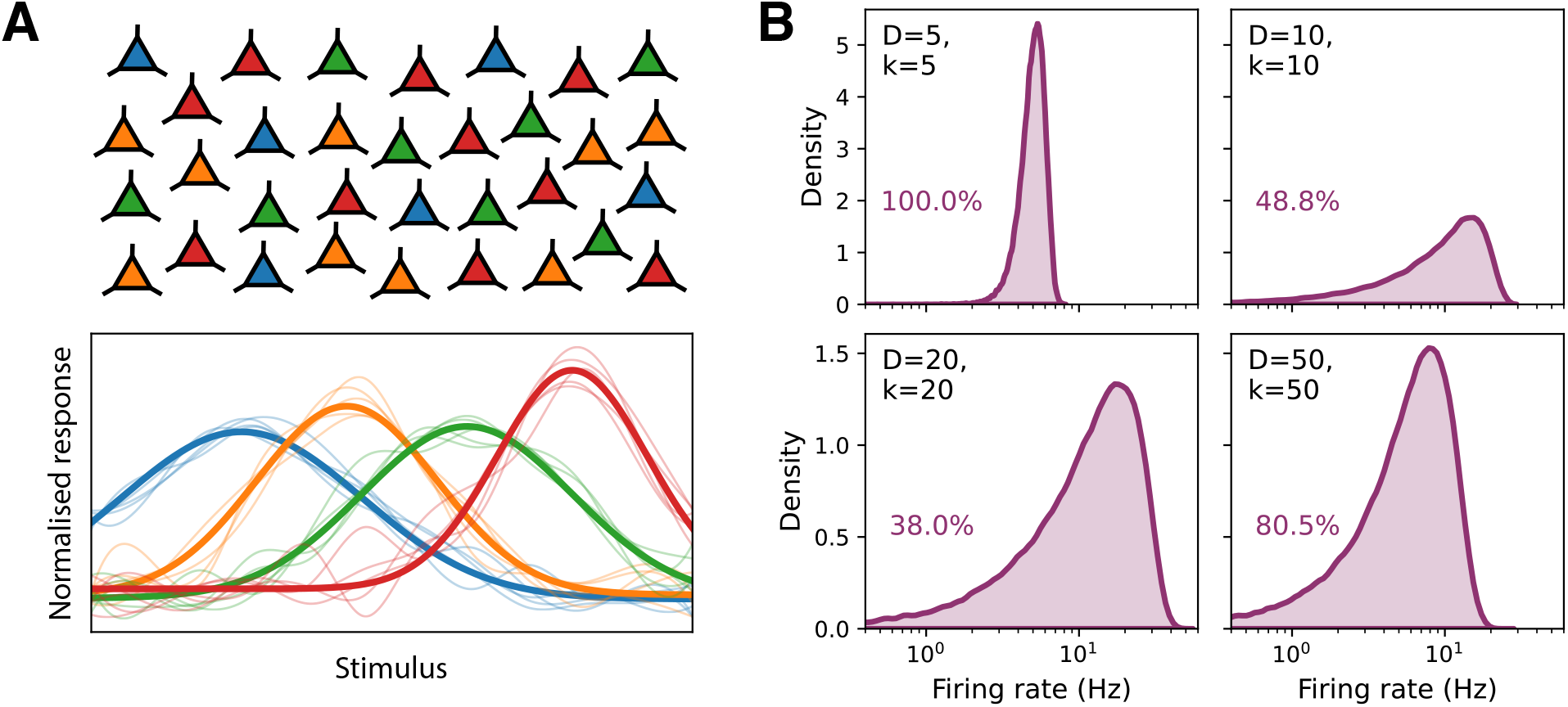
Optimal gain adaptation in a clustered population accounts for the broad distribution of cortical single-neuron firing rates. (A) A cartoon description of the clustered neural population model. Top: The single neurons in the population belong to different functional groups or clusters (shown by different colors). Bottom: The normalised tuning curves of the single neurons belonging to the same cluster (thin lines of the same color) are very similar, and close to their cluster’s stereotype normalised tuning curve (solid lines) which can be very different across clusters. (B) Each panel shows the distribution of log firing rates of the active single neurons for different choices of cluster size *k* and effective dimensionality, *D*, of the within-cluster tuning curve variations. The distributions shown are for the active neurons with non-zero firing rate, with their percentage of the total population given in each panel (purple text). For all simulations we fixed the cluster firing rate to *k ×* 5 Hz (corresponding to *µ* = 10, if we assume a coding interval of 0.1 s), such that the mean rate of single neurons is 5 Hz on average.

We assume single-neuron gains adapt to optimise the same efficient coding objective defined above in Eq. (9), but with units now clearly interpreted as single neurons. Note that optimisation in terms of the neural gains is equivalent to optimisation in terms of the neurons’ mean firing rates (because the objective depends on the neurons’ gains only via their mean fire rates each neuron’s mean rate is proportional to its gain by a fixed constant of proportionality, given by the mean value of its fixed and unoptimised representational curve, Eq. (7)). For *ϵ* = 0 (identical tuning curve shapes within each cluster), the efficient coding objective function is blind to the precise distribution of single-cell firing rates among the neurons of a cluster, as long as the total rate of the cluster is held fixed. This is because for clusters of identically-tuned and independently-firing neurons with Poisson-like noise, the cluster can be considered as a single coherent unit whose firing rate is given by the sum of the firing rates of its neurons. Thus at zeroth order, the distribution of individual neuron rates are completely free and undetermined by the optimisation, as long as they give rise to optimal total cluster rates.

For small but non-zero perturbation strength *ϵ* (corresponding to non-identical but sufficiently similar within-cluster tuning curve shapes), we showed that the efficient coding optimisation breaks down into two optimisation problems: one at the level of clusters, wherein the mean cluster rates are optimised, and one at the level of individual neurons within each cluster (Methods Sec. 4.2 and App. B.3). The cluster-level problem remains decoupled from the within-cluster distribution of single-neuron rates; thus the optimisation of cluster firing rates reduces to the same optimisation problem considered in the previous sections, Eqs. (4)–(5), with the units now interpreted not as neurons but as clusters of similarly tuned neurons. On the other hand, within-cluster optimisation of individual neuron rates decouples across clusters. Within each cluster, we show that this optimisation is approximately given by a quadratic program, subject to the constraint that the individual neuron rates sum up to the optimal total cluster rate found by solving the cluster-level problem (Eqs. (28)–(30) in Sec. 4.2).

The within-cluster quadratic program optimisation is specified by the signal covariance structure of the normalised tuning curves in the cluster, or more precisely, the covariance matrix of their deviations from the cluster stereotype normalised tuning curve (see Eq. (30)). A key characteristic of any covariance matrix is its effective dimensionality (corresponding roughly to the number of principal components with significant variance). Instead of making arbitrary choices about the signal covariance matrix defining the within-cluster optimisation, in our simulations we sampled them from a simple random ensemble which was controlled only by an effective dimensionality parameter, *D*, and the cluster size, *k* (see Sec. 4.3 for details). Figure 8 (see also Supp. Fig. 18) shows the resulting firing rate distributions for neurons aggregated across all clusters (corresponding to optimisations with independently sampled covariance matrices), for different values of *k* and *D*. Firstly, note that, depending on these parameters, a significant fraction of neurons can be silent, *i*.*e*., with zero firing rate in the optimal solution; the figure shows the percentage of and the rate distributions for the active neurons. Our theory therefore predicts that a significant fraction of cortical neurons are silent in any given sensory environment, and that which neurons are silent can shuffle following shifts in environmental stimulus statistics (specifically shifts that result in significant changes in the signal correlation structure of the cluster neurons). Secondly, note that the distributions span multiple orders of magnitude, in agreement with empirical observations in the cortex (see *e*.*g*. Slomowitz et al. (2015); Hengen et al. (2013); Maffei and Turrigiano (2008)), and is approximately log-normal (albeit with a skewed tail towards small log-rates), also consistent with empirical findings (Buzsáki and Mizuseki, 2014). Lastly, for fixed *k*, as the effective tuning curve dimension, *D*, increases, the fraction of silent neurons decreases (Supp. Fig. 18).

To summarise, we have shown that the objective ℒ considered in Sec. 2.2, reinterpreted to correspond to the cluster level, can arise from optimisation of a similar objective, defined at the level of single neurons in a functionally clustered population (Eqs. (26)–(27)). Furthermore, the correction to the cluster-level objective (arising from within-cluster variations in neural tuning) serves to break the symmetry within clusters, generating a diverse range of firing rates which matches the distribution of cortical single-neuron firing rates. In the rest of the paper, we will pursue the problem at the coarser level of clusters.

### 2.4 Analytical insight on the optimality of homeostasis

The numerical results of Sec. 2.2 showed that, across a range of neural noise models, optimal gain modulation for combating noise can give rise to firing rate homeostasis and uniformisation across neural clusters defined based on similarity of stimulus tuning. To shed light on these results, we analytically investigated how and in what parameter regimes these results arise. We found that homeostasis and uniformisation emerge in the regime where the matrix

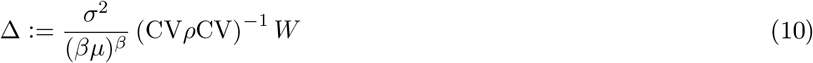

is small (according to an appropriate matrix norm); here *β* = 2 − 2*α*. The intuitive meaning of this matrix is that its inverse, Δ^−1^, characterises the strength of the signal-to-noise ratio (SNR) along different directions in the code space, *i*.*e*., the space of cluster responses. Therefore the above condition can be roughly interpreted as the requirement that signal-to-noise ratio is strong.^6^ We will discuss this condition, and the constraints it imposes on various model parameters, further in Sec. 2.5, and will provide evidence that it is likely to hold in cortex. Here, we focus on its consequences.

We developed a perturbative expansion in Δ, to obtain approximate analytical solutions for the cluster gains that maximise our objective ℒ, Eq. (9) (see App. B.4). In the leading approximation, *i*.*e*., to zeroth order in Δ, this expansion yields

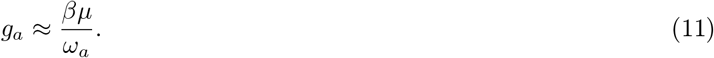

We denote this approximate solution by 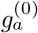. To see the significance of this result, note that the average response of cluster *a, i*.*e*., its stimulus-averaged total spike count is given by 𝔼[*h*_*a*_(***s***)] = *g*_*a*_*ω*_*a*_. Thus, in the leading approximation, the stimulus-average spike count of cluster *a* is given by the constant 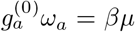. Thus, since *βµ* is constant across both clusters and environments, the zeroth order solution yields homeostasis and uniformisation of cluster firing rates.

In Sec. 2.6, we numerically quantify the performance of the homeostatic approximation, ***g***^(0)^, in terms of maximising the efficient coding objective, under different noise models. Here, we also note the following exact analytical result regarding homeostasis that holds even for non-small Δ. It follows form the structure of the objective function, Eq. (9), that, irrespective of how pre-modulation mean rates, *ω*_*a*_, change between two stimulus environments, as long as Δ and *W*, the effective noise covariance, remain invariant across the environments, optimal gains imply exact homeostasis of each cluster’s average firing rate across the two environments. However, even in this case, unless Δ is small, in general the optimal solution does not result in uniformisation of firing rates across clusters.

### 2.5 Conditions for the validity of the homeostatic solution hold in cortex

We asked whether cortical populations satisfy the conditions under which the homeostatic solution, Eq. (11), is expected to provide a good approximation to optimal gains according to efficient coding. As noted above, the general mathematical condition is that the matrix Δ, defined in Eq. (10), is in some sense small (recall that through its inverse, Δ represents the high-dimensional structure of signal-to-noise ratio in the coding space, under the homeostatic solution Eq. (11)). We can make this statement more precise by looking at the size of the leading-order correction to the approximate homeostatic solution Eq. (11), in the perturbative expansion. We found that the first-order correction modifies the approximate homeostatic solution for the gain *g*_*a*_ by a multiplicative factor 1 − Δ_*aa*_ (see Eq. (106) in App. B.4). In other words, to leading order, the relative error in the homeostatic approximation to the optimal gain (and thus the optimal mean response) for unit *a* is given by |Δ_*aa*_|; we can thus reformulate the condition for the accuracy of the homeostatic approximation as

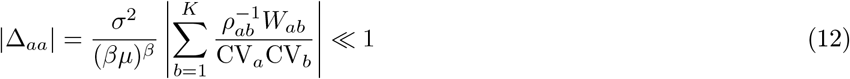

for all *a*. Conversely, in the first approximation, deviations from homeostasis depend on the SNR structure, as captured by Δ_*aa*_, and how that structure shifts between environments.

Before proceeding to empirical estimates of Δ_*aa*_, let us first delineate the dependence of Δ on different model parameters. Firstly, note that Δ scales linearly with the noise strength *σ*^2^; thus, if noise strength is scaled up, so is Δ. Next, consider increasing the noise scaling exponent *α*. This decreases (*βµ*)^*β*^ provided *α* 1 *≤* − *e/*2*µ* (which holds in our simulations above). Moreover, increasing *α* increases the diagonal elements of *W* (see Eq. (8)). Intuitively, this increases the magnitude of *W*. Through these two mechanisms, increasing the noise scaling exponent *α* will generally increase the size of Δ. Next, note that Δ has an inverse relationship with the coefficients of variation, CV, which increase as neurons become more stimulus selective; accordingly, the more selective the tuning curves, the smaller the Δ. Lastly, Δ has a complex dependency on the spectra of both CV*ρ* CV and the noise correlation structure *W* as well as the alignment of their eigenbases (which represent the geometric structure of signal and noise in the neural coding space, respectively). In Sec. 2.6, we will numerically investigate these dependencies, and their effects on the validity of the homeostatic approximation.

Translated to biological quantities, it follows from the above dependencies that the smallness of Δ (or Δ_*aa*_) requires that (1) the cluster firing rates are sufficiently high, (2) noise variance scales sufficiently slowly with mean response, (3) individual clusters are sufficiently selective in responding to stimuli, and (4) the neural representation has a high-dimensional geometry in the population response space. We used the available biological data to estimate the size of these variables for cortical populations; see Sec. 4.4. Here we summarise the resulting estimates.

#### Sufficiently high firing rate

In the homeostatic approximation, Eq. (11), *βµ* is equal to the average total spike count of a cluster over the rate-coding time interval. Condition Eq. (12) thus requires that the stimulus-averaged firing rate of all clusters are sufficiently high. Based on the available data (see Sec. 4.4), we estimate the mean firing rate in rodent V1 to be around 5 Hz. Assuming a coding interval of 0.1 seconds, and a cluster size of *k* = 20, this yields *βµ ~* 10.

#### Sufficiently slow scaling of noise variance

We can obtain an estimate for the prefactor *σ*^2^*/*(*βµ*)^*β*^, by fitting the following mean-variance relationship for cluster spike counts: variance = *σ*^2^mean^2−*β*^. Empirical estimates of this relationship lead to estimates of *σ*^2^*/*(*βµ*)^*β*^ ranging from 0.05 (Shadlen and Newsome, 1998) to 0.31 (Koyama, 2015), with a median value 0.14 (see Sec. 4.4 for details). This is close to the value for Poissonian noise (*σ*^2^ = 1, *β* = 1), which is 0.1.

#### Sufficiently selective responses

The coefficient of variation CV for neural responses is a measure of neural selectivity. Accordingly, it is related to another measure of stimulus selectivity, namely the coding level (*i*.*e*., the fraction of stimuli to which a neuron responds). Using empirical estimates of the latter quantity for cortical neurons (Lennie, 2003), we estimated CV^2^ *≈* 20. Below we take CV^2^ *≈* 10 as a more conservative estimate.

#### High-dimensional signal geometry

Assuming, for simplicity, that the coefficients of variation are approximately the same for all clusters, we see from Eq. (12) that Δ_*aa*_ are proportional to the diagonal elements of the matrix *ρ*^−1^*W*. Assuming further, that these elements are all comparable in size,^7^ we estimated their order of magnitude based on the known structure of the signal correlation matrix, *ρ*, in V1 (see below), and for the aligned noise family of average noise covariance matrices, *W*, studied in Sec. 2.2. This family includes two biologically relevant cases: (1) an uncorrelated noise model corresponding to *W* = *I*, and (2) a *W* with so-called information-limiting noise correlations (Moreno-Bote et al., 2014; Kanitscheider et al., 2015). In the first case Δ_*aa*_ is proportional to [*ρ*^−1^]_*aa*_. It can be shown that [*ρ*^−1^]_*aa*_ *≥* 1, with equality if and only if cluster *a* has zero signal signal correlation with every other cluster. [*ρ*^−1^]_*aa*_ can thus be seen as a measure of representational redundancy. Many traditional efficient coding accounts predict zero signal correlation between neurons, at least in the low noise limit (Barlow, 2012; Atick and Redlich, 1990; Nadal and Parga, 1994, 1999), providing a normative justification for low signal correlations. However, complete lack of signal correlations is not necessary for [*ρ*^−1^]_*aa*_ to be small. Geometrically, small signal correlations correspond to neural responses forming a representation with high intrinsic dimensionality. Stringer *et al*. (Stringer et al., 2019) found that the signal covariance matrix of mouse V1 neurons responding to natural stimuli possesses an approximately 1*/n* eigen-spectrum. In this case, for *K* large, we obtain the estimate [*ρ*^−1^]_*aa*_ *≈* ln(*K*)*/*2 (see App. B.6). Given the slow logarithmic increase with *K*, we expect that our condition can hold even for very large neural populations. For example, suppose we take the entire human V1 as our neural population, to obtain an arguably generous upper bound for *K*. This contains roughly 1.5 *×* 10^8^ neurons (Wandell, 1995), leading to 7.5 *×* 10^6^ clusters of 20 neurons, and therefore an average value of [*ρ*^−1^]_*aa*_ just under 8. This, together with our other estimates above, yields the following overall estimate for Δ_*aa*_

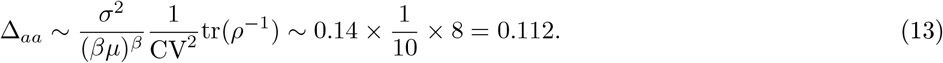

We found that in the case of *information-limiting noise correlations* the |Δ_*aa*_| tend to be even smaller than the above estimate. Information-limiting noise correlations refer to the case where directions of strong noise in the coding space align with those of strong signal variation. The extreme case, *i*.*e*., perfect alignment of noise and signal geometry, corresponds to *ρ* = *W*, or equivalently *ρ*^−1^*W* = *I*, for which Δ_*aa*_ *∝* [*ρ*^−1^*W*]_*aa*_ = 1 does not grow with *K* at all. In App. B.6 we analysed less extreme forms of alignment (corresponding to the model numerically studied in Fig. 7, Sec. 2.2), and found (unsurprisingly) that for those cases, the estimates of Δ_*aa*_ lie between the estimates for the two above cases. Accordingly, even partial signal-noise alignment can (potentially strongly) reduce the size of Δ below the estimate Eq. (13), further encouraging homeostatic coding.

The above analysis makes it clear when we should expect homeostasis in general – when cluster firing rates are high enough, the noise scaling of cluster responses is sufficiently slow, responses are highly selective, and signal correlation structure corresponds to a high-dimensional geometry (Stringer et al., 2019), possibly with information-limiting noise correlations (Moreno-Bote et al., 2014; Kanitscheider et al., 2015). On the other hand, when these conditions are violated, the optimal gain configuration can potentially deviate strongly from the homeostatic solution. Moreover, from the above estimates, leading to Eq. (13), we see that in V1 and possibly other cortical areas, we can expect the leading corrections to the homeostatic solution to not exceed 10%, in relative terms.

### 2.6 Numerical comparison of homeostatic gains to optimal gains

We now return to the numerical simulations performed in Sec. 2.2, to evaluate the performance and accuracy of the approximate homeostatic solution, 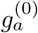, Eq. (11), and the first-order correction to it, 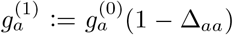, relative to the true optimal solution of Eq. (9). We also evaluate the performance and accuracy of the best possible homeostatic solution, *i*.*e*., the best configuration of cluster gains, 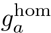, that, by construction, are guaranteed to yield homeostasis and uniformisation. In other words, under such gains the mean responses *r*_*a*_ = *g*_*a*_*ω*_*a*_ must equal a constant (*i*.*e*., independent of *a*, the cluster index, and constant across environments). Denoting this *a priori* unknown constant by *χ*, we thus have

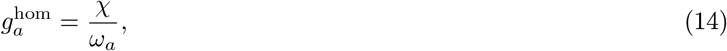

under which the average spike count of clusters in the coding interval is given by *χ*. The variable *χ* is then chosen to optimise the expectation of ℒ, Eq. (9), across a relevant family of environments (note that we need to optimise the average objective across environments, since otherwise the optimal *χ* will in general depend on the environment, invalidating homeostasis). With such an optimal *χ*, ***g***^hom^ will, by definition, perform better than ***g***^(0)^ in terms of our objective function, ℒ.^8^ In fact, as we will show below, in our simulations (with model parameters within the biologically plausible ranges identified in the previous section), the homeostatic ***g***^hom^ performed very close to the true optimal gains.

For each of the different noise models considered in Sec. 2.2 we compare the performance of the homeostatic gains ***g***^(0)^ and ***g***^hom^, as well as the perturbative correction, ***g***^(1)^, to the numerically optimised gains. To examine the adaptation properties of these gains, we once again work with the sequence of environments indexed by *ν* ∈ [0, 1], with the same specifications as in the numerical simulations of Sec. 2.2. To measure the accuracy of the approximations to the optimal gains, for each environment *ν*, we calculated the mean relative errors

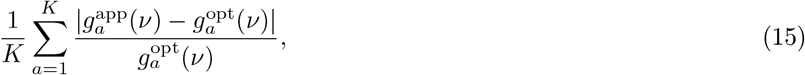

for each of the approximate solutions, denoted by ***g***^app^(*ν*), with ***g***^opt^(*ν*), the numerically optimised solution in environment *ν*. To measure the performance of an approximate solution, we use the following *relative improvement* measure, defined by

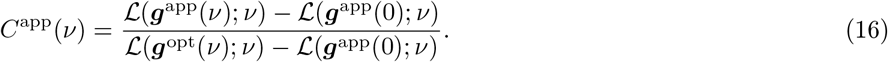

which can be interpreted as the improvement in the objective ℒ (*·*; *ν*) (the efficient coding objective Eq. (9) according to the statistics of environment *ν*) achieved by the adaptive gains ***g***^app^(*ν*) over the unadapted gains from the original environment ***g***^app^(0), relative to the improvement obtained by the optimally adaptive gains ***g***^opt^(*ν*). The better the approximation the closer *C*^app^(*ν*) will be to 1. Note that, in general, the relative improvement worsens when *ν* goes to 0 (see Fig. 9B, D and F); this follows from the definition of *C*(*ν*) and the fact that when environmental change is weak, using gains optimised in the original environment without adapting them in the new environment is superior to adaptive but only approximately optimal gains. We now summarise the results for the different noise models discussed in Sec. 2.2 (see that section for the detailed specification of the three noise models).

**Figure 9:**
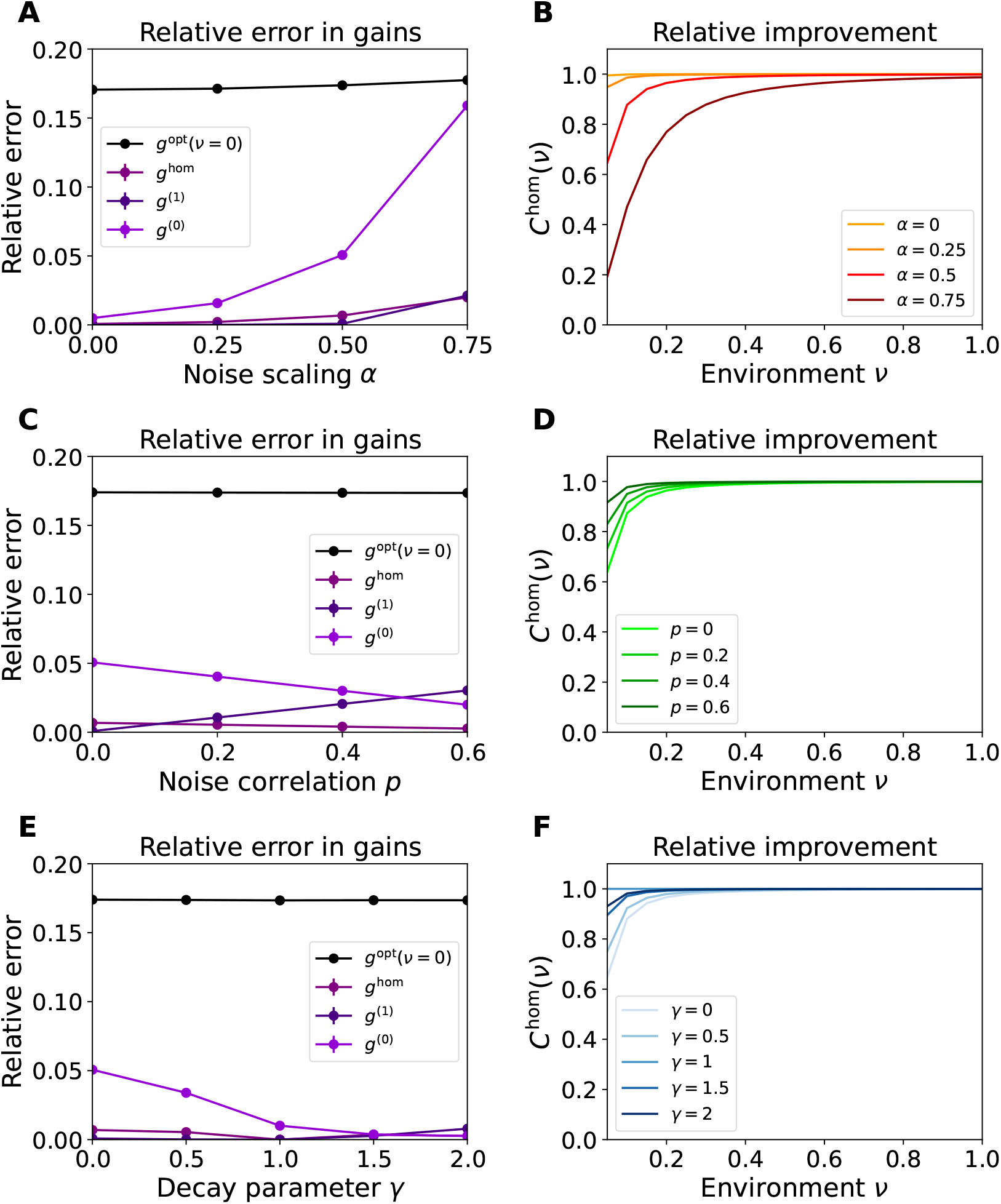
Accuracy of homeostatic approximation to optimal gains. (A) The relative error, Eq. (15), of different approximate solutions for the optimal gains, averaged across environments (the standard deviations of these values across environments were negligible relative to the average), as a function of the noise scaling parameter *α*. To give a sense of the scale of variation of the optimal gains across environments, the black line shows the relative change in the optimal gains between the most extreme environments (a measure of effect size), 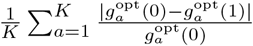. (B) Relative improvement in the objective for the homeostatic approximation ***g***^hom^ as measured by Eq. (16). Panels C and E (panels F and D) are the same as panel A (panel B), but for the uniformly correlated noise and the aligned noise models, respectively.

#### Uncorrelated power-law noise

First, we consider the uncorrelated noise model with general power-law scaling of noise strength with mean responses. In this case, the homeostatic solution ***g***^hom^, and the first order correction, ***g***^(1)^, had very low (*<* 3%) relative errors for all noise scaling exponents, *α*, while the relative error in ***g***^(0)^ becomes large for significantly super-Poissonian noise scaling, *i*.*e*., *α* = 0.75 (Fig. 9A).^9^ The relative improvement measure *C*^hom^(*ν*) for the homeostatic solution is also close to one, the optimal value, for all *α*, except for 0.75 (Fig. 9B). For the other approximate solutions too (Supp. Fig. 17A,B), the relative improvement decreases with increasing noise scaling. This is consistent with our analysis, since increasing *α* increases the size of Δ (see Sec. 2.4) and hence the influence of terms of higher order in Δ.

#### Uniform noise correlation coefficients

We now examine the effect of noise correlations on the accuracy of the approximate homeostatic solutions, first within the noise family with uniform noise correlation coefficients, denoted by *p*, and Poisson-like scaling (*α* = 1*/*2). For all *p*, the relative deviation of the homeostatic solution ***g***^hom^ from optimal gains was below 1% (Fig. 9C), while ***g***^(0)^ and ***g***^(1)^ were less accurate overall. Note that the accuracy of the homeostatic solutions ***g***^hom^ and ***g***^(0)^ increases with *p*, suggesting that positive noise correlations improve the optimality of homeostatic gains. The same trend can be seen in the relative improvement on the objective, *C*^hom^, for homeostatic gain adaptation, which increases with *p* (Fig. 9D).

#### Aligned noise

Lastly, we considered the algined noise model, in which noise has Poisson-like scaling (*i*.*e*., *α* = 1*/*2), and is correlated across clusters with the noise and signal covariance matrices sharing the same eigenbasis. In this noise family, the parameter, *γ*, controls the scaling of the eigenvalues of the noise covariance matrix with their rank, *n*, as 1*/n*^*γ*^. As in the previous cases, for all *γ*, the homeostatic solution ***g***^hom^ was very close to the optimal gains ***g***^opt^ (Fig. 9E), and preformed nearly equally (Fig. 9F). The case *γ* = 1, which corresponds to perfect alignment between signal and noise correlations, *W* = *ρ*, is particularly interesting. As noted in the previous section, this case corresponds to that of so-called information-limiting noise correlations (Moreno-Bote et al., 2014; Kanitscheider et al., 2015). As we show in App. B.5, in this case the optimal gains, without any approximation, lead to homeostasis and uniformisation of cluster firing rates. Thus, in this case ***g***^opt^ = ***g***^hom^ exactly, leading to zero relative error (Fig. 9E) for ***g***^hom^, and perfect relative improvement *C*^hom^ (Fig. 9F). For other cases, we can see that for increasing *γ* (at least for the values we simulated), which corresponds to more correlated and lower-dimensional noise (as noise power falls off faster as a function of eigenvalue rank), the homeostatic gains ***g***^hom^ tend to perform better.

Our numerical simulations across the three noise subfamilies (with varying noise power scaling, noise correlations, and noise spectrum) demonstrate that homeostatic strategies can - robustly and across a wide range of noise distributions - combat coding noise by effectively navigating the trade-off between energetic costs and coding fidelity.

Finally, note that adaptation in ***g***^hom^ uses only information local to each neuron, *i*.*e*., its firing rate. This implies that good performance does not require the use of complex regulation mechanisms which take into account how the encoding of each cluster relates to the population at large; homeostatic regulation, implemented by purely local mechanisms, is sufficient. In fact, homeostatic regulation can be implemented via a simple mechanism of synaptic scaling (Turrigiano et al., 1998; Turrigiano, 2008). Synaptic scaling involves the normalisation of synaptic weights onto a neuron in order to keep the total synaptic mass constant. In Appendix B.8, we demonstrate that, in a linear feedforward network, synaptic normalisation, via synaptic scaling, is both necessary and sufficient to allow for the propagation of homeostatic coding from the first layer to the subsequent ones. Thus, homeostatic coding in feedforward networks provides an additional normative interpretation of synaptic normalisation.

### 2.7 Homeostatic noisy DDCs

When exposed to over- or under-represented stimuli, neurons across sensory systems exhibit stimulus-specific adaptation whereby they decrease or increase their gains only for a limited subset of stimuli (Movshon and Lennie, 1979; Müller et al., 1999; Dragoi et al., 2000, 2002; Nelken, 2014). In particular, homeostatic adaptation in the primary visual cortex (V1) has been observed to be stimulus-specific (Benucci et al., 2013). In this section and the next (Sec. 2.8), we combine our efficient coding theory of homeostatic adaptation with a specific computational theory of neural representations. Then, in Sec. 2.9, we show that the resulting theory is able to account for stimulus-specific adaptation effects observed in V1, which have been difficult to account for under other efficient coding frameworks (see our discussion of previous attempts in Sec. 3). This theory thus provides a unified normative explanation for these two types of adaptation.

Recall that up to this point we have made no assumptions about the nature of cortical representations (beyond rate coding, and the condition described by Eq. (12)), and thus our framework has been independent of the computational goals of the circuit. We now apply our framework to a specific theory of neural representation and computation, namely the distributive distributional code (DDC) (Sahani and Dayan, 2003; Vertes and Sahani, 2018), introducing what we term a *homeostatic DDC*.

While there is ample behavioural and psychophysical evidence for the hypothesis that animal perception and decision making approximate optimal Bayesian probabilistic inference (Kersten et al., 2004; Stocker and Simoncelli, 2006; van Bergen et al., 2015; Geurts et al., 2022), our knowledge of the neural implementation of computations serving Bayesian inference (such as computations of posterior distributions and posterior expectations) are rudimentary at best. The DDC framework is one proposal for neural implementations of Bayesian computations underlying perception. In a DDC model, neural responses directly encode posterior expectations of the inferred latent causes of sensory inputs according to an internal generative model. This internal model mathematically relates the latent causes or latent variables, which we denote by ***z***, to the sensory inputs or stimuli, ***s***, via a family of conditional distributions, *f* (***s*** | ***z***), as well as a prior distribution, *π*(***z***), over the latent variables. The sensory system is assumed to implement a so-called *recognition model* corresponding to this generative model. Namely, given a stimulus ***s***, the task of the sensory system is to invert the generative process by calculating and representing the posterior distribution of latent variables given the current sensory input (see the schema in Fig. 10).

**Figure 10:**
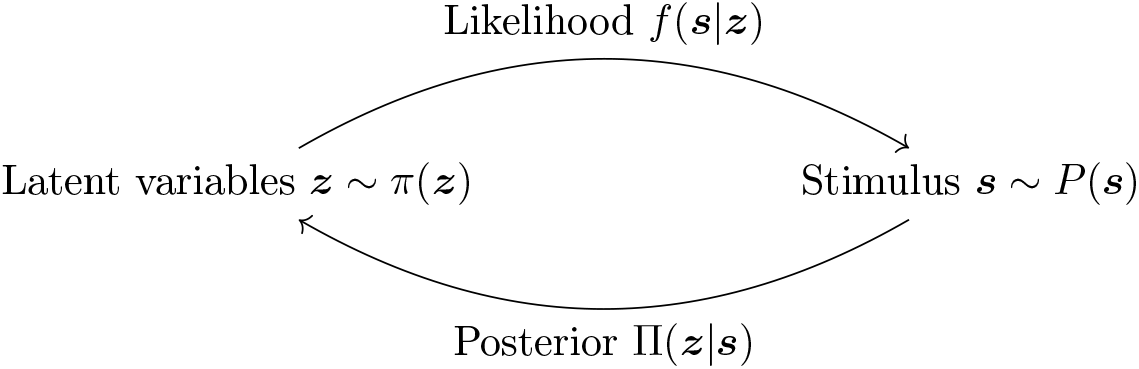
Schema of the generative and recognition models underlying a DDC. According to the internal generative model, sensory inputs, ***s***, are caused by latent causes, ***z***, that occur in the environment according to a prior distribution *π*(***z***). A conditional distribution *f* (***s*** | ***z***), the so-called likelihood function, describes how the latent causes generate or give rise to the sensory inputs ***s***. The task of the brain is to invert this generative process by inferring the latent causes based on the current sensory input, which is done by computing the posterior distribution Π(***z***|***s***).

In a DDC, neural responses are specifically postulated to directly represent the posterior expectations of the latent variables, ***z***, or rather, the posterior expectations of a sufficiently rich (and fixed) class of so-called kernel functions of ***z***. Specifically, in the standard DDC, each neuron, say neuron *a*, is assigned to a so-called kernel function of the latent variables, *ϕ*_*a*_(***z***), and its response is given by the posterior expectation 𝔼[*ϕ*_*a*_(***z***) | ***s***]. Here, we modify the standard DDC scheme in two ways. First, we assume single-trial neuron responses are corrupted by noise and represent the posterior expectations of the kernel functions only on average. Second, we adopt the key premise of our general efficient coding framework, namely that neural gains adapt in order to optimally mitigate the effect of neural noise on representational fidelity. Specifically, we assume this gain adaptation takes the homeostatic form, Eq. (14) (which we found approximates the optimal solution of the efficient coding problem for noise mitigation). These modifications lead to a new coding scheme which we call *noisy homeostatic DDC*. In this scheme, as in the standard DDC, the tuning curve of a neuron, with assigned kernel function *ϕ*_*a*_(***z***), is proportional to the posterior expectation 𝔼[*ϕ*_*a*_(***z***) | ***s***]. However, unlike the standard DDC, the corresponding constants of proportionality are not absolute constants but adapt (independently for different neurons) in such a way as to give rise to firing rate homeostasis. In other words, we assume the representational curve of neuron *a* is proportional to 𝔼[*ϕ*_*a*_(***z***) | ***s***], while its gain adapts across contexts or environments so as to maintain homeostasis. Since (under an ideal-observer internal model) the posterior expectation of a quantity averaged over stimuli is equal to that quantity’s prior expectation, we have 𝔼[𝔼[*ϕ*_*a*_(***z***) | ***s***]] = 𝔼[*ϕ*_*a*_(***z***)]. Therefore, keeping average firing rates constant at level *χ* (independent of the kernel function or the environmental statistics) requires the gains to be inversely proportional to the prior expectations, 𝔼[*ϕ*_*a*_(***z***)], and thus the tuning curve of neuron *a* to be given by

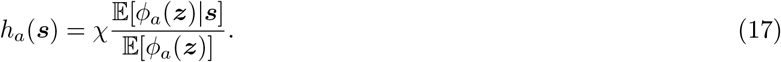

In other words, in noisy homeostatic DDCs, the neural tuning curves are (up to the absolute constant *χ*) the ratio of the posterior to the prior expectations of the neurons’ assigned kernel functions. Note that the prior expectation in the denominator, which by definition does not depend on the stimulus, ***s***, does depend implicitly on the environmental stimulus statistics. Indeed, this is also true of the posterior expectation in the numerator (as it combines information about the current stimulus with prior expectations about the latent causes of the stimulus input which can depend on the environment). Thus, in homeostatic DDCs, not only the gains, but also the shapes of tuning curves can adapt to changes in stimulus statistics, and thus vary across environments. This corresponds to a mixture of the two special scenarios sketched in the two panels of Fig. 1: as the stimulus distribution changes (Fig. 1A second row), (1) the tuning curve shapes change (Fig. 1B top row), as dictated by the computational goal of representing posterior expectations of latent variables under the changed statistics, and (2) the gains adapt in order to mitigate the effect of noise at minimal metabolic cost, resulting in mean response homeostasis (right columns). As we will see in Sec. 2.9, this combined adaptation can explain stimulus-specific adaptation.

### 2.8 Homeostatic DDCs can be implemented by divisive normalisation

We now show that for the special case when the DDC kernel function of neuron *a* is approximately a delta function, *ϕ*_*a*_(***z***) *≈ δ*(***z*** − ***z***_*a*_), the homeostatic DDC can be implemented by divisive normalisation with adaptive weights (Westrick et al., 2016). For *ϕ*_*a*_(***z***) *≈ δ*(***z*** − ***z***_*a*_), Eq. (17) becomes

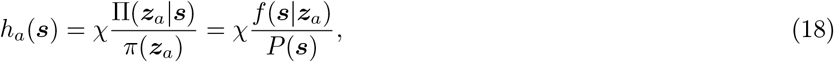

where in the last equality we used Bayes’ rule: Π(***z*** | ***s***) = *f* (***s*** | ***z***)*π*(***z***)*/P* (***s***). Thus, in this case, the tuning curve of neuron *a* is the ratio of the posterior distribution to the prior distribution both evaluated at that neuron’s preferred latent variable value, ***z***_*a*_. Accordingly, we call this coding scheme *Bayes-ratio coding*.

Divisive normalisation has been dubbed a canonical neural operation (Carandini and Heeger, 2012), as it appears in multiple brain regions serving different computational goals. Given the feedforward inputs, *F*_*a*_(***s***), the divisive normalisation model computes the tuning curve of neuron *a* as

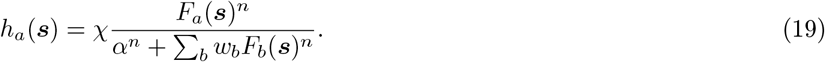

Here, *w*_*b*_ are a collection of so-called normalisation weights, and *α ≥* 0 and *n ≥* 1 are constants. To show that Bayes-ratio coding can be implemented by a divisive normalisation model, we note that *P* (***s***), which appears in Eq. (18), is given by the marginalisation of the joint density *P* (***s***) = ∫ *P* (***s, z***)*d****z*** = ∫ *f* (***s***|***z***)*π*(***z***)*d****z***. We approximate the integral by a sum as *P* (***s***) *≈* ∑_*b*_ *f* (***s***|***z***_*b*_)*π*(***z***_*b*_)Δ***z***_*b*_ =_*b*_ *w*_*b*_*f* (***s***|***z***_*b*_), where we defined the weights by *w*_*b*_ = *π*(***z***_*b*_)*δ****z***_*b*_.^10^ Substituting this approximation in Eq. (18), and regularising the denominator by an additive small constant *α >* 0, we obtain

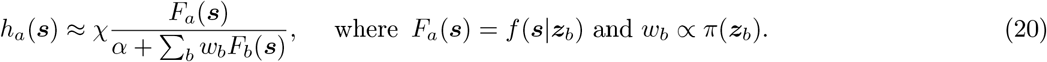

By Eq. (19) this corresponds to a divisive normalisation model with *n* = 1. Not only does this result show that implementing Bayes-ratio coding is biologically plausible, it also provides a normative computational interpretation to both the feedforward inputs and the normalisation weights of the divisive normalisation model, as the internal generative model’s likelihoods and prior probabilities, respectively. Note also that the normalisation weights of this model are adaptive, and vary between environments that have different latent variable distributions, *π*(***z***).

Finally, we consider the propagation of Bayes-ratio coding between populations. Suppose the generative model has the hierarchical structure ***z***^(2)^ ***z***^(1)^ → ***s***, and consider a feedforward network with two populations or layers that receives the stimulus and computes a Bayes-ratio encoding of the posterior distributions over ***z***^(1)^ and ***z***^(2)^. Assume that the first layer is implementing Bayes-ratio coding for the lower-level latent variables ***z***^(1)^ (*e*.*g*., using divisive normalisation Eq. (20)). We show in App. B.9 that if the feedforward synaptic weights into the second layer are given by 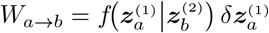, the downstream population implements Bayes-ratio coding for ***z***^(2)^ with linear neurons. This result is significant for three reasons. Firstly, while the downstream population is part of a *recognition* model that represents the posterior distribution of ***z***^(2)^, the synaptic weights needed to implement this representation only require knowledge of the *generative* probabilities (of ***z***^(1)^ given ***z***^(2)^). Secondly, the synaptic weights make no reference to the prior distribution over ***z***^(2)^. There is therefore no need to adapt the weights upon environmental changes that only affect the statistics of the higher-level latent causes ***z***^(2)^ but not the generative process (as encoded by *f*). Thirdly, this result can clearly be extended to a multi-level generative model by induction.

### 2.9 Homeostatic DDCs account for stimulus specific adaptation

We found that homeostatic DDCs can provide a normative account of stimulus specific adaptation (SSA). Here, we start by showing this in the special case of Bayes-ratio coding, which is mathematically simpler and provides good intuitions for a more general subset of homeostatic DDCs. We then discuss this more general case and build a homeostatic DDC model of empirically observed SSA in the primary visual cortex (V1) (Benucci et al., 2013).

According to Eq. (18), for Bayes-ratio codes the tuning curve of neuron *a* is given by *h*_*a*_(***s***) = *χf* (***s*** | ***z***_*a*_)*/P* (***s***). Suppose now that the stimulus distribution in the environment, *P* (***s***), changes due to a change in the statistics of the latent causes, *π*(***z***), while the causal processes linking the latent causes and observed stimuli, as captured by *f* (***s*** | ***z***), remain stable. In this case, by Eq. (18), the tuning curve of *all* neurons are modified by a multiplicative factor given by the ratio of the *P* (***s***) in the old environment to that in the new environment. This will lead to a multiplicative suppression of the response of all neurons to stimuli that are over-represented in the new environment relative to the old one, in a way that is independent of the neurons’ identity or preferred stimulus. Such a suppression thus constitutes a pure form of stimulus specific adaptation. In particular, neurons do not reduce their responsiveness to stimuli that are not over-represented in the new environment. Thus, tuning curves are suppressed only near over-represented stimuli (and may in fact be enhanced near under-represented stimuli), leading to a repulsion of tuning curve peak from over-represented stimuli. This repulsion is a typical manifestation of stimulus specific adaptation; in V1, for example, orientation tuning curves display a repulsion from an over-represented orientation (Movshon and Lennie, 1979; Müller et al., 1999; Dragoi et al., 2000, 2002; Benucci et al., 2013).

Heretofore we have implicitly made a so-called ideal observer assumption by assuming that the internal generative model underlying the DDC or Bayes-ratio code is a veridical model of the stimulus distribution in the environment. We now provide a significant generalization of the above results to the more general and realistic case of a non-ideal-observer internal model. As we will see, in this case, the effects of adaptation on tuning curves are captured by both a stimulus-specific factor as well as a neuron-specific factor. For a non-ideal observer model, there will be a discrepancy between the environment’s true stimulus distribution, *P*^*E*^(***s***), and the marginal stimulus distribution predicted by the internal generative model, *i*.*e*.,

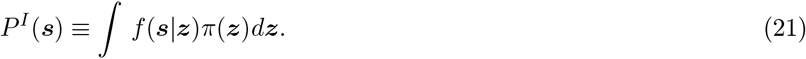

In App. B.10, we demonstrate that the Bayes-ratio tuning curves in this case are given by

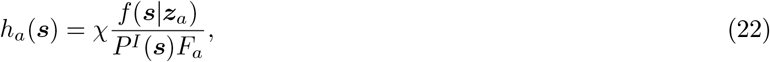

(instead of Eq. (18)) where

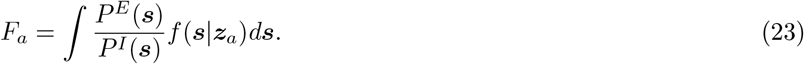

Note that, for ideal-observer internal models, *P*^*E*^ = *P*^*I*^, in which case the integral yields *F*_*a*_ = 1 due to the normalisation of *f* (***s*** | ***z***), and we recover Eq. (18). As we can see, the tuning curves neatly decompose into a “base tuning curve” *χf* (***s*** | ***z***_*a*_) multiplied by a stimulus-specific factor *P*^*I*^ (***s***)^−1^ and a neuron-specific factor 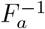. Consequently, if we consider a pair of environments indexed by *ν* = 0, 1, and assume the likelihood function of the internal model does not change between environments, the transformation of the tuning curves due to adaptation will be given by Eq. (24),

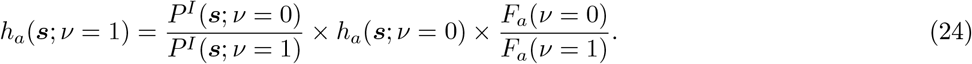

Thus, adaptation from environment *ν* = 0 to environment *ν* = 1 causes the tuning curves to transform via multiplication by a *stimulus specific factor P*^*I*^ (***s***; *ν* = 0)*/P*^*I*^ (***s***; *ν* = 1) and a *neuron specific factor F*_*a*_(*ν* = 0)*/F*_*a*_(*ν* = 1). Now suppose an environmental change results in an increase in the prevalence of an adaptor stimulus. As long as the internal model is a sufficiently good (if not perfect) model of *P*^*E*^(***s***), it should adapt its prior distribution, *π*(***z***), in a way that results in an increase of its predicted stimulus distribution, *P*^*I*^ (***s***), around the adaptor stimulus. This results in a stimulus specific factor that goes below 1 around the adaptor ***s***, thus causing suppression and repulsion of tuning curves away from the adaptor. Now consider a neuron whose preferred stimulus (roughly represented by ***z***_*a*_) is close to the adaptor stimulus. We further make the assumption that the change in the internal model, in response to the environmental change, is sufficiently conservative such that it leads to a smaller increase in *P*^*I*^ (***s***) around the adaptor, compared with the increase in the veridical *P*^*E*^(***s***) (this is a reasonable assumption especially for artificially extreme adaptation protocols used in experiments). In this case, we expect the ratio *P*^*E*^(***s***)*/P*^*I*^ (***s***) to increase in the support of *f* (***s*** | ***z***_*a*_), which (by our assumption about the preferred stimulus ***z***_*a*_) is near the adaptor. According to Eq. (23) this will lead to an increase in *F*_*a*_, and hence a suppressive neuron-specific factor *F*_*a*_(*v* = 0)*/F*_*a*_(*v* = 1) for neurons with preferred stimulus near the adaptor. In accordance with this picture, Benucci et al. (2013) found that homeostatic and stimulus-specific adaptation in V1 can indeed be modelled via separable stimulus-specific and neuron-specific factors.

Note that the assumption of constancy of the likelihood across the two environments need not be framed as a veridical assumption about the objective environmental change, but rather as an assumption about the inductive biases of the internal generative model, according to which it tends to model changes in the observed stimulus distribution, *P*^*E*^(***s***), as primarily resulting from changes in the statistics of the latent causes in the environment (which the model captures in *π*(***z***)), rather than in the causal process itself, as modelled by *f* (***s*** | ***z***). As long as this is good enough an assumption to result in a *P*^*I*^ (***s***) which changes in similar ways to *P*^*E*^(***s***) (and in particular exhibits an increase around adaptor stimuli), the conclusions reached above will hold.

In the case of DDC codes that use more general kernel functions, other than the delta functions underlying Bayes-ratio codes, we arrive at an analogous transformation, given by Eq. (140) in App. B.10. However, in this case, neuron-specific and stimulus-specific effects are mixed, and the wider the DDC kernel the stronger the mixing. Nevertheless, for kernels that are unimodal and sufficiently narrow, we expect an approximate factorisation of the effect of adaptation into a stimulus-specific and a neuron-specific suppression factor, as seen in Eq. (24).

We now apply this result to the experiments performed by Benucci et al. (2013). These experiments examined the effects of adaptation on orientation-tuned neurons in the cat primary visual cortex. Anaesthetised cats were shown a sequence of oriented gratings chosen randomly from a distribution that had an increased prevalence (3 to 5 times more likely) of one particular orientation (arbitrarily defined to be to 0^°^). A control group was exposed to a uniform distribution of gratings. After about 2 seconds (on the order of 50 stimulus presentations), V1 tuning curves had adapted, and exhibited both the suppressive and repulsive effects mentioned above.

To model these findings we constructed a homeostatic DDC model as follows, and fit it to the data from Benucci et al. (2013). We took the model’s stimulus and latent variable spaces to be the circular orientation space [−90, 90), and we took the model’s generative likelihoods to be given by translation invariant functions *f* (*s* | *z*) = *f* (*s* − *z*), proportional to the Gaussian distribution with standard deviation *σ*_*f*_, normalised over the circle.^11^ Likewise, we take the DDC kernel functions *ϕ*_*a*_(*z*) = *ϕ* (*z* − *z*_*a*_) proportional to the Gaussian distribution with standard deviation *σ*_*ϕ*_.

The distributions of stimulus orientations used in the experiment is highly artificial, in the way they jump discontinuously near the adaptor orientation. The internal prior distribution employed by the brain is more likely to be smooth, reflecting the brain’s inductive bias adapted to natural environments. Thus in our model we chose the internal prior to be a smooth distribution. The orientation distribution used in the experiment can be understood as a mixture of a uniform distribution, and a spike concentrated at the adaptor (Fig. 11A, blue curve). To obtain the smooth internal prior in our model, we replace the spike component of the experimental orientation distribution with a Gaussian distribution centred at the adaptor with standard deviation *σ*_*π*_ (Fig. 11A, red curve). Thus, our model has only three free parameters, *σ*_*f*_, *σ*_*ϕ*_ and *σ*_*π*_, *i*.*e*., the widths of the generative likelihood, kernel functions, and the internal orientation prior, respectively.

**Figure 11:**
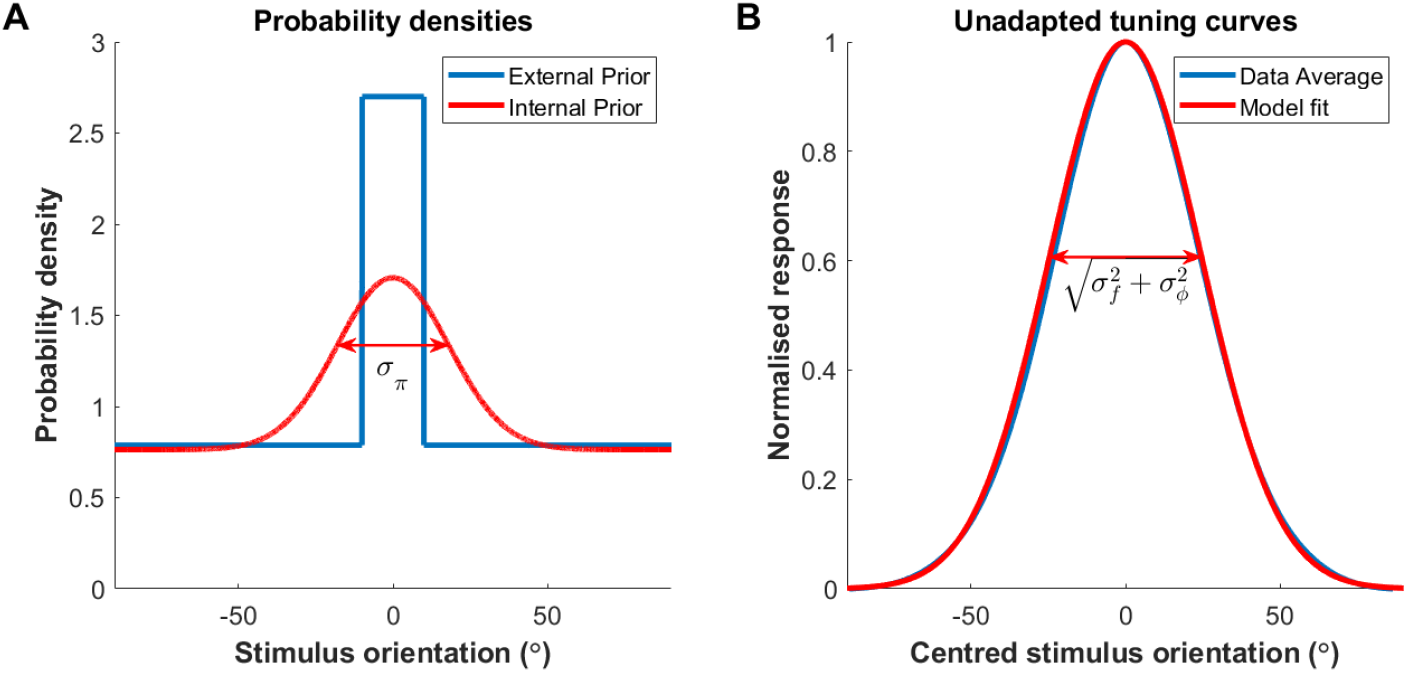
Components of the model for stimulus-specific adaptation in V1. (A) The distribution of stimulus orientations used in the experiments (Benucci et al., 2013), for an adaptor probability of 30% is shown in blue. We assume V1’s internal model use continuous prior distributions and thus we used instead a mixture of the uniform distribution with a smooth Gaussian, with standard deviation *σ*_*π*_, centered at the adaptor orientation (red curve). (B) The baseline tuning curves, *i*.*e*., the tuning curves adapted to the uniform orientation distribution. The blue curve was obtained by averaging the experimentally measured tuning curves adapted to the uniform orientation distribution, after centering those curves at 0^°^. The model’s baseline tuning curves (red) are given by the convolution of the Gaussian likelihood function, with width *σ*_*f*_, of the postulated internal generative model, and the Gaussian DDC kernel, with width *σ*_*ϕ*_, and thus are themselves Gaussian with width 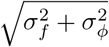. We fit this quantity to the experimentally observed baseline curves as described in Sec. 4.5.

Using the dataset from Benucci et al. (2013), we found the baseline tuning curves, *i*.*e*., the tuning curves of the population adapted to the uniform orientation distribution, by assuming that neural response is a function only of the difference between the preferred orientation and stimulus orientation (see Sec. 4.5). We then calculate the changes in the neurons’ preferred orientations (due to adaptation and the resulting tuning curve repulsion) in each of the experimental conditions (corresponding to different levels of over-representation of the adaptor stimulus). We then fit our model’s three parameters to both the unadapted tuning curves (Fig. 11B) and the changes in preferred orientation (see Sec. 4.5 for further details). The best fit values for the model parameters were found to be *σ*_*π*_ = 18^°^, *σ*_*f*_ = 14.2^°^, and *σ*_*ϕ*_ = 20^°^. The predictions of our model are compared to experimental data in Fig. 12. We found that the adaptive tuning curves within our model display suppression and repulsion away from the adaptor stimulus (Fig. 12A,D). In both the experimental data and our model, unadapted tuning curves exhibit a heightened average firing rate near the adaptor stimulus (Fig. 12B,E, blue curves). However, adaptation suppresses responses near the adaptor (Fig. 12A,D, red curves) to just the right degree to return the firing rates to the value before the environmental shift, leading to homeostasis of firing rates (as demonstrated by the uniformity of the red curves in Fig. 12B,E). Lastly, our model recapitulates the repulsion of preferred orientations found experimentally (Fig. 12C,F). Repulsion here is reflected by the change in preferred orientation having the same sign as the difference between the pre-adaptation preferred orientation and the adaptor orientation. As seen, in both the experiment and the model, repulsion is strongest for neurons with pre-adaptation preferred orientation about 30^°^ from the adaptor.

**Figure 12:**
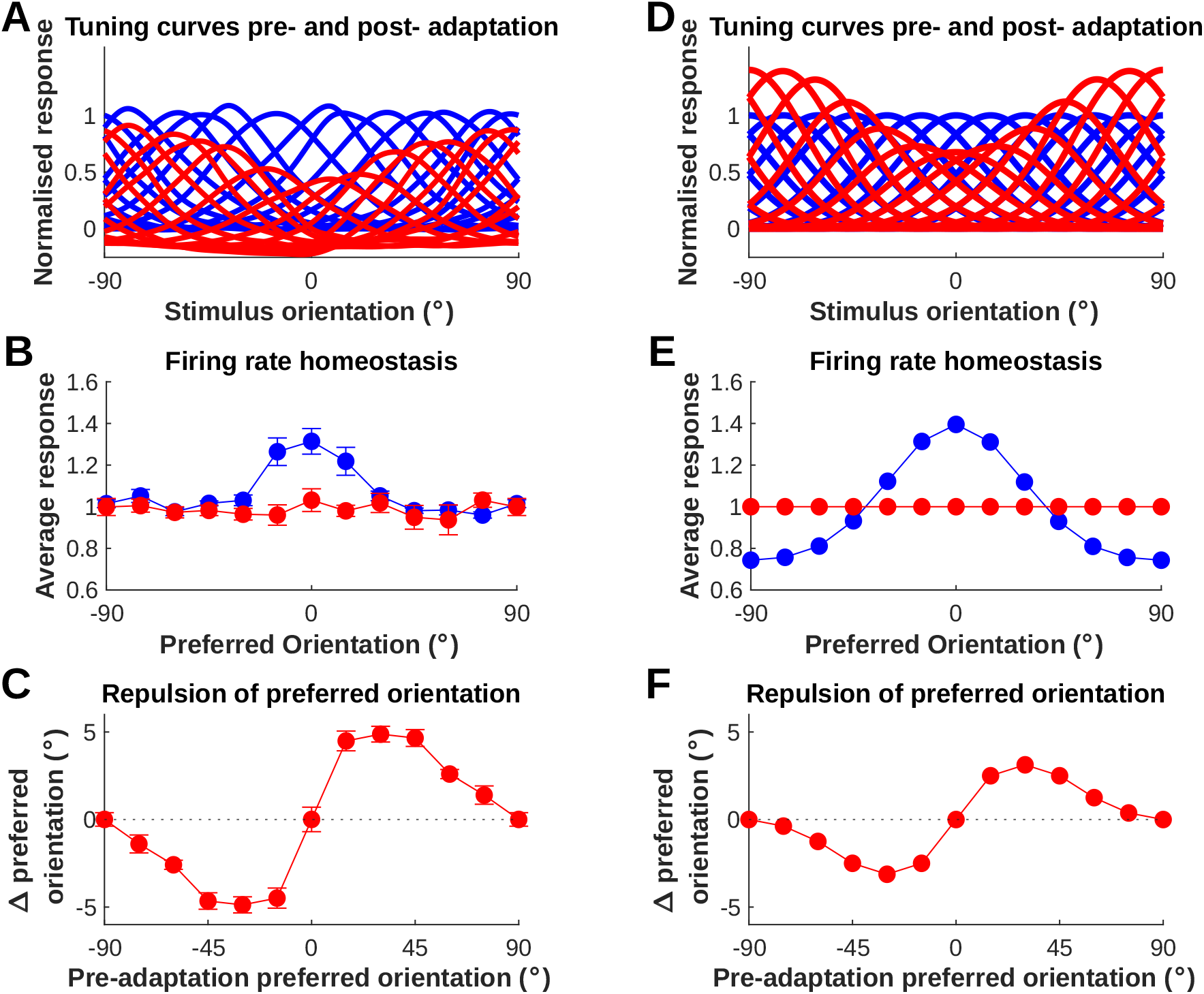
A homeostatic DDC model accounts for the observations of stimulus-specific adaptation in V1 Benucci et al. (2013). Panels A-C (left column) and E-F (right column) correspond to the experimental results and the predictions of our model, respectively. (A, D) The tuning curves for the adapted (red) and unadapted (blue) populations, averaged across experimental conditions. (B, E) The trial-averaged firing rates of the adapted (red) and unadapted (blue) populations exposed to the non-uniform stimulus ensemble. The trial-averaged population responses to the uniform stimulus ensemble were normalised to 1. (C, F) The repulsion of preferred orientations, obtained from the average tuning curves in panel A, as a function of the deviation of the neurons’ preferred orientation from the adaptor orientation.

In sum, the application of our homeostatic coding framework to a specific DDC model of Bayesian representations (which approximates what we have termed Bayes-ratio coding) explains quantitatively the empirical observations of both stimulus-specific and homeostatic adaptation in V1 (Benucci et al., 2013).

## 3 Discussion

We developed a theory of optimal gain modulation for combating noise in neural representations serving arbitrary computations. We demonstrated that — when mean neural firing rates are not too small; neurons are sufficiently selective and sparsely activated; and their responses form a high-dimensional geometry — the trade-off between coding fidelity and metabolic cost is optimised by gains that react to shifts in environmental stimulus statistics to yield firing rate homeostasis. Specifically, our theory predicts the homeostasis of the mean firing rate for clusters of similarly tuned neurons, while, at the same time, it accounts for the optimality of a broad distribution of firing rates within such clusters. By examining empirical estimates of parameters characterising cortical representations, we argued that the conditions necessary for the homeostatic adaptation to be optimal are expected to hold in the visual cortex and potentially other cortical areas. We further validated our approximation by demonstrating numerically that it performs well compared to the calculated optimal gains. Having developed a normative theory of neural homeostasis divorced from the computational aims of the neural population, we next showed how our theory can account for stimulus specific adaptation when coupled with distributed distributional codes (DDC). In particular, we focused on a special case of homeostatic DDC codes which we termed Bayes-ratio coding and showed that this model can account for stimulus specific adaptation effects empirically observed in the primary visual cortex. In the following, we will situate our work within the broader context of efficient coding theory, Bayesian neural representations, and alternative theories of stimulus specific adaptation. We will also discuss the shortcomings of the present work and possible directions for extension.

Within efficient coding theory, our work is closely related to that of Ganguli and Simoncelli (2014), but differs from it in important ways. Ganguli and Simoncelli (2014) considered a population of tuning curves, defined on a one-dimensional stimulus space, which was homogeneous and translationally invariant up to a warping (*i*.*e*., reparametrization) of the stimulus space and possible variations in gain. Their normative framework then optimised the neural gains as well as the stimulus warping (*i*.*e*., the reparametrization mapping from the original stimulus parameter to one over which the tuning curve population is homogeneous) to maximise a similar objective function to ours. Specifically, their objective employed the Fisher Information lower bound on the mutual information (Brunel and Nadal, 1998), as opposed to the Gaussian upper bound we used; our choice of the Gaussian upper bound was motivated by increased analytic tractability in the multidimensional stimulus case covered by our theory. In the case of unimodal tuning curves, they showed that the optimal warping amounts to a concentration and sharpening of tuning curves around stimulus values that are over-represented in the environment. On the other hand, in their setting, the optimal gains stay constant and do not adapt. Nevertheless, their solution also exhibits homeostasis of single-neuron firing rates.

Our framework extends and generalises that of Ganguli and Simoncelli (2014) in different ways. In their framework, the stimulus space is one-dimensional, and rigid constraints are placed on the shape and arrangement of the tuning curve population (they consider only unimodal or sigmoidal tuning curves, and assume that, up to a reparameterization of the stimulus space, all tuning curves have the same shape and are homogeneously placed). Wang et al. (2016) and Yerxa et al. (2020) have generalised the framework of Ganguli and Simoncelli (2014) to the case of alternative coding fidelity objective functions and high-dimensional stimuli, respectively; however, both of these studies maintain the same rigid constraint over the shape and arrangement of the tuning curve population. In contrast, our framework is applicable to tuning curves with heterogeneous (in particular multi-modal) shapes and arbitrary arrangement over a high-dimensional stimulus space; our homeostatic solution only requires sufficiently high signal-to-noise ratio (which is also the condition for the tightness of the mutual information lower-bound employed in Ganguli and Simoncelli (2014) or the approximation to the reconstruction-error loss used by Wang et al. (2016)). On the other hand, we only optimised the neural gains, and not the shapes or arrangement of the tuning curves, as we assumed the latter are determined by the computational goals of the circuit and not by the aim of optimally combating coding noise; in this way, our framework is agnostic to the computational goals of the population. Additionally, Ganguli and Simoncelli (2014) only considered the case of uncorrelated Poissonian noise. In contrast, our framework can handle correlated noise with general (non-Poissonian) power-law scaling. Furthermore, we analytically calculated first-order corrections to the homeostatic solution that arises in the high signal-to-noise ratio limit.

As noted above, the theory of Ganguli and Simoncelli (2014) predicts that tuning curves optimised to provide an efficient code under a stimulus distribution with over-represented stimuli cluster around those stimuli. This has been used to explain asymmetries in neural selectivities, such as the over-abundance of V1 neurons with preferred orientations near cardinal orientations, which are more abundant in natural scenes; or more generally, the aggregation of tuning curves around stimuli that are more prevalent in the natural environment. Many of these asymmetries have likely resulted from adaptation at very long (*e*.*g*., ontogenetic, or possibly evolutionary) timescales; however, long-term exposure on the order of few minutes to adaptor stimuli has also been found to result in the attraction of tuning curves towards adaptor stimuli (Ghisovan et al., 2009). This is in contrast to the repulsive effects seen in short-term adaptation (Movshon and Lennie, 1979; Müller et al., 1999; Dragoi et al., 2000, 2002; Benucci et al., 2013). It is likely that the effects of long-term exposure are the results of mechanisms which operate in parallel to those underlying short-term adaptation, with both types co-existing at different timescales. We can therefore interpret Ganguli and Simoncelli (2014) as a model of adaptation to natural stimulus statistics at long timescales. Our results obtained here are applicable to adaptation on shorter timescales (Müller et al., 1999; Dragoi et al., 2000, 2002; Benucci et al., 2013). In particular, when married with Bayesian theories of neural representation, our framework predicts repulsive effects around an adaptor stimulus, as shown in Sec. 2.9.

Westrick et al. (2016) also propose a model of the experimental findings of Benucci et al. (2013). Their model (which is not explicitly normative or based on efficient coding) uses divisive normalisation (Carandini and Heeger, 2012) with adaptive weights to achieve homeostasis and stimulus specific repulsion. As discussed above in Sec. 2.8, Bayes-ratio coding (a special case of homeostatic DDCs, which we showed can account for the findings of Benucci et al. (2013)) can be accomplished by such a divisive normalisation scheme. Our framework therefore yields a normative interpretation of the model of Westrick et al. (2016), and links divisive normalisation with Bayesian representations and efficient coding theory. In particular, our theory provides an interpretation of the normalisation weights of the divisive normalisation model as the internal prior over latent causes of sensory inputs. This prior should naturally adapt as stimulus statistics change across environments. Additionally, the feedforward inputs to the divisive normalisation model are interpreted as the likelihood of stimuli given latent causes.

Snow et al. (2016) developed two normative models of temporal adaptation effects in V1, based on statistics of dynamic natural visual scenes (natural movies). These models are based on generative models of natural movies in the class of mixtures of Gaussian scale mixture (MGSM) distributions. In MGSMs the outputs of linear oriented filters applied to video frames are assumed to be Gaussian variables, multiplied by positive scale variables. These scale variables can be shared within pools of filter outputs at different times (video frames) and orientations. The two models of Snow et al. (2016) differ according to which pools of filter outputs are able to share scale variables. In both models, V1 neural responses are assumed to represent the inferred Gaussian latent variables of the corresponding MGSM. While each model was able to account for some of the findings of Benucci et al. (2013), neither one could account for all. In particular, each of their two models could only account for either stimulus-specific or neuron-specific adaptation factors found by Benucci et al. (2013). In particular, only the model that accounted for neuron-specific adaptation was able to produce firing rate homeostasis. As we showed in Sec. 2.9, our framework, which combines homeostatic gain modulation with the DDC theory of representation, robustly accounts both for stimulus-specific adaptation and firing rate homeostasis. Furthermore, when there is a discrepancy between the internal model’s stimulus prior and the true environmental stimulus distribution, it additionally accounts for a neuron-specific adaptation factor as well (see Eq. (24)).

Tring et al. (2023) replicates the experiment performed in Benucci et al. (2013), this time in awake mice (rather than anesthetised cats). However, in contrast to the results of Benucci et al. (2013), Tring et al. (2023) do not find firing rate homeostasis in mouse V1. Additionally, Tring et al. (2023) find that adaptation mainly changes the scale of the population response vector, while minimally affecting its direction. Mathematically, this amounts to pure stimulus-specific adaptation without the neuron-specific factor found in by Benucci et al. (2013) - see Eq. (24). As we discuss in Sec. 2.7, a Bayes-ratio code based on an ideal-observer generative model also displays pure stimulus-specific adaptation with no neuron-specific factor. Our final model for Benucci’s data did contain a neuronal factor, because we used a non-ideal observer DDC (specifically, we assumed a smoother prior distribution over orientations compared to the distribution used in the experiment — which has a very sharp peak — as it is more natural given the inductive biases we expect in the brain). The resultant neuron-specific factor suppresses the tuning curves tuned to the adaptor stimulus. Interestingly, when gain adaptation is incomplete, and happens to a weaker degree compared to what is necessary for firing rate homeostasis, an additional neuronal factor emerges that is greater than one (*i*.*e*., facilitatory) for neurons tuned to the adaptor stimulus. These two multiplicative neural factors can potentially cancel each other approximately; such a theory would thus predict both deviation from homeostasis and approximately pure stimulus-specific adaptation. We plan to explore this possibility in future work.

Benucci et al. (2013) find that in cases where the adaptor probability is particularly extreme (*i*.*e*., 50%), homeostasis is imperfect, and neurons tuned to the adaptor display higher firing rates. These results cannot be accounted for within our homeostatic DDC framework, since we take as a starting point homeostatic regulation of firing rates in our model. However, note that our efficient coding framework does predict deviation from homeostasis when signal-to-noise ratio falls (see Sec. 2.4). Thus, a more complete model in which we both specify DDC representational curves and numerically optimise gains according to our efficient coding objective may be able to account for the observed deviation from homeostasis.

Our approach has relied on a number of assumptions and simplifications; relaxing or generalising these assumptions provide opportunities for future research. Firstly, our toy model (see Sec. 2.3) imposed a cluster structure on the neural tuning curves, whereby neurons within a cluster are similarly tuned, while neurons in different clusters have distinct tuning. We found that total firings rates are equalised across clusters (Sec. 2.2), but the optimal rates of individual neurons within clusters span a broad range (Sec. 2.3). This results from the fact that the efficient coding objective function depends sharply on total cluster firing rates, but changes only slightly when the partition of the cluster firing rate among the cluster’s neurons changes. In reality, tuning similarity has a more graded distribution. As the distinction between clusters with different tuning (or similarity of tuning within clusters) weakens, higher order terms in our expansion of the objective Eq. (27) cannot be neglected, and hence the justification for dividing the problem into two separate, cluster-level vs. within-cluster, problems breaks down. However, arguably, in a more realistic model with a more graded variation in neural signal correlations, the optimisation landscape will still contain shallow directions (*i*.*e*., directions over which the objective changes slowly), corresponding to partitions of firing among similarly tuned neurons, and non-shallow directions corresponding to total firings of such groups of neurons; if so, we would expect a similar solution to emerge, whereby similarly tuned neurons display heterogeneous firing rates, with the total firing rate of such a group of neurons displaying approximate homeostasis.

Secondly, we showed analytically that, at the level of cluster responses, homeostasis emerges universally in the high signal-to-noise regime. In general, firing rate homeostasis is no longer optimal in the low signal-to-noise ratio limit, as correction terms to our solutions become large. However, as we argued in Sec. 2.5, based on empirical values we expect the signal-to-noise ratio within cortex to be sufficiently high for our results to hold. Generally, within efficient coding theory (except for simple cases such as linear Gaussian models), the regime of arbitrary signal-to-noise ratio is analytically intractable. Many approaches are therefore limited to the regime of high signal-to-noise ratio. For example, the Fisher Information Lower Bound used frequently within efficient coding theory breaks down outside of this regime (Yarrow et al., 2012; Bethge et al., 2002; Wei and Stocker, 2016). Thus, inevitably, exploring the low signal-to-noise ratio regime would require numerical simulation. However, such numerical simulations require concrete models of signal and noise structure. The manner in which optimal coding deviates from homeostatic coding will not be universal across these models, but depend on each model’s specific details. This limits our ability to draw general conclusions from numerical simulations, which is why we have chosen not to pursue that strategy extensively here.

Thirdly, our analysis here considered a specific class of noise models. These noise models include those of particular biological relevance, such as information-limiting noise correlations (Moreno-Bote et al., 2014; Kanitscheider et al., 2015), signatures of which have been found in the cortex (Rumyantsev et al., 2020), and power-law variability (Goris et al., 2014) (see Sec. 2.6). As we noted, our homeostatic solution is particularly well suited to the case of information-limiting noise correlations, performing optimally in that case (see App. B.5). The core ideas of our framework and derivation could potentially be applied to an even wider class of noise models.

Fourthly, up to Sec. 2.7, we make no assumptions about what the neural population is attempting to represent via the representational curves. We then apply our framework to a specific theory of Bayesian encoding (namely DDCs), and develop the new idea of Bayes-ratio coding. There is therefore an opportunity to apply our framework to other theories of neural representation, and in particular to alternative theories of Bayesian representations, such as Probabilistic Population Codes (PPCs) (Beck et al., 2007).

Finally, it is possible to generalise our theoretical framework (both the general efficient coding framework and its combination with frameworks for Bayesian inference) to the case of temporally varying stimuli and dynamic Bayesian inference. In particular, Młynarski and Tkacik (2022) have applied optimal gain modulation, according to efficient coding (to reduce metabolic cost), to sparse coding (an example of a probabilistic model with latent variable representation), in order to model gain modulation by attention. Accordingly, their efficient coding objective was task-dependent (*i*.*e*., depended on the choice of latent variables that are desired to be in the focus of attention). However, we are not aware of similar work that has applied such a combination to model the dynamics of task-independent (*i*.*e*., bottom up, rather than top-down) adaptation.

In summary, we showed that homeostatic coding can arise from optimal gain modulation for fighting corruption by noise in neural representations. Based on this coding scheme, we derived a novel implementation of probabilistic inference in neural populations, known as Bayes-ratio coding, which can be achieved by divisive normalisation with adaptive weights to invert generative probabilistic models in an adaptive manner. This coding scheme accounts for adaptation effects that are otherwise hard to explain exclusively based on efficient coding theory. These contributions provide important connections between Bayesian representation, efficient coding theory, and neural adaptation.

## 4 Methods

### 4.1 Specification of environments for simulations

For our numerical simulations, we used *K* = 10, 000 units. To generate the *K × K* signal correlation matrix, *ρ*(*ν*), for the different environments in our numerical simulations, in Sec. 2.2, we first generate a covariance matrix Σ(*ν*), and let *ρ*(*ν*) be the corresponding correlation matrix. The covariance matrix Σ(*ν*) is in turn constructed such that it has (1) the same eigenspectrum for all *ν*, with eigenvalues that decrease inversely with rank, as observed in V1 (Stringer et al., 2019), and (2) an eigenbasis that varies smoothly with *ν*. Thus we defined Σ(*ν*) = *U* (*ν*)Λ_1_*U* (*ν*)^T^, where Λ_1_ = diag(1, 1*/*2, 1*/*3, …, 1*/K*) and *U* (*ν*) is an orthogonal matrix that was generated as follows. We randomly and independently sample two *K×K* random iid Gaussian matrices *R*_0_, *R*_1_ *~* 𝒩 _*K×K*_(0, *I*_*K×K*_) and obtain symmetric matrices 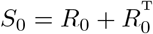 and 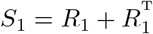. As is well known, the eigen-basis (represented by an orthogonal matrix) of such a random Gaussian matrix is distributed uniformly (*i*.*e*., according to the corresponding Haar measure) over the orthogonal group *O*(*K*). For the extreme environments, *ν* = 0 and 1, we let *U* (*ν*) be the matrix of eigenvectors of *S*_0_ and *S*_1_, when ordered according to eigenvalue rank. For 0 *< ν <* 1, we let *U* (*ν*) be the matrix of the eigenvectors (again ordered according to eigenvalue rank) of the interpolated symmetric matrix *S*(*ν*) = (1 − *ν*)*S*_0_ + *νS*_1_. Thus, for the extreme environments the eigenbases of the covariance matrices Σ(0) and Σ(1) are sampled independently and uniformly (*i*.*e*., from the Haar measure), and the eigenbases of Σ(*ν*) for intermediate environments (*i*.*e*., for 0 *< ν <* 1) smoothly interpolate between these.

In the case of simulations for aligned noise, we also generate a noise correlation matrix *W* (*ν*) which has an approximately 1*/n*^*γ*^ spectrum. Ideally, *W* and *ρ* would have the same eigen-basis. However, this is impossible, since *W* and *ρ* are both correlation matrices. Instead, we generate *W* (*ν*) by normalising the positive definite matrix *U* (*ν*)Λ_*γ*_*U* (*ν*)^T^ where Λ_*γ*_ = diag(1, 1*/*2^*γ*^, 1*/*3^*γ*^, …, 1*/K*^*γ*^)

### 4.2 Analysis of the clustered population

Here, we provide a summary of the analysis of the clustered model introduced in Sec. 2.3 (for derivations and a more detailed account see App. B.3). In this model, a population of *N* neurons are sorted into *K* clusters. Neurons within a cluster have very similar stimulus tuning, but neurons in different clusters are tuned differently. More concretely, we will assume that the representational curves of the neurons, indexed by *i*, belonging to cluster *a*, are small perturbations to the same representational curve characterising that cluster; *i*.*e*.,

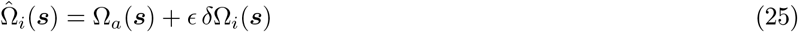

where *ϵ* is the perturbation strength parameter controlling the deviation from perfect within-cluster similarity. Below, hatted symbols 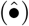 denote variables defined analogously to Eq. (7)–(8) but for single neurons, with Ω_*a*_(***s***) replaced by 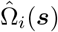.

We assume single-neuron gains adapt to optimise an objective ℒ _pop_, defined analogously to the objective ℒ, Eq. (9), but for individual neurons rather than clusters. In the case of uncorrelated noise with Poisson-like scaling, this objective is given by

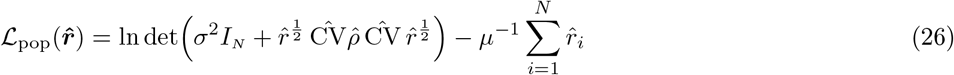

where 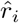 are the mean firing rates of individual neurons, given by 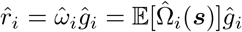; note that maximising ℒ_pop_ in terms of the single-neuron gains is equivalent to maximising it in terms of 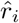. In App. B.3, we show that this objective can be expanded in *ϵ* as

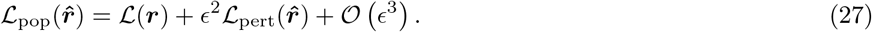

Here, the leading order term, ℒ (***r*** = *ω****g***), is a cluster-level objective and is given byLEq. (9), now viewed as a function of the cluster rates *r*_*a*_, *i*.*e*., the total mean firing rate of neurons in a cluster; thus 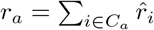 where *C*_*a*_ is the set of neurons in cluster *a*. Note that this term is a function only of the cluster firing rates. Thus, for *ϵ* = 0 (perfect similarity within clusters), the efficient coding objective function is blind to the precise distribution of single-cell firing rates, 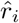, among the neurons of a cluster, as long as the total rate of the cluster is held fixed. This is because for clusters of identically-tuned and independently-firing neurons with Poisson-like noise, the cluster can be considered as a single coherent unit whose firing rate is given by the sum of the firing rates of its neurons. Thus at zeroth order, the distribution of individual neuron rates are free and undetermined by the optimisation, as long as they give rise to optimal cluster rates.

According to Eq. (27), at small but non-zero 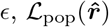 is still dominated by ℒ; we thus expect this term to play the dominant role in determining the cluster rates, with terms of order *ϵ*^2^ and higher having negligible effect. However, the perturbation term 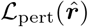 breaks the invariance of the loss with respect to changes in the distribution of single neuron gains (or rates) within clusters. We will therefore approximate the maximisation of the total objective function ℒ_pop_ as follows. We first specify the cluster rates by maximising ℒ (***r***), obtaining the optimal cluster rates 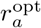. We then specify the rates of individual neurons within the population by maximising the leading correction term in the objective, 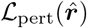, subject to the constraint that the single-neuron rates in each cluster sum to that cluster’s optimal rate. We reinterpret the optimisation in the previous sections to correspond to the first stage of this two-stage optimisation, with population units there corresponding to clusters of similarly tuned neurons. As we have seen in Sec. 2.2, optimisation of the cluster-level objective, ℒ, can result in homeostasis and uniformisation across clusters.

As for the distribution of individual neuron rates, we show in App. B.3.5 (see the derivation of Eq. (87)) that, in the parameter regime (see Secs. 2.4 and 2.5) in which cluster-level homeostasis and uniformisation occurs, the perturbation objective ℒ _pert_ decomposes into a sum of objectives over different clusters. Thus, in the second stage of optimisation the problem decouples across clusters, and (as shown in App. B.3.5) the optimal single-neuron rates in cluster *a* maximise the objective

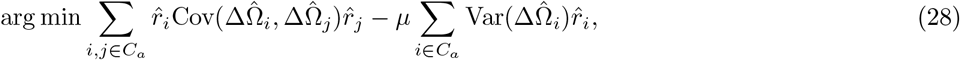

subject to the constraint

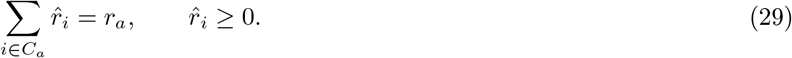

Here ΔΩ_*i*_ is defined to be the centered, zero-mean version of *δ*Ω_*i*_:^12^

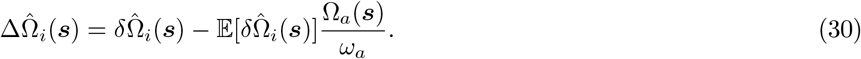

Note that the optimisation problem, Eqs. (28)–(29), is a quadratic program. Moreover, the terms in the objective for this quadratic program are homogeneous in the sense that if the information-energy trade-off parameter *µ* is scaled a factor, the optimal solutions for 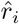 scale with the same factor. In our cluster-level simulations Sec. 2.2 we used *µ* = 10 (in agreement with our biological estimates in Sec. 4.4 for mean spike count of clusters of 20 V1 neurons in a coding interval of 0.1 seconds). That dimensionless *µ* (which in the homeostatic approximation determines the mean spike count of clusters over the coding interval) thus corresponds to a total cluster rate of 100 Hz, and using this value for *µ* in Eqs. (28)–(29) the solutions for 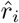 can be interpreted as being firing rates in units of Hz.

### 4.3 Within-cluster optimisation

To calculate the single-neuron firing rate distributions in Fig. 8, we solved the optimisation problem Eqs. (28)–(29) numerically for 10,000 randomly generated covariance matrices (note that different instances of the optimisation can be thought of as characterising different clusters within the same population). In general, the covariance matrix, 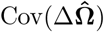 (which defines this optimisation problem), depends on the stimulus distribution, the cluster tuning curve, and properties (*e*.*g*., degree of smoothness) of the intra-cluster variations in single-cell tuning curves. In our simulations, instead of making specific arbitrary choices for these different factors, we used a minimal toy model where the covariance matrix was generated as the matrix of inner products of *k* random vectors in a *D*-dimensional space; specifically, we generated this matrix as the the Gram matrix of *k* random *D*-dimensional vectors with isotropic Gaussians distributions, *i*.*e*., 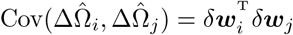, where *δ****w***_*i*_ *~* 𝒩 (0, *I*_*D*_). In App. B.3.6 we show that *D* can be interpreted as the dimensionality of the space of independent perturbations to the cluster-wide tuning curve Ω_*a*_ which vary significantly on the portion of stimulus space to which the cluster tuning curve responds (below, we will refer to *D* as the effective tuning curve dimension). Additionally, we fixed the cluster rate to *µ* = *k ×* 5Hz, such that the average firing rate per single neuron is 5Hz, and fixed *σ*^2^ = 1 (corresponding to a Fano factor of 1). For various values of *k* and *D*, we used the above method to generate 10, 000 samples of the covariance matrix Cov(ΔΩ). For each such sample, we then solved the problem (28–29) numerically using the `cvxpy` package in Python (Diamond and Boyd, 2016). Neurons with firing rates less than 10^−4^Hz had their firing rate set to zero. We aggregated the set of non-zero rates obtained from each of these optimisations to create the histograms in Fig. 8.

### 4.4 Estimation of biological parameters justifying the homeostatic approximation

In this section we provide the details for our estimates of cortical mean firing rates and noise scaling used in Sec. 2.5.

#### Firing rate estimates

A wide range of mean firing rates for individual neurons have been reported in cortex. Here we focus on firing rates in rodent V1 during free behaviour (which tends to be lower compared to rates in cat or monkey cortex). Reported values tend to range from 0.4 Hz (Greenberg et al., 2008) to 14 Hz (Parker et al., 2022), depending on the cortical layer or area or the brain state, with other values lying in a tighter range of 4-7 Hz (Hengen et al., 2013; Szuts et al., 2011; Torrado Pacheco et al., 2019). Therefore, for rodent V1, we take a mean firing rate of 5 Hz to be a reasonable rough estimate.

#### Scaling of noise variance with mean response

A number of studies (Shadlen and Newsome, 1998; Gershon et al., 1998; Moshitch and Nelken, 2014; Koyama, 2015) have characterised the relationship between the trial-to-trial mean and variance of neural spike counts as a power law of the form

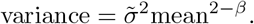

and have provided best fit estimates of 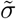 and *β* in different species and cortical areas. We have summarised these in Table 1.

**Table 1:** Table summarising empirical findings from four papers which estimate a scaled power-law relationship relationship between mean and variance of cortical spike count responses. To find the values in the last column we adjust the reported 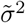 to account for different coding intervals and cluster sizes used in each experiment (see Eq. (31)), and assume *βµ* = 10.

| Paper | Area | $\tilde{\sigma}^2$ | $\beta$ | Time interval (s) | $\sigma^2/(\beta\mu)^\beta$ |
| --- | --- | --- | --- | --- | --- |
| Shadlen and Newsome (1998) | MT (rhesus monkey) | 0.4 | 0.7 | 0.5 | 0.05 |
| Gershon et al. (1998) | V1 (rhesus monkey) | 1.30 | 0.57 | 0.27 | 0.15 |
| Gershon et al. (1998) | V1 (rhesus monkey) | 1.82 | 0.82 | 0.27 | 0.19 |
| Gershon et al. (1998) | IT (rhesus monkey) | 1.36 | 1.18 | 0.3 | 0.13 |
| Moshitch and Nelken (2014) | Auditory cortex (cat) | 1.37 | 0.88 | 0.05 | 0.12 |
| Koyama (2015) | MT (rhesus monkey) | 4.57 | 1.43 | 0.5 | 0.31 |

While the power-law exponent *β* is directly identifiable with the same parameter in our noise model, we need to adjust the empirical estimates of 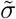 in Table 1 in order to obtain estimates for our model parameter *σ*, for two reasons. First, the duration of the time interval in which spikes were counted in the above papers differs from our assumed coding interval of 0.1*s*. We therefore must adjust the reported values to be for an interval of this size. We do this by assuming temporal homogeneity over the timescales of interest; specifically we assume that mean and variance are linear in the time interval. Second, we must also adjust the data to account for the fact that our responses are for clusters of *k* = 20 similarly tuned neurons. We do this by assuming uncorrelated response noise within each cluster, making mean and variance both linear functions of cluster size. Our adjustments do not affect the value of *β*, but do change the value of the pre-factor 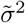 Specifically, we perform the transformation:

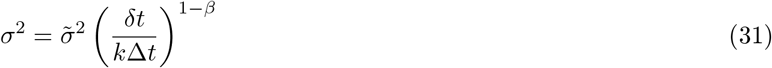

where *δt* is the reported time interval, and Δ*t* is our coding interval, Δ*t* = 0.1s. Fig. 13 shows the adjusted values on a scatter plot. Using the adjusted values, *σ*^2^, and assuming that *µβ* = 10 (to maintain a firing rate of 5Hz per neuron), we can find the value of *σ*^2^*/*(*βµ*)^*β*^. These are reported in the furthest right column of Table 1.

**Figure 13:**
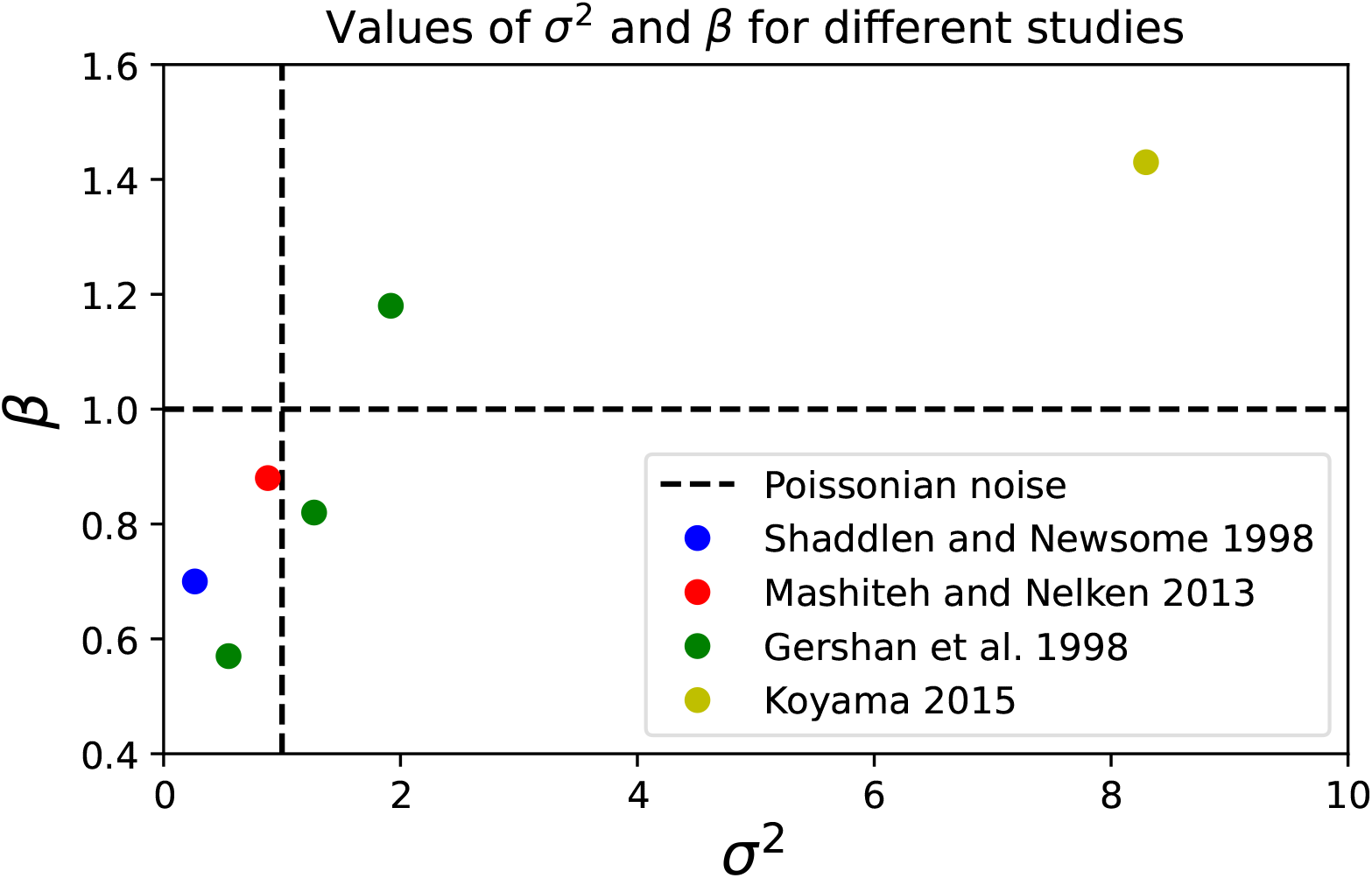
Empirically estimated values of the noise scaling parameter *β* and the base noise-level *σ*^2^, after adjusting for coding interval duration and cluster size using the formula Eq. (31). The dashed lines show the values for Poissonian noise (*σ*^2^ = 1, *β* = 1).

#### Sufficiently selective responses

The coefficient of variation CV is a measure of neural selectivity, and therefore is related also to measures of sparseness of neural responses, or so-called coding level. To see this, consider a toy model in which the cluster responds at a fixed level to a fraction *p* of stimuli and is silent otherwise. In this case, CV^2^ = (1−*p*)*/p* ≈1*/p* for small *p*. Our condition therefore requires that neurons are sufficiently selective in their responses and respond only to a small fraction of stimuli. Lennie (2003) places the fraction of simultaneously active neurons (which we use as a proxy for the response probability of a single cluster) at under 5%. For our toy model, this yields the estimate CV^2^ *≈* 20. The binary distribution in the toy model is particularly sparse, and so we take CV^2^ *≈* 10 as a more conservative estimate.

#### High-dimensional signal geometry

Assuming, for simplicity, that the coefficients of variation are approximately the same for all clusters, we see that the expression for Δ_*aa*_ are proportional to the diagonal elements of *ρ*^−1^*W*. To estimate the latter, we will assume that these elements are all comparable in size,^13^ and will therefore estimate their average value, which is given by the normalised trace 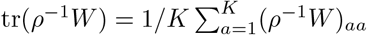. By the invariance of trace, this can equally be characterised as the mean eigenvalue of *ρ*^−1^*W*. We estimated this trace for the aligned noise family (corresponding to information-limiting noise correlations, but also including uncorrelated noise) which we numerically studied in Sec. 2.2 and Sec. 2.6, and a biologically-motivated form of *ρ* used there. In the aligned noise model, *W* and *ρ* share the same eigenbasis, but have different eigenvalue spectra. Our choice of *ρ* has spectrum that scales like 1*/n* with the rank of the eigenvalue (based on the findings of Stringer et al. (2019) that the signal covariance matrix of mouse V1 neural populations responding to natural stimuli possesses an approximately 1*/n* spectrum), while the spectrum of *W* scales like 1*/n*^*γ*^ for a positive exponent parameter *γ*.

We show in App. B.6 that, for 0 *≥ γ ≥* 1 (relatively high-dimensional and weakly correlated noise), we have 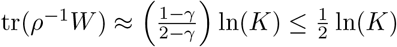. The upper bound is saturated in the case of uncorrelated noise (which correspond to *γ* = 0 yielding *W* = *I*). In the other extreme where *γ* = 1, we have *W* = *ρ* or *ρ*^−1^*W*, which yields a normalised trace of 1; which does not scale with *K*. On the other hand, for *γ >* 1 (relatively low-dimensional and strongly correlated noise), tr(*ρ*^−1^*W*) decays like *K*^− min(*γ*−1,1)^ as *K* grows. Thus, in this case, the estimate of Δ_*aa*_ is smaller than that of Eq. (13). However, this case is arguably not biologically relevant, due to the fact that neural responses always contain some amount of “private” uncorrelated noise. Mathematically this would contribute an additive correction to *W* proportional to the identity matric. In the case *γ >* 1 (where noise eigenvalues decay faster than signal eigenvalues) this term, even when the size of the correction is small, prevents the inverse power-law scaling with *K*. Therefore we did not use this case in the Results section in our estimates of Δ_*aa*_.

### 4.5 Fitting a homeostatic DDC to Benucci *et al*. 2013

The data we obtained from Benucci et al. (2013) comprised 11 datasets corresponding to different experimental sessions. In each dataset, the recorded neural population was clustered into 12 groups based on preferred orientation. The datasets contained the trial-averaged responses of these clusters to gratings of different orientation (which took one of 6 or 12 possible values depending on the dataset). In each experimental context, the grating orientations were randomly drawn from a distribution that was either uniform or biased; in the latter case, one particular grating (arbitrarily defined as 0°) had higher prevalence, by either 30%, 35%, 40%, or 50%. We discarded all datasets with a 50% prevalence, since (Benucci et al., 2013) report that homeostasis was less perfect in this case, and they similarly discarded it from their main analysis. After exposure to the distribution of gratings, the responses (*i*.*e*., spike count) of each cluster to a test set of 20 oriented gratings were then measured. In each dataset, the trial-averaged tuning curves had been normalised by an affine transform (see Benucci et al. (2013) for details) such that tuning curves in the uniform stimulus ensemble context ranged from 0 to 1. We further interpolated and up-sampled all normalised tuning curves using a cubic spline (Fig. 12A).

To fit our model parameters (*σ*_*f*_, *σ*_*π*_, and *σ*_*ϕ*_), we started by centering the cluster tuning curves in the uniform ensemble context at 0° and averaged across clusters with different preferred orientations, yielding a single average tuning curve (Fig. 11B, blue). We fit the resultant curve with a Gaussian (Fig. 11B, red) with a standard deviation matched to 1/4 the width of this curve at height *e*^−2^. Subsequently, 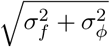 (which is the model’s prediction for the standard deviation of the Gaussian tuning curves in the uniform ensemble context) was constrained to equal this standard deviation. This reduces the number of free model parameters to 2, which we choose to be *σ*_*π*_ and *σ*_*f*_. We found the best value of these parameters by fitting the curves for repulsion of tuning curves (adaptation-induced change in preferred orientation vs. pre-adaptation preferred orientation – Fig. 12C and F) between the model and experimental data. Specifically, we performed a grid search over values of *σ*_*π*_ and *σ*_*f*_ and chose the pair of values which minimised the sum of the absolute differences between the empirical and model curves.

We obtained the empirical repulsion curves (Fig. 12C) as follows. For each preferred orientation cluster and context (*i*.*e*., exposure to a uniform or biased ensemble), a smoothing kernel was applied before using cubic interpolation to generate up-sampled tuning curves. The smoothing kernel was applied to ensure that none of the tuning curves were multimodal (multimodality was a minor effect — reasonably attributable to finite-trial averaging noise — but could have nevertheless introduced relatively large noise in the estimated preferred orientations). The argmax of these tuning curves was found to give the preferred orientation. The preferred orientation was then compared across conditions (uniform ensemble vs. biased ensemble) to give a change in preferred orientation. These were then averaged across contexts with the same adaptor probability to obtain the repulsion curve for each adaptor probability.

## 5 Acknowledgements

We thank Máté Lengyel for helpful discussions and comments on the manuscript. EY was supported by the UKRI Engineering and Physical Sciences Research Council Doctoral Training Program grant EP/T517847/1. YA was supported by UKRI Biotechnology and Biological Sciences Research Council research grant BB/X013235/1.

## A Extended data figures

**Figure 14:**
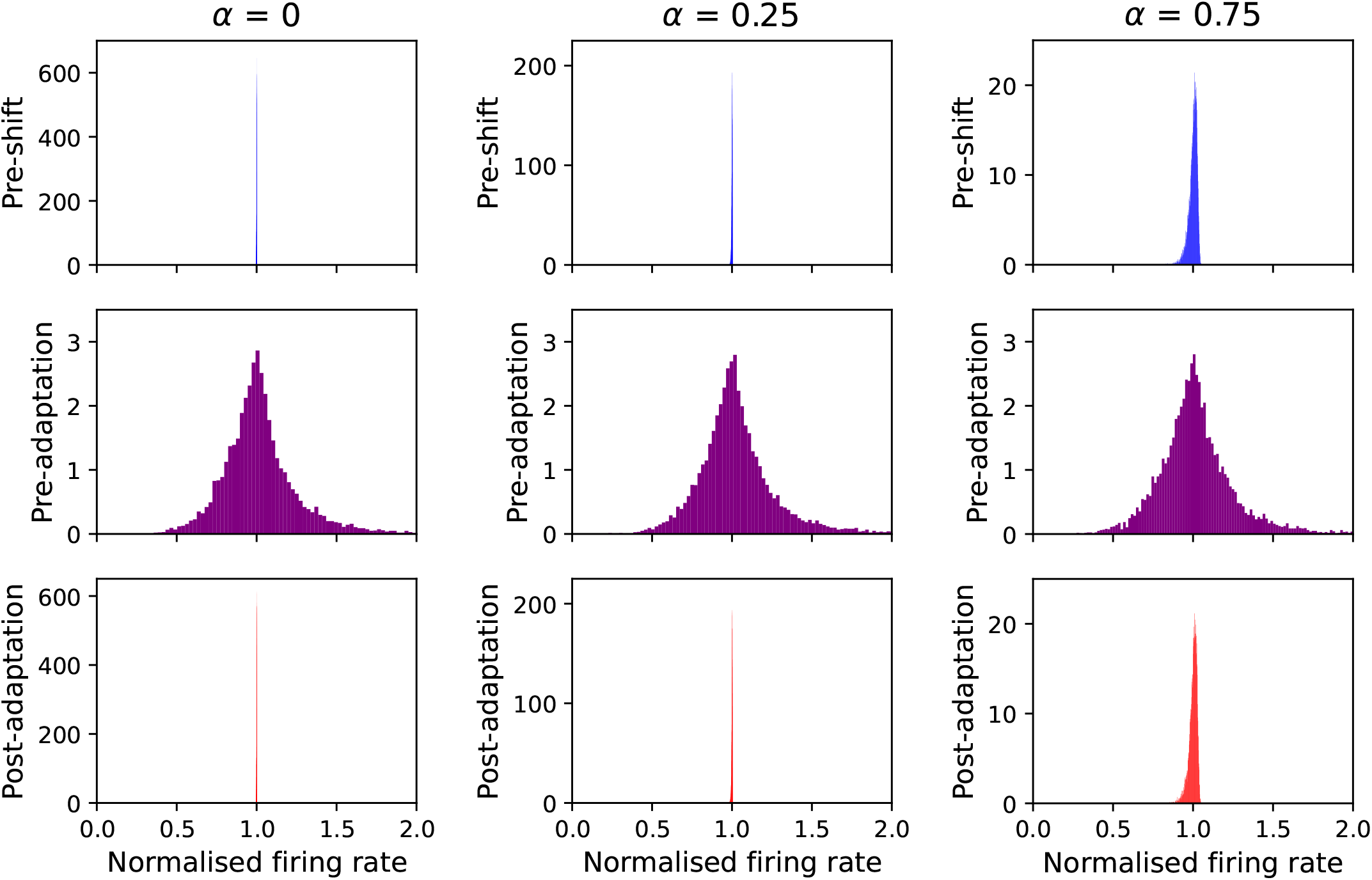
Distribution of average firing rates before and after a discrete environmental shift for the uncorrelated power-law noise subfamily. Each panel has the same format as in Fig. 5B, but for different values of the noise scaling parameter *α*. The case *α* = 0.5 is shown in Fig. 5B.

**Figure 15:**
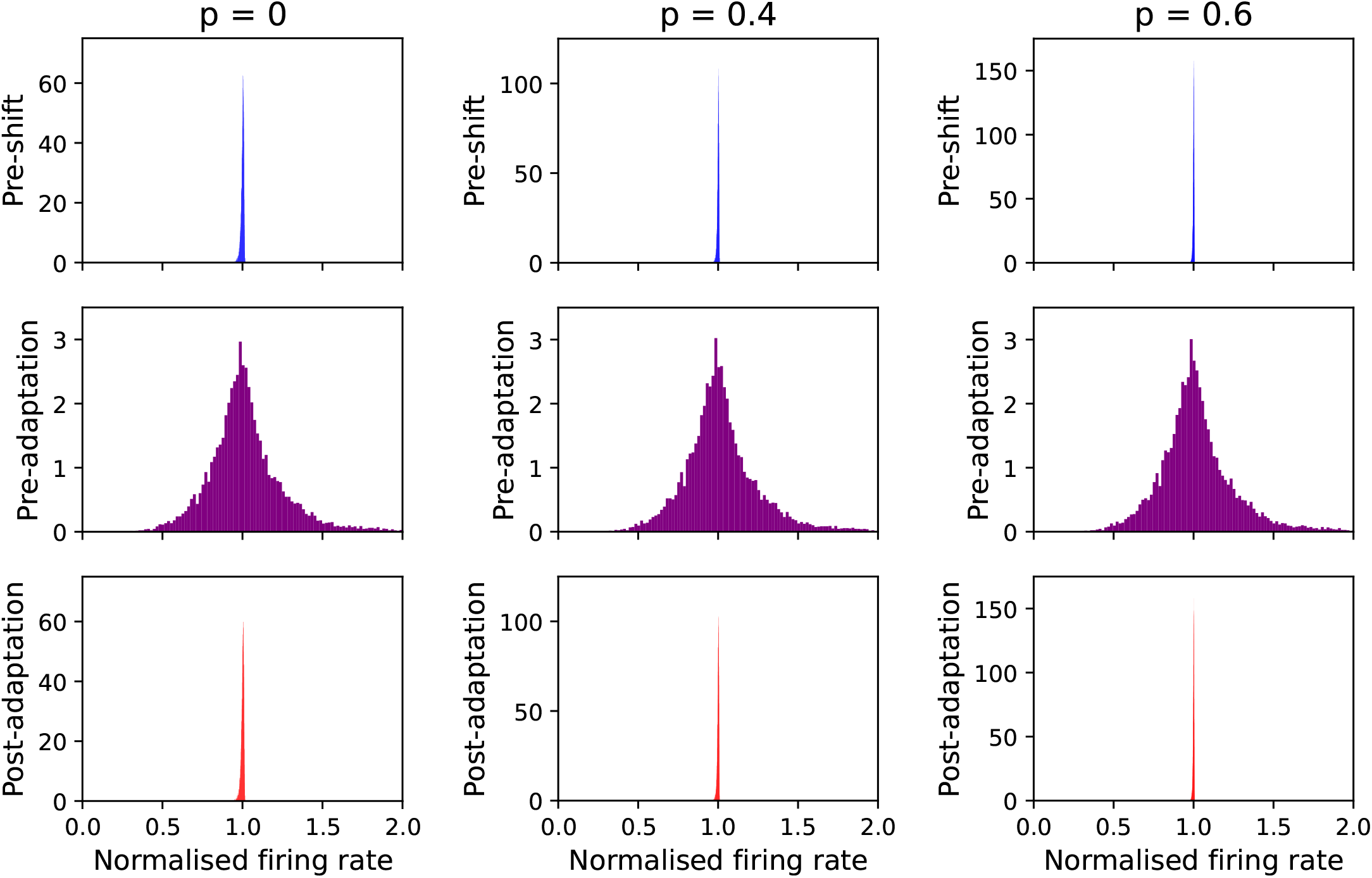
Distribution of average firing rates before and after a discrete environmental shift for the Poissonian noise subfamily with uniform noise correlations. Each panel has the same format as in Fig. 6B, but for different values of the noise correlation coefficient *p*. The case *p* = 0.2 is shown in Fig. 6B.

**Figure 16:**
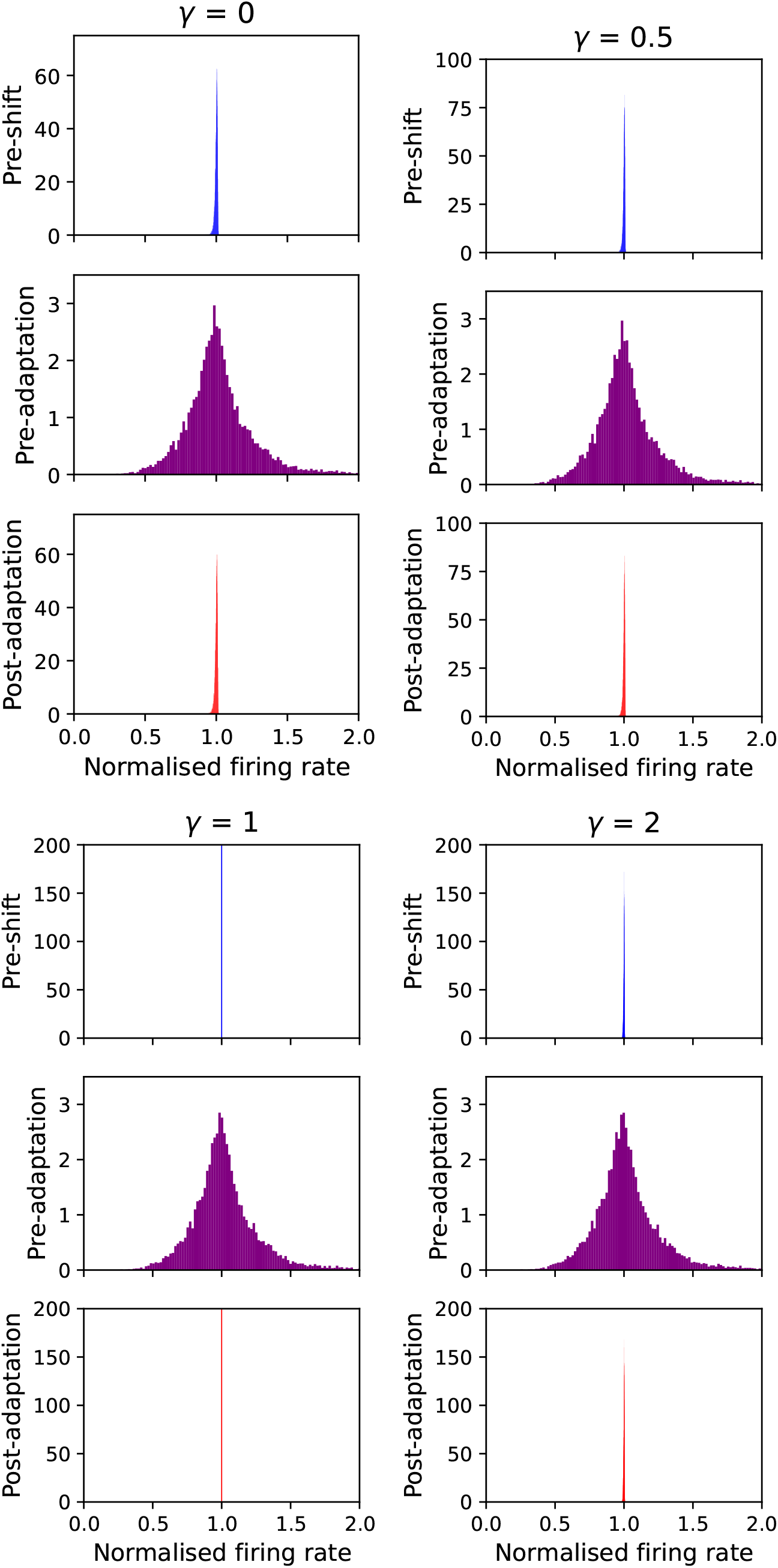
Distribution of average firing rates before and after a discrete environmental shift for the aligned Poissonian noise subfamily. Each panel has the same format as in Fig. 7B, but for different values of the noise spectrum decay parameter *γ*. The case *γ* = 1.5 is shown in Fig. 7B.

**Figure 17:**
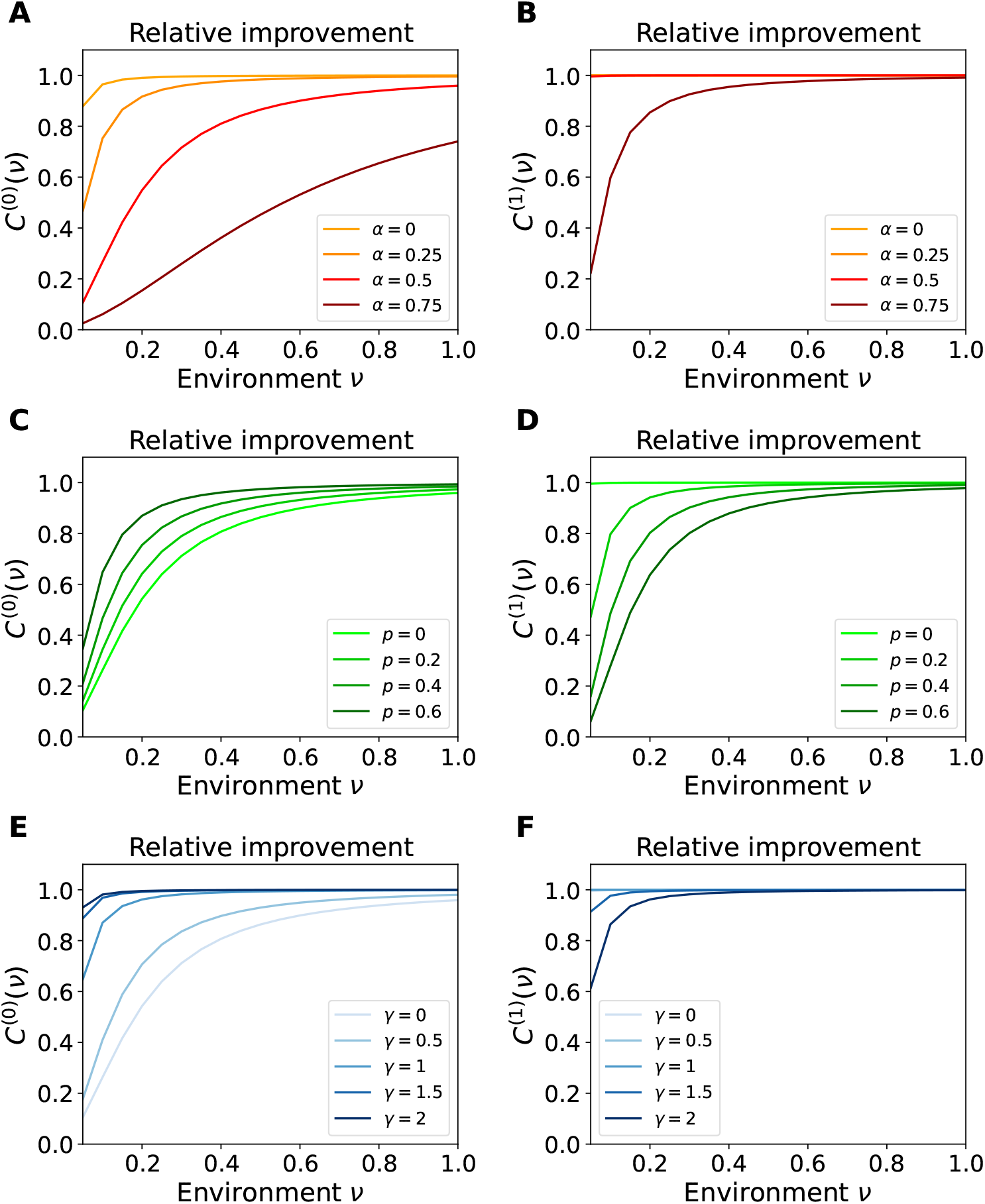
Accuracy of homeostatic approximation to optimal gains. The plots in the panels A, C, and E show the relative improvement in the efficient coding objective for the zeroth-order approximation for the optimal gains ***g***^(0)^, Eq. (11), for the three noise sub-families; see the caption of Fig. 9 for a description of panels B, D, and F there, respectively. Plots in panels B, D, and F here similarly show the relative improvement in the efficient coding objective for the first-order approximation ***g***^(1)^, Eq. (106), for the three noise sub-families.

**Figure 18:**
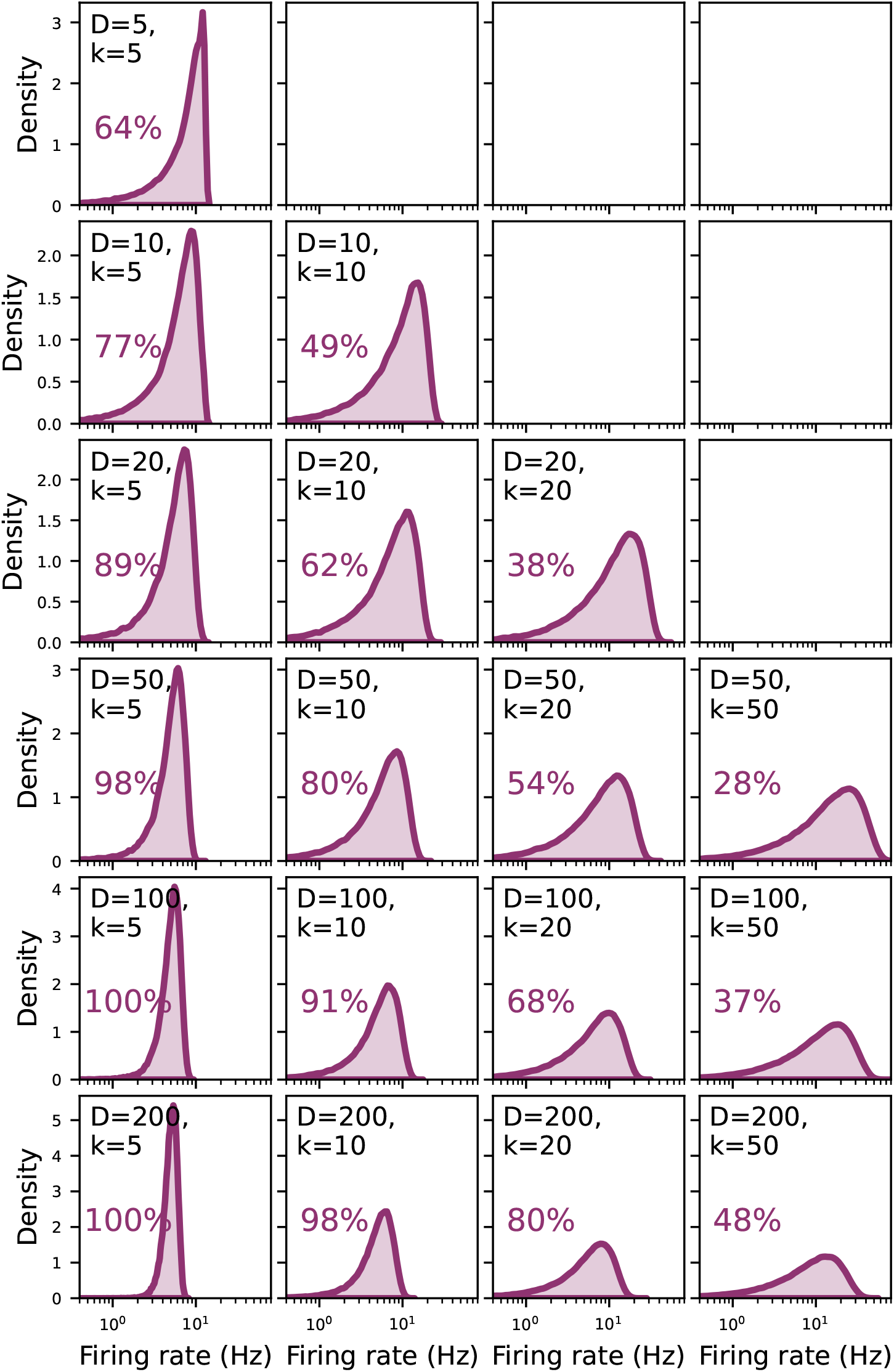
Systematic exploration of how the firing rate distribution of individual neurons varies as the statistical properties of cluster responses change. The panels here have the same description as those in Fig. 8, but show the single-neuron firing rate distributions for different choices of cluster size *k* and effective tuning curve dimension, *D*. (Note that the upper-right panels corresponding to cases in which *D < k* are left empty; this is because in such cases the covariance matrix is singular, and therefore the quadratic program does not have a unique solution. Such cases are also less relevant biologically given realistic estimates for the size of the similarly tuned neurons vs. the effective dimensionality of tuning curves, noting that the nominal dimension of the tuning curve function space is infinite.)

## B Supplementary Information: mathematical derivations

### B.1 Analytic expression for ℒ

In this appendix we derive an analytic expression for the upper bound objective ℒ. We adopt the noise model Eq. (6), repeated here for convenience:

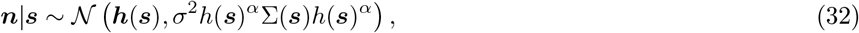

where Σ(***s***) is the stimulus-dependent noise correlation matrix, and 0 *< α <* 2 is a scaling parameter. Here we have made use of our convention that the non-bold version of a vector (without indices) denotes the diagonal matrix formed from that vector, so that, for example, *h*(***s***) is the diagonal matrix with *aa*-th entry equal to *h*_*a*_(***s***).

We decompose the mutual information between the spike counts and the stimulus as

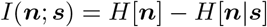

and then obtain ℒ by replacing the term *H*[***n***] in 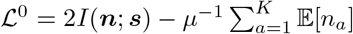 with the entropy of a Gaussian with equal covariance to ***n***. Doing so gives us the expression

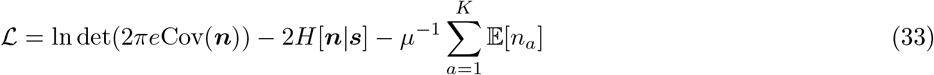

We consider ℒ as a function of the rates *r*_*a*_ = *ω*_*a*_*g*_*a*_ in order to keep the derivation clean.

We start by deriving an expression for the noise entropy *H*[***n***|***s***]. We use *≐* to denote equality up to additive pure constants or terms independent of *r* and *g*. We have

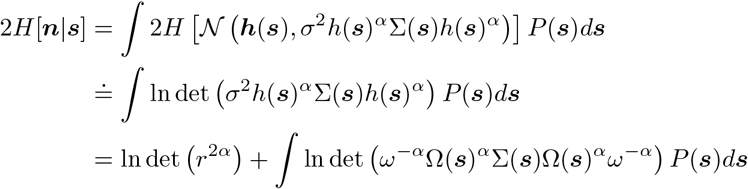

(where in deriving the third line we used *h* = *g*Ω = *rω*^−1^Ω), hence

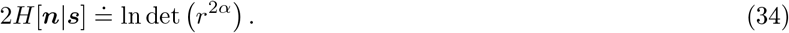

Next, we find an expression for Cov(***n***), using the orthogonal decomposition Cov(***n***) = 𝔼[Cov(***n*** |***s***)] + Cov(𝔼[***n***|***s***]). For the first term have

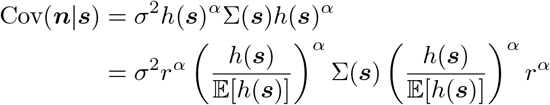

where we used *r* = 𝔼[*h*(***s***)]. Thus

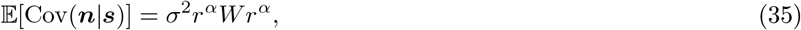

where we have defined (c.f. Eq. (8) of main text) the normalised stimulus-averaged noise covariance matrix *W* as

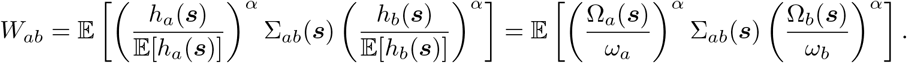

Next, we find an expression for Cov(𝔼[***n***|***s***]). Using ***h***(***s***) = 𝔼[***n***|***s***], we find

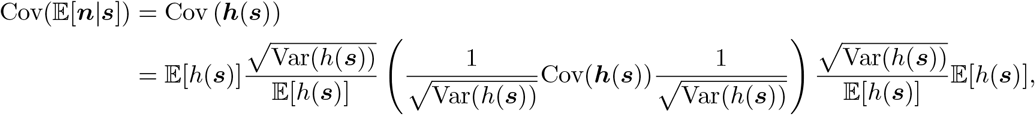

or

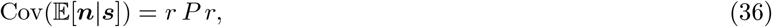

where we defined

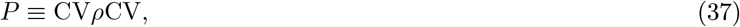

and introduced the coefficients of variation (CV) of trial-averaged responses, given by *h*(***s***) (or equivalently, the CV’s of the representational curves)

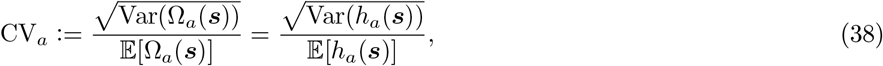

and the Pearson’s correlation matrix of *h* (or equivalently of the Ω’s), defined by

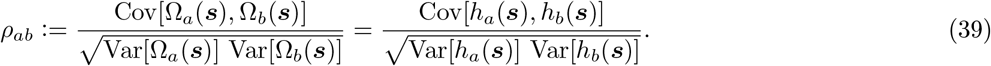

Thus *ρ* is the matrix of signal correlations.

Putting Eq. (35) and Eq. (36) together, we obtain

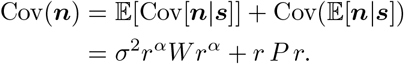

Finally, plugging this result and Eq. (34) into Eq. (33), up to an additive constant, we obtain

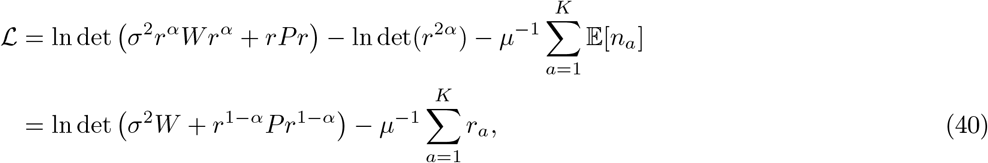

as required. Substituting back in *r*_*a*_ = *ω*_*a*_*g*_*a*_ gets us Eq. (9).

### B.2 Upper bound for Poissonian noise

In this appendix, we consider the following model for cluster spike counts.

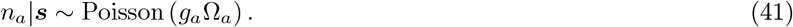

Here we derive a Gaussian upper bound to the mutual information, and show that an approximation to it leads to the same expression Eq. (9) for ℒ derived for the Gaussian noise case, for *σ*^2^*W* = *I* and *α* = 1*/*2 (*i*.*e*., uncorrelated unit-Fano-factor noise). We start with the objective

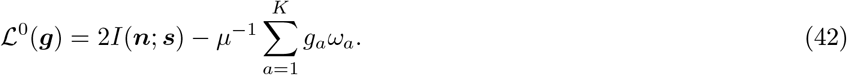

We decompose *I*(***n***; ***s***) = *H*[***n***] − *H*[***n***|***s***]. We will once again upper bound the marginal entropy *H*[***n***]. A difficulty arises from the fact that a Poisson random variable is discrete and the Gaussian upper bound we previously used is for a continuous random variables. We address this problem as follows: Consider a random variable ***U*** which is uniformly distributed on [0, 1)^*K*^, independent of ***n***. Then ***n*** + ***U*** is a continuous random variable. We apply the Gaussian bound to this. Let *p* be the p.d.f. of ***n*** + ***U***, *P* the p.m.f. of ***n***, and *u* the p.d.f. of ***U*** .

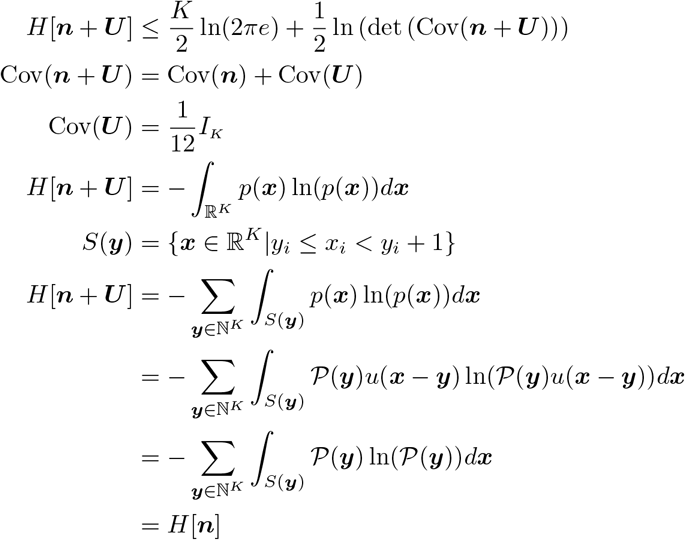

Putting this together gives us the upper bound

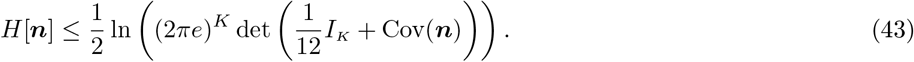

We now address the problem of the marginal entropy *H*[***n***|***s***]. By conditional independence, we have that

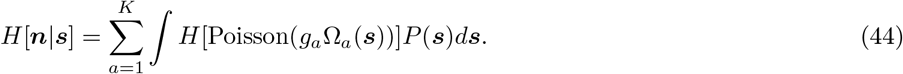

We now make use of an additional assumption, namely that the representational curves Ω_*a*_ have a baseline, and in particular

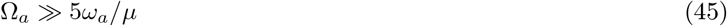

everywhere. Under this condition we can make the approximation that for fixed ***s***,

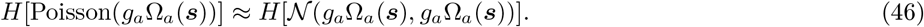

This means we can obtain the approximate upper bound:

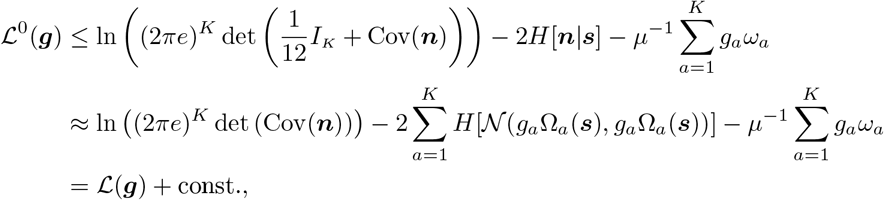

where ℒ (***g***) is the same functional that we defined earlier for Gaussian random variables, in the case of uncorrelated, unit-Fano-factor noise, Eq. (9). We have made an additional approximation by neglecting the *I*_*K*_*/*12 term, which relevant parameter conditions is negligible compared the covariance.

### B.3 Mathematical treatment of the clustered population

In this section we consider a clustered population in which the representational curves are a small perturbation to cluster-wide representational curves. Let *C*_*a*_ denote cluster *a* (or rather the set of indices of single neurons belonging to that cluster). Then for *i* ∈ *C*_*a*_,

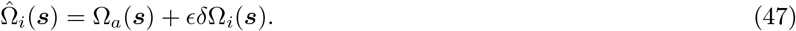

where *ϵ* is a small parameter controlling the deviation from perfect within-cluster similarity. We use hatted symbols 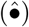 to denote single-neuron quantities (defined analogously to Eqs. (7)–(8) but for single neurons, with Ω_*a*_(***s***) replaced by 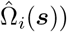, and corresponding no-hat symbols for either cluster averages or zeroth-order values of tuning curves, etc, that are uniform across neurons in a cluster and hence only depend on cluster indices. In particular, we use *P* = CV*ρ*CV, defined in Eq. (37), at cluster level (hence a *K × K* matrix), and use 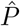 to denote the equivalent matrix defined for individual neurons (an *N × N* matrix). As we noted in the main text Sec. 2.3, for this analysis, we limit ourselves to the case of uncorrelated noise, Σ(***s***) = *I*, with Poisson-like scaling *α* = 1*/*2. Mirroring Eq. (40), the population level objective in this case can be written as

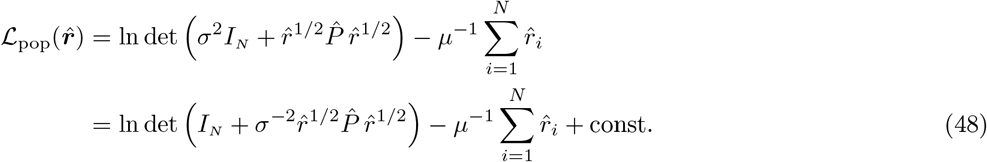

In the rest of this section we will set *σ* to 1 by absorbing it into 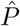; at the end, we can replace all 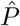—or factors proportional to them— by 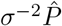 to recover the general case).

#### B.3.1 Expansion in *ϵ*

To simplify the mathematical derivation, we will assume that clusters are the same size *k* = *N/K*, and that w.l.o.g. the population is sorted so that neurons in the same cluster appear adjacent to each other in the ordering. However, the assumption of equal size clusters is not essential, and our final results are valid for the case of clusters of variable size as well. To zeroth order in *ϵ*, the elements of 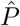 are constant over blocks corresponding to clusters; in other words

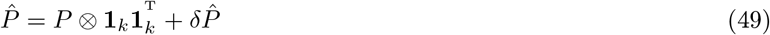

for some deviation 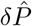 that is *O*(*ϵ*) (here **1**_*k*_ denotes the *k*-dimensional vector with all components equal to 1). However, we will find that the first non-zero corrections to the loss (for *ϵ >* 0) arise from *O*(*ϵ*^2^) corrections to 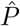 (or equivalently to 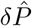 as defined by Eq. (49)). We will therefore (1) expand ℒ to second order in 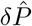, (2) (in the next section) expand 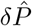 to second order in *ϵ*, and (3) substitute 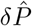 in the loss and eventually neglect terms that are *o*(*ϵ*^2^).

We start by expanding ℒ. Using Eq. (56), and plugging Eq. (49) into Eq. (48) and expanding to second order in 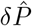, we obtain

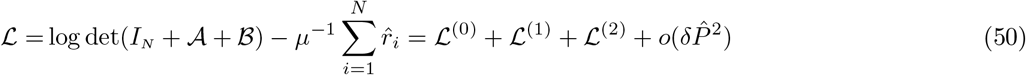

where

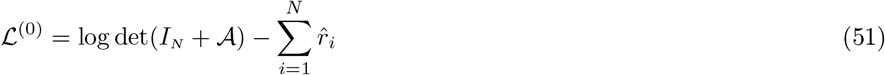

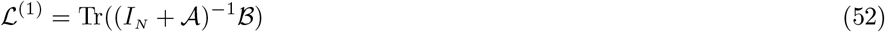

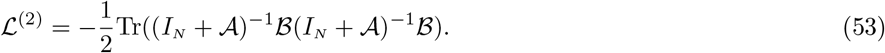

and we defined the following matrices:

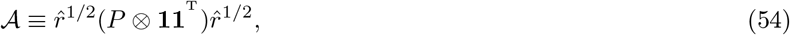

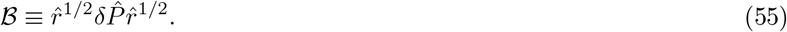

Here and in the rest of this appendix we use **1**, instead of **1**_*k*_, to denote the *k*-dimensional vector with all components equal to 1.

#### B.3.2 Cluster level problem at the zeroth order

We now show that the zeroth order loss, Eq. (51), depends only on the cluster rates 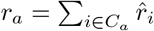. This objective has two components: the information term and the energetic term. For the energetic term the claim is obvious, as we have

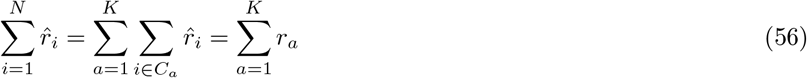

where *r*_*a*_ denotes the total rate of cluster *a*. As for the information term, and will show that

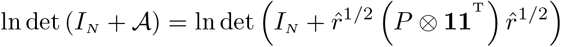

can be re-written (up to a constant) as

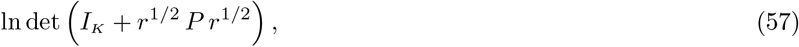

reducing the problem to a cluster-level one (the second determinant is of a *K × K* matrix, rather than *N × N*, and only depends on cluster rates *r*_*a*_ and the zeroth-order signal correlations encoded in *P*). Let us denote the Cholesky factor of (the positive definite) *P* by *U*, such that *P* = *UU*^T^. We thus have

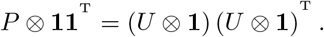

Further defining 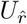 to be the *N × K* matrix 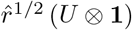, we find

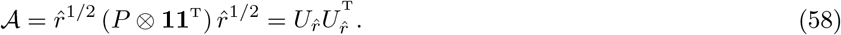

By the matrix determinant lemma we then obtain

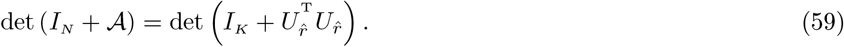

Finally,

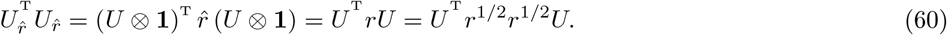

Using this and making use of the matrix determinant lemma one more time, we see that the determinant on the right hand side of Eq. (59) can be written as Eq. (57).

Putting together the information and energetic terms (and momentarily returning the *σ*^2^), we obtain that (up to an additive constant)

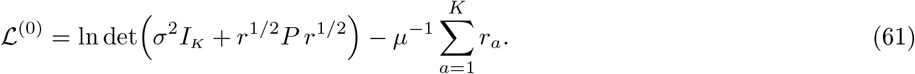

#### B.3.3 Expansion of 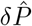 **in** *ϵδ*Ω

We will denote *ϵδ*Ω_*i*_ above by *V*_*i*_ in this subsections (so *V* = *O*(*ϵ*)), and will use *δ*’s to denote deviation from expectation: *δX* = *X* − 𝔼[*X*]. We have

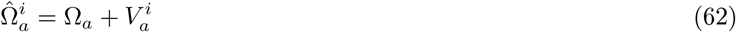

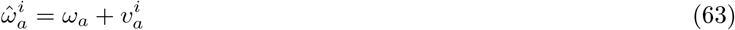

where 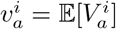. We also have

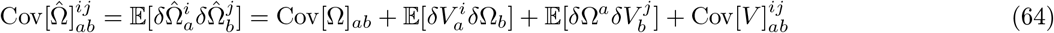

or (keeping cluster indices explicit but hiding witihn-cluster ones)

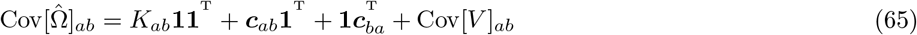

where we defined the vector ***c***_*ab*_ to have components

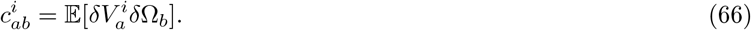

and

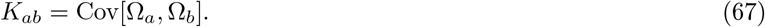

Since

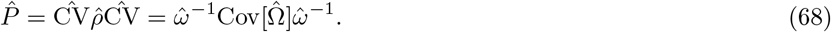

we also expand the diagonal factors to second order in *V* :

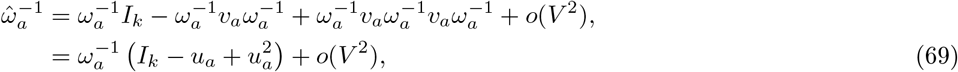

where we defined

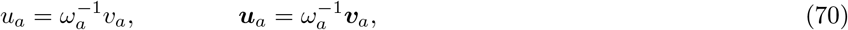

(note that *ω*_*a*_ are numbers, but *ŵ* _*a*_, *v*_*a*_ and *u*_*a*_ are *k × k* diagonal matrices).

First we consider the first order (in *V* or equivalently in *ϵ*) contributions to 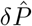. Using Eq. (65) and Eq. (69) in Eq. (68) we have

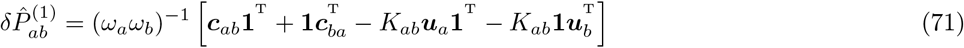

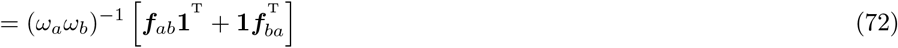

where we defined

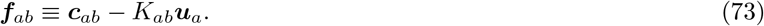

Next, we obtain the second order contributions to 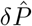. Using Eq. (65) and Eq. (69) in Eq. (68), we obtain

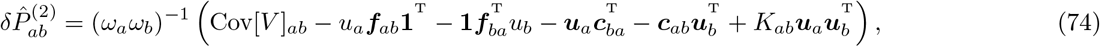

and in particular

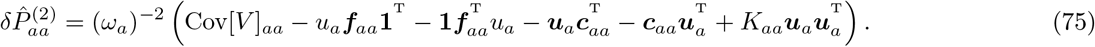

#### B.3.4 *δ*Ω_*i*_ corrections to the loss

We now consider the contributions of the first and second order corrections to 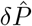 to the loss, via Eqs. (52) and (53): to calculate corrections to the loss to *O*(*ϵ*^2^), we need to plug 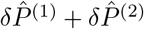 into *ℒ* ^(1)^, and plug in 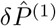 (only) in ℒ ^(2)^. Firstly, consider the matrix inverse that appears in expressions Eqs. (52)–(53) for ℒ ^(1)^ and ℒ ^(2)^. Using 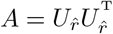 from Eq. (58) and the Woodbury matrix identity, we can write

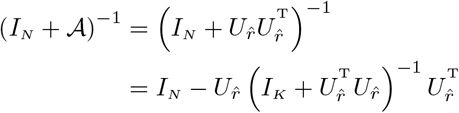

Using 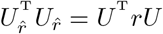 –see Eq. (60)– we find

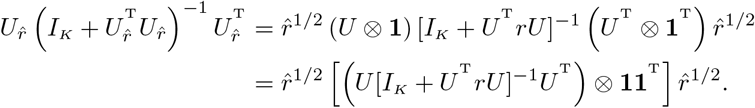

Assuming *P* is full-rank (the generic case), we have *U* [*I*_*K*_ + *U rU*]^−1^*U* = ((*U*^−1^)^T^ *U*^−1^ + *r*) ^−1^ = (*P*^−1^ + *r*)^−1^ yielding

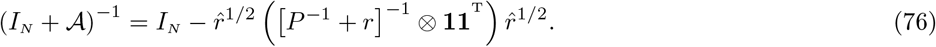

Up until this point, all expressions we have obtained have been exact (apart from the *ϵ*-expansion itself). We now make an approximation to the full perturbation objective which is valid in the high signal-to-noise ratio regime as outlined in Sec. 2.4. Specifically, we will take the matrix

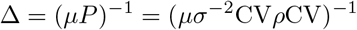

to be small, and expand to zeroth order in this matrix. Δ represents the high-dimensional structure of signal-to-noise ratio. As we show in App. B.4, to zeroth order in this matrix, *r* = *µI*_*K*_. Using the approximation

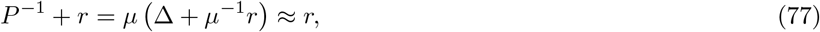

valid to zeroth-order in Δ, in Eq. (76) we obtain

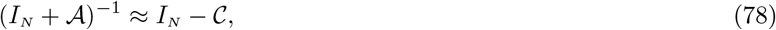

where we defined

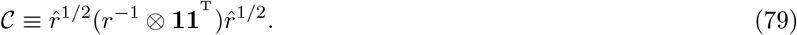

Note that in writing Eq. (77) we have assumed that cluster rates are *O*(*µ*), so that the second term in the parenthesis is *O*(1) and dominates Δ; this will be justified *a posteriori*, by the homeostasis result for the inter-cluster problem (as the approximate optimiser of Eq. (61), in the small Δ regime).

We will show that 𝒞 and hence *I* − 𝒞 are projection operators and we will characterise the latter’s kernel (*i*.*e*., the vectors annihilated by it). First 𝒞 is clearly a symmetric matrix. So we just need to show that 𝒞^2^ = 𝒞. To prove this (and other statements), we note that products of *N × N* matrices (or their products with *N*-dimensional vectors) can be written in terms of products of their blocks (corresponding to the clustering of neurons) as follows: 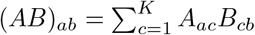 where the subscripts index blocks, with *a, b* ∈ {1, … *K*}, and *A*_*ac*_ and *B*_*cb*_ are multiplying as matrices. As we will see our results do not truly rely on the tensor product structure we have assumed for various matrices (here 𝒜 and C); but only on their constancy within blocks defined by clusters. *Thus our results generalise to the case where clusters contain different number of neurons*. For simplicity, however, we will stick to the tensor structure, corresponding to the same number of neurons in different clusters. Using the above observation, we can write (note that below *r*_*a*_ and *δ*_*ab*_ are scalars, while 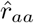 denotes the *a*-th diagonal block of 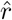 and is a diagonal matrix)

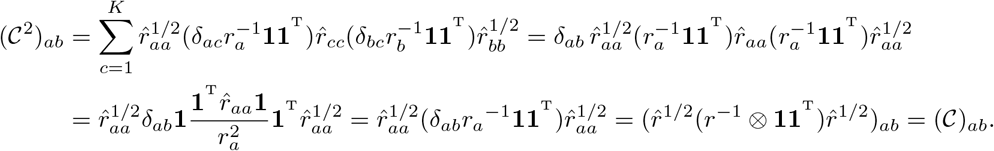

where we used 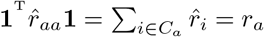.

Similarly for vectors of the form ***ŵ*** = ***w*** ⊗ **1** (namely vectors with components that are constant over each cluster), we have 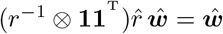, as

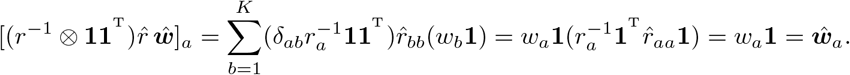

(note that *w*_*a*_ are scalars). Thus if ***ŵ*** is any such vector, then 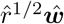 is left invariant by 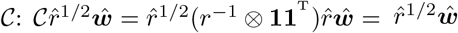. Finally, this means that 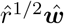 is in the kernel of *I* − *𝒞* = (*I* + *𝒜*)^−1^ and is annihilated by it. Moreover, since 𝒞 is symmetric, row vectors of the form ***w***^T^ ⊗**1**^T^ are annihilated when multiplied by 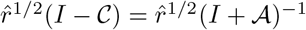 on the right. Finally, matrices with cluster blocks that have uniform rows (columns) are annihilated when multiplied by 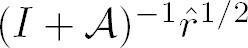 (by 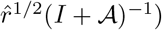 on the left (right).

It follows immediately that, given the structure of Eqs. (52)–(54), any of the terms in the expressions Eqs. (72)–(85) for blocks of 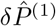 and 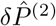 that have **1** as an outer product factor contribute nothing to *δ*ℒ. In particular, the *O*(*ϵ*) part, 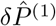, makes no contribution to the loss. Since we are interested in leading order corrections, it thus suffices to only consider the contribution of 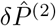—after dropping terms involving **1** in Eq. (85)— as it enters ℒ ^(1)^ (in the Δ ≪ 1 limit). Denoting this correction by *δL*, from Eqs. (52)–(54) and Eq. (78) we find

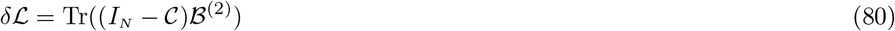

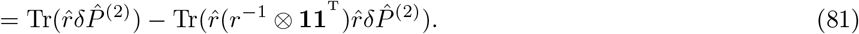

Traces of *N × N* matrix products can be written in terms of traces of products of their blocks as Tr(*AB*) = ∑_*ab*_ tr(*A*_*ab*_*B*_*ab*_) where tr denotes trace over *k × k* blocks. We get a further simplification because 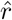 is diagonal and 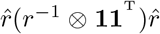 is block-diagonal (due to the diagonality of 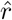 and *r*), with the diagonal blocks given by 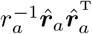. But if *D* is a block-diagonal matrix, then Tr(*DB*) = ∑ _*a*_ tr(*D*_*aa*_*B*_*aa*_). Thus

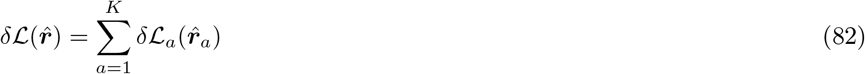

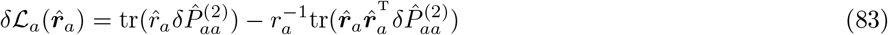

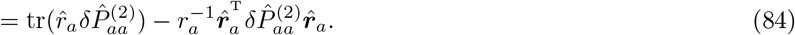

In other words, the correction to the loss is a sum of terms, each of which depends only on the rates in one of the cluster only. As a result, given the total cluster rates *r*_*a*_, the optimisation of rates of single neurons decouples across clusters. Which means we can optimise the rates in cluster *a* under the constraint that they sum up to *r*_*a*_, to maximise Eq. (84). Using Eq. (85) we now express *δ* ℒ _*a*_ explicitly.

#### B.3.5 Within-cluster loss to leading order in *ϵ*

From Eq. (75), after dropping the terms with factors of **1**, we have

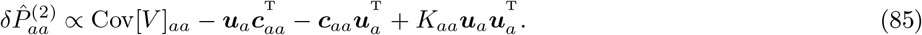

We further ignored the factor 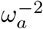 which multiplies both terms in the loss and is thus irrelevant to the optimisation; we will similarly ignore a factor of *µ* in the loss below. The right hand side of the above expression is the covariance of the zero-mean vector (of random variables)

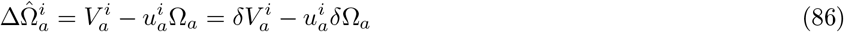

(as can be easily checked: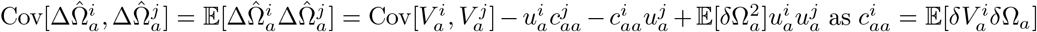).

Since the problem has decoupled across clusters we will now drop the cluster index *a*; in particular we will denote the unperturbed Ω_*a*_ by Ω_0_, and we will also denote the given/fixed total cluster rate, *r*_*a*_, by *R*_0_. Substituting the above in Eq. (84) and ignoring irrelevant overall prefactors, we find that the single-neuron rates in a cluster with total rate *R*_0_ are solutions of the following quadratic programming problem:

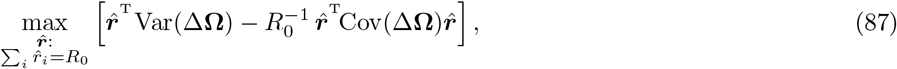

where

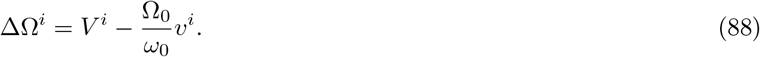

This is equivalent (up to replacement of *R*_0_ with *µ*) to the optimisation problem, Eq. (28), of the main text. Finally, note the scaling property of this quadratic programming problem: if we scale the cluster rate, *R*_0_, by some *α*, the solution 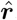, and thus the rates of individual neurons in the cluster, scale by the same factor. Note also that the scaling of ΔΩ^*i*^ by a constant (within a cluster) factor does not make a difference, and we can make it “dimensionless” by dividing it by *ω*_0_ if helpful (in fact this amounts to returning the prefactor 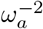 that we dropped after Eq. (85)).

#### B.3.6 A toy model for a cluster’s tuning curves

Here we will develop a toy model for Cov(Δ**Ω**), *i*.*e*., for the statistics of deviations of representational curves of neurons in the same cluster from the unperturbed curve Ω_0_. Assuming a degree of smoothness for Ω^*i*^(***s***) (as a function on the stimulus space), we will adopt a basis of smooth functions *b*_*µ*_(***s***) for 1 ≤ *µ* ≤ *N*_*b*_, and assume that the log tuning curves are in the span of this set of functions. In other words, we assume

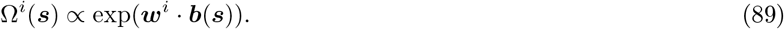

for some vector of coefficients 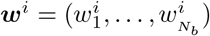. Similar to Eq. (47) we assume 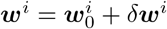 (we absorb *ϵ* into *δ****w***^*i*^). Expanding the above equation we get

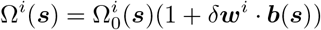

or equivalently

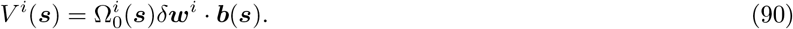

It follows that *v*^*i*^ = 𝔼[Ω_0_***b***] *· δ****w***^*i*^, and thus (from Eq. (88))

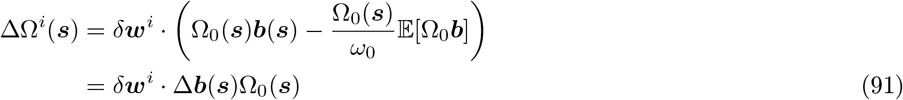

where we defined

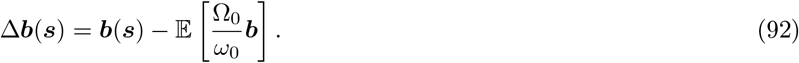

We thus obtain:

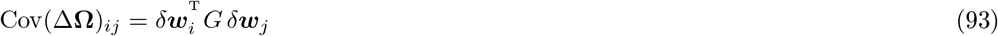

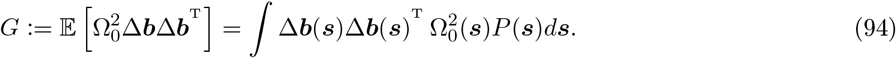

We think of *G* as a metric defining the inner product of the vectors *δ****w***_*i*_. Note also that given the final comment in the previous subsection, we can replace Ω_0_(***s***) inside the integral expression for *G* with Ω_0_(***s***)*/ω*_0_; this is nice as it means that if it happens that the cluster tuning curve Ω_0_ is mostly supported on regions with small stimulus frequency, the above metric does not go to zero. Alternatively, since the scaling of Cov(Δ**Ω**) is irrelevant to the within-cluster rate optimisation, we can assume that the expectation in the definition of *G* is taken, not with respect to *P* (***s***), but with respect to the *normalised* measure with density proportional to 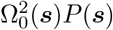. If we denote expectation under this “tilted” measure by 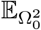 we can also write:

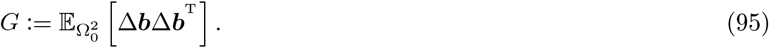

Similarly we could have written

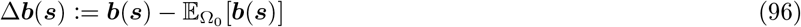

(note that this last expectation is tilted by Ω_0_ and not by 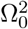). To summarise, we have found that the quadratic form for our quadratic programming optimisation is given by the Gram matrix of the neurons’ coefficients *δ****w***_*i*_ with respect to the inner product defined by *G*.

For our minimal toy model, we will assume that the perturbations *δ****w***_*i*_are isotropic, mean-zero Gaussians, *i.e.*, 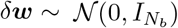. We will additionally assume that the positive-definite matrix *G* has *D* eigenvalues which are roughly equal, and all significantly larger than the other eigenvalues. In this case, (up to scaling) we can approximate *G* as a projection matrix onto *D*-dimensions. This allows us to generate 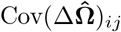 simply as 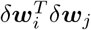, where the vectors 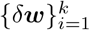 were drawn independently from a *D*-dimensional isotropic Gaussian, *δ****w***_*i*_ ~ 𝒩 (0, *I*_*D*_).

### B.4 First order maximisation of ℒ

In this section we compute the maximiser of ℒ, to first order in the parameter Δ, defined by **??**:

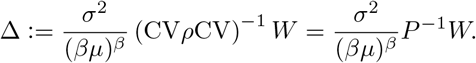

To simplify the derivation we will work with as a function of the rates *r*_*a*_ = *g*_*a*_*ω*_*a*_, rather than the gains. Writing Eq. (9) in terms of *r*_*a*_, we have

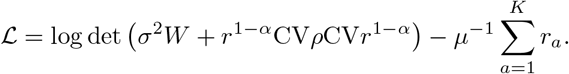

We will also introduce the matrix

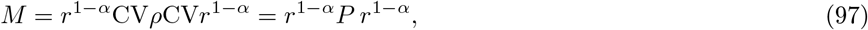

so that

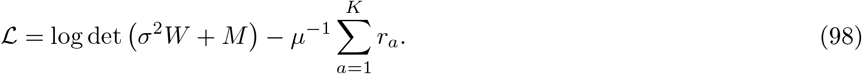

Note that

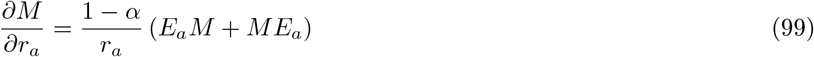

where *E*_*a*_ is the diagonal matrix which is zero everywhere except having 1 as the *a*-th element along the diagonal.

Using Eq. (98) and Eq. (99), we obtain that

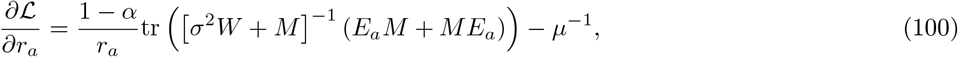

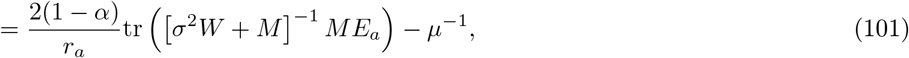

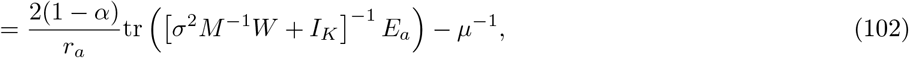

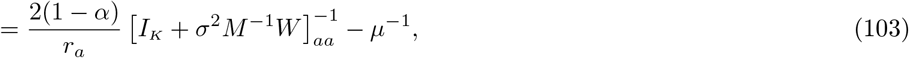

where we used the cyclicity of trace and the symmetry of *M* and *W*. Setting the derivative in Eq. (100) to zero and rearranging, and defining *β* = 2(1 − *α*), we obtain

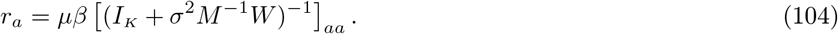

We now apply the ansatz

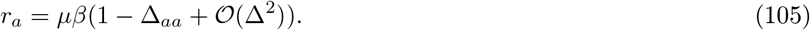

Plugging this into Eq. (97) we have that

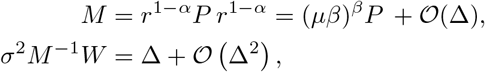

which after substitution in the right hand side of Eq. (104) yields

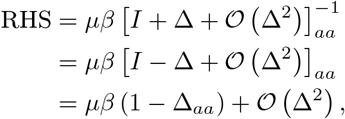

where on the penultimate line we have used the Neumann expansion of *I* [+ Δ + 𝒪 (Δ^2^)] ^−1^. Thus to first order in Δ, the optimal solution is given by *r*_*a*_ = *µβ*(1 − Δ_*aa*_). In terms of the gains, this corresponds to the solution

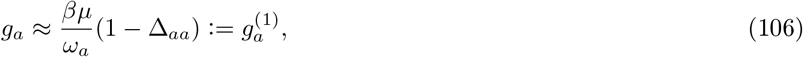

which also yields Eq. (11) in the limit Δ → 0.

### B.5 Aligned noise leads to perfect homeostasis and uniformisation

In this appendix we demonstrate that (for constant coefficient of variation) if *ρ* and *W* are perfectly aligned *i*.*e*.,, *ρ* = *W*, then the optimal rates are all equal and invariant of the environment, *i*.*e*., we have perfect homeostasis and uniformisation.

In App. B.4 we derived an equation for the maximiser of the objective function ℒ (***r***), Eq. (104),

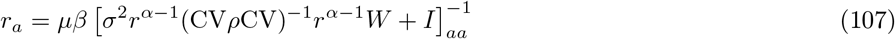

We now show that, when *W* = *ρ* and CV is constant, this equation is satisfied by constant ***r*** = *χ***1**_*k*_ for some *χ*, and find an equation for *χ*.

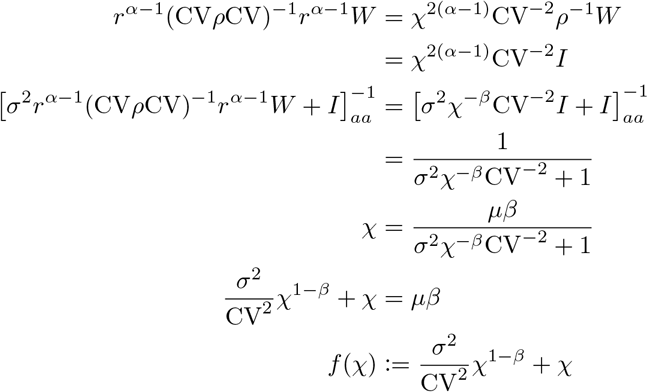

We will now show that this equation has a solution, provided certain conditions on the parameters hold. We divide into two cases. If *β* ≤ 1 then *f* (*χ*) is strictly increasing and limits to 0 as *χ* → 0 and infinity as *χ* → ∞. By continuity we must have *f* (*χ*) = *µβ* at some point.

If *χ >* 1 then we still have *f* (*χ*) → ∞ as *χ* → ∞. We now find the minimum of *f*. Taking derivatives, we see that this occurs at

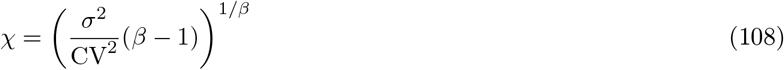

At this point, we have

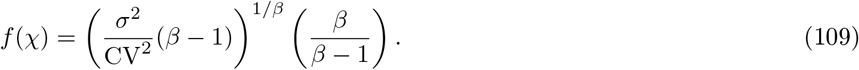

This is less than *µβ* provided that

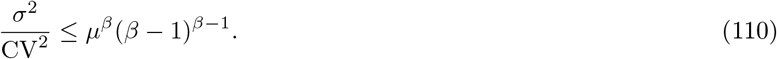

So provided Eq. (110) holds, there is a solution by continuity. Recall that *µβ* is approximately the average firing rate of a cluster. Typically we are interested in holding the product of these parameters constant at a value *r*. If we do so, we can take the derivative of the right hand side of Eq. (110) in *β*, obtaining that this expression is minimised when *β* = *r/*(*r* − 1), where it attains a value of *r* − 1. Putting this together, we can say that there exists a uniform, homeostatic solution provided

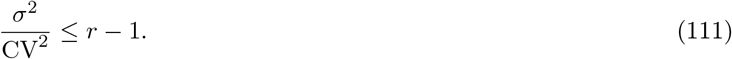

where *r* is the product *βµ*.

### B.6 Estimates for 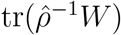

We define *A*_*γ*_ to be the normalisation constant of a correlation matrix with an eigenspectrum proportional to 1*/n*^*γ*^. Since a *K × K* correlation matrix has trace *K*, we have

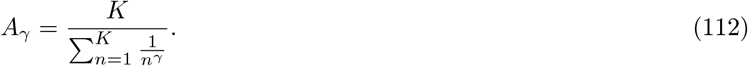

We start with the case *γ*≠ 1. Then we have that

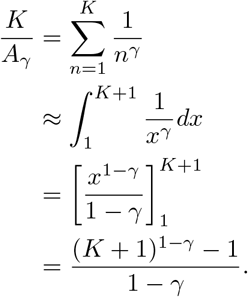

Now, taking the limit as *γ* → 1, we obtain that

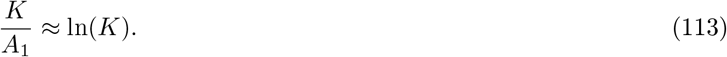

Now when *γ <* 1 and *K* is large, we obtain

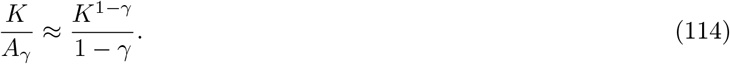

Lastly, when *γ >* 1 and *K* is large, we obtain

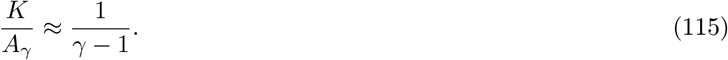

These approximations allow us to arrive at the following estimates. Firstly, consider tr(*ρ*^−1^)*/K*.

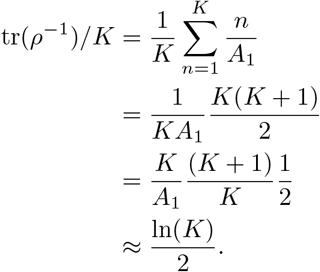

Next, consider the case of aligned noise, when *W* has a *A*_*γ*_*/n*^*γ*^ eigenspectrum and aligned eigenbases with *ρ*. In this case,

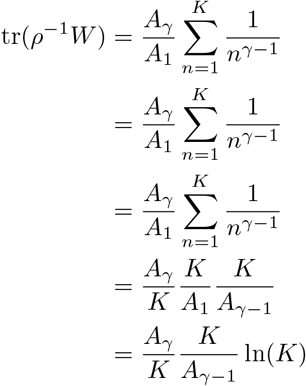

We can then derive the following approximations for tr(*ρ*^−1^*W*)*/K* using Eqs. (113)–(115).

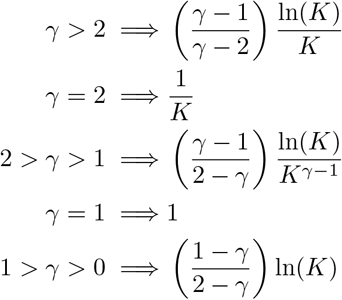

### B.7 Optimal homeostatic gains

We now consider the case where we enforce homeostasis on the gains. Prior to taking expectations across environments, our objective, considered as a function of *r*_*a*_ = *g*_*a*_*ω*_*a*_ is (up to additive constants)

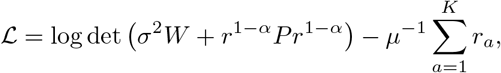

where *P* = CV*ρ*CV. Enforcing homeostasis at the cluster level means setting 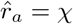 for all clusters. Substituting this in, we obtain the function

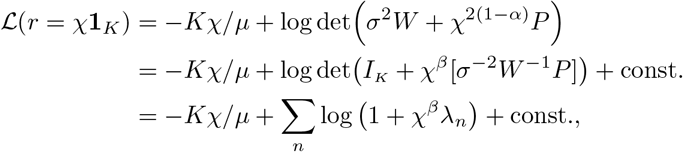

with *λ*_*n*_ are the eigenvalues of *σ*^−2^*W*^−1^*P*. Note that this is only a function of the spectrum, and not of the eigenbasis. Therefore, under the assumption that the spectrum remains fixed across environments, we can drop the need to take expectations, and work with ℒ as a function only of the spectrum. We work under this assumption going forward.

The optimal *χ*, within this family of approximate solutions, obeys

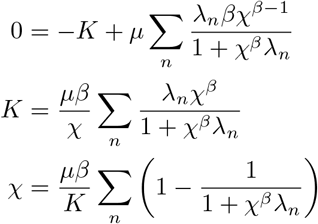

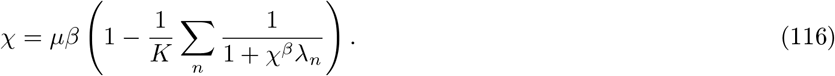

We now demonstrate that, under appropriate conditions, 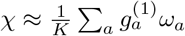

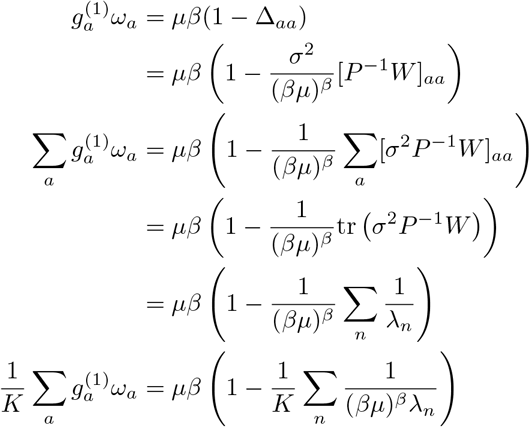

Now, under the assumption that (*βµ*)^*β*^*λ*_*n*_ ≫ 1 for each *n*, we can say that *χ* ≈ *µβ* and (*βµ*)^*β*^*λ*_*n*_ ≈ *χ*^*β*^*λ*_*n*_ + 1. Substituting this in gives us the required result. Note that this condition is equivalent to the high signal-to-noise ratio condition we have used throughout.

We now consider the special cases discussed in Sec. 2.2 in which more precise solutions can be obtained. Note that in all three cases the analytics correspond to idealisations of the numerical simulations we actually perform in Sec. 2.6.

#### Uncorrelated power-law noise

In the first special case under consideration, *W* = *I*_*K*_, CV^2^ is constant, and *ρ* has approximately a *A*_1_*/n* eigenspectrum, where *A*_1_ is chosen to normalise the trace of *ρ* to be equal to *K, i*.*e*.,

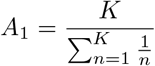

The spectrum of *σ*^−2^*W*^−1^*P* is therefore *λ*_*n*_ = *b/n* where

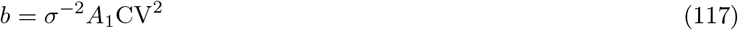

In this case, (defining *u* = *n/K*) we can make the following approximation to the right hand side of Eq. (116):

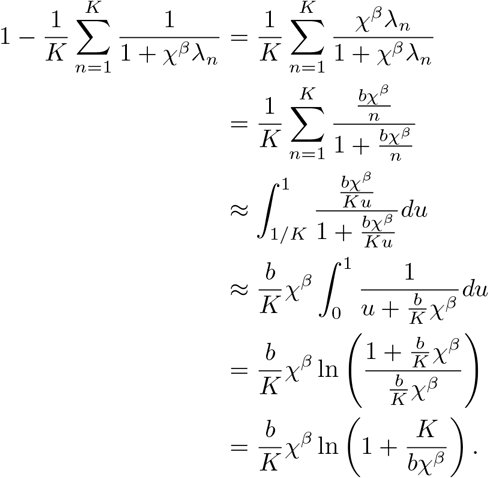

This gives the new equation

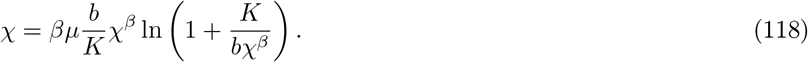

Approximating *A*_1_ *≈ K/* ln(*K*) (see App. B.6) and using Eq. (117), we obtain that

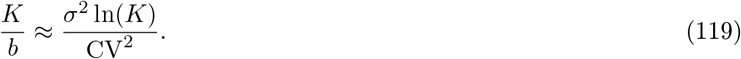

Substituting Eq. (119) into Eq. (118) gives us

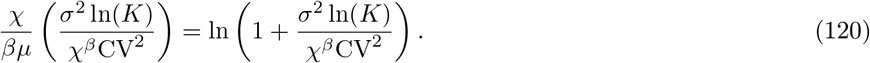

#### Aligned noise

In the aligned noise case, we approximate *ρ* and *W* as having the same eigenbasis. *ρ* still has an eigenspectrum of *A*_1_*/n*, and we take *W* to have an eigenspectrum of *A*_*γ*_*/n*^*γ*^ where *A*_*γ*_ normalises the trace of *W* to be equal to *K*,

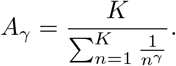

The matrix *σ*^2^*W*^−1^*P* therefore has eigenspectrum *bn*^*γ*−1^ where *b* = *σ*^−2^*A*_1_CV^2^*/A*_*γ*_. Inserting this into Eq. (116) gives us

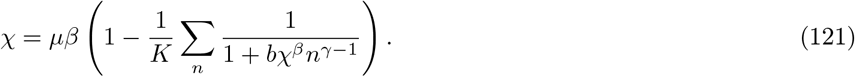

See App. B.6 for estimates of *A*_*γ*_.

#### Constant correlation noise

The next special case occurs when *β* = 1, and *W* = (1 − *p*)*I*_*K*_ + *p***11** ^T^. Using the Sherman-Morrison formula, we obtain

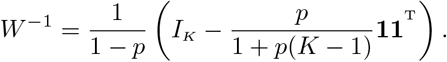

Since 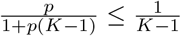 we neglect this term and approximate 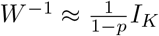. This therefore has the same effect as making the replacement *σ*^2^ ↦ (1 − *p*)*σ*^2^. Substituting this into Eq. (120), and using *β* = 1, we obtain

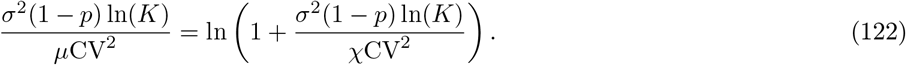

We define

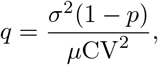

and rearrange to get

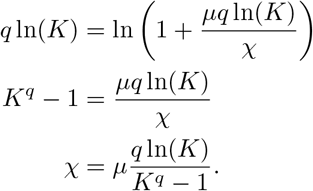

### B.8 Synaptic normalisation allows for propagation of homeostasis between populations

In this appendix, we derive how the weights between neural populations must change in order for homeostasis to be propagated between them (see the last paragraph of Section 2.6). We work in a linear (or linearised) rate model.

Consider two neural populations, with an upstream population providing feedforward input to a downstream population. We will suppose that the upstream population is engaged in homeostatic coding, and ask what is necessary for the downstream population to be also engaged in homeostatic coding. We will denote the tuning and representational curves of the upsteam population by 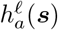 and 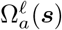, respectively, where *ℓ* = 1 (*ℓ* = 2) for the upstream (downstream) population or layer. We drop corrections to the zeroth order solution for optimal gains, and work within the homeostatic coding regime; that is we assume the gains of the upstream population are given by

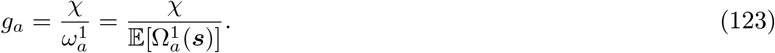

The downstream population of tuning curves will be given by 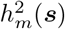 for *m* = 1, …, *M*. Let *W*_*ma*_ be the synaptic weight from neuron *a* in the upstream population to neuron *m* in the downstream population. Working in a linearised rate model, this gives us that the tuning curve of neuron *m* is

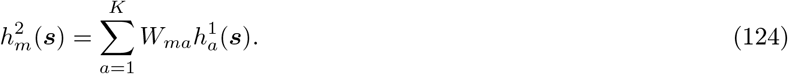

Suppose that, as dictated by the computational goals of the circuit, the downstream population has representational curves 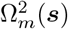 for *m* = 1, …, *M*. We similarly write each representational curve as a linear combination of the upstream cluster representational curves,

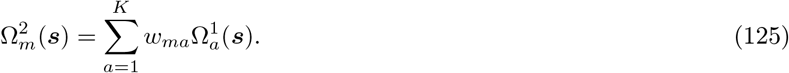

We call these weights *w*_*ma*_ the *computational weights*, since they are determined by the computational goals of the downstream population. In line with our approach so far, we treat these computational goals, and hence also the computational weights, as given (*i*.*e*., set independently of optimal gain adaptation).

If the downstream population is also implementing homeostatic coding, we know that

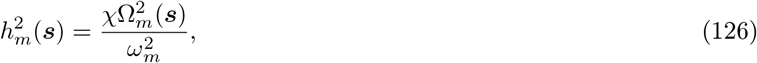

where 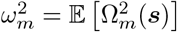. We asked how the synaptic weights *W*_*ma*_ should depend on the computational weights *w*_*ma*_, and how they should adapt as stimulus statistics change in order for Eq. (126) to hold, *i*.*e*., for the downstream population to also engage in homeostatic coding. But then

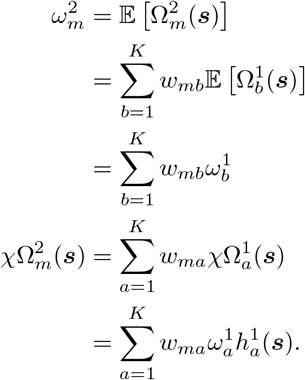

Substituting this in, we get

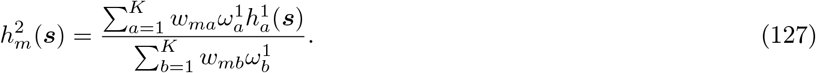

Comparing coefficients, we can see that

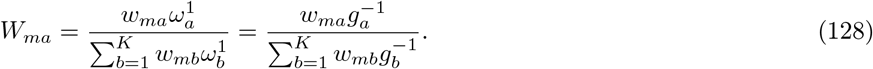

In other words, this scheme requires that the synaptic weights are normalised such that the total synaptic weights received by each downstream neuron remains constant, independently of adjustments of the gains of the pre-synaptic neurons or of possible changes in the computational weights.^14^ Thus, homeostatic coding, applied sequentially to two populations, provides an additional normative interpretation of synaptic normalisation, in which synapses onto a neuron are jointly scaled to keep total input weights constant (Turrigiano et al., 1998; Turrigiano, 2008).

### B.9 Hierarchical Bayes-ratio coding

In this appendix, we consider homeostatic propagation (as defined in the Appendix B.8) in the specific case of a feedforward net implementing Bayes-ratio coding. In particular, we calculate and interpret the feedforward synaptic weights needed for this propagation. We start with a two-layer generative model ***z***^2^ → ***z***^1^ → ***s*** (with joint density given by *P* (***s, z***^1^, ***z***^2^) = *π*_2_ (***z***^2^)*f*_2→1_(***z***^1^|***z***^2^)*f*_1→0_ (***s***|***z***^1^)), and consider a two-layer feedforward recognition network that inverts that generative model. Recall that in Bayes-ratio coding, the representational curves are given by the posterior distribution; in particular, in the second layer of the recognition network these are given by posteriors of the higher level variable ***z***^2^:

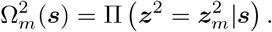

We first show that the second layer’s representational curves can be written as a linear combination of the first layer’s representational curves, *i.e.*, 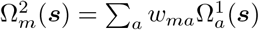. To find the proper value of *w*_*ma*_ (which would lead to the correct inversion of the generative model), note that due the Markov structure of the generative model, the correct posterior factorises as Π(***z***^1^, ***z***^2^|***s***) = Π(***z***^2^|***z***^1^)Π(***z***^1^|***s***). Thus Π***z***^2^|***s*** = Π(***z***^2^|***z***^1^)Π(***z***^1^|***s***)*d****z***^1^, and

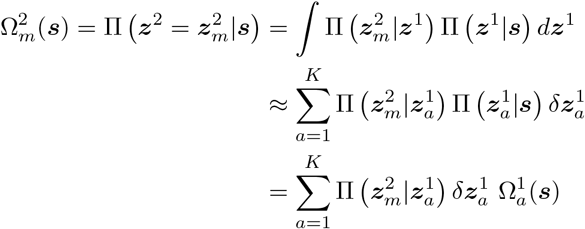

hence

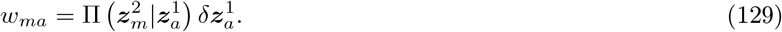

Recall from Appendix B.8 that the feedforward synaptic weights, *W*_*ma*_, needed for homeostatic propagation are given by Eq. (128), in terms of the “computational weights”, *w*_*ma*_. Using Eq. (129) and that (in an ideal observer model) 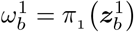 (*i*.*e*., the average posterior over ***z***^1^ is equal to their prior distribution) we obtain

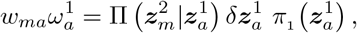

and

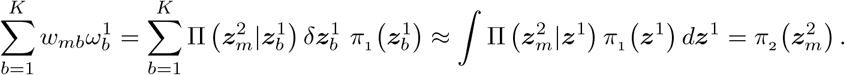

Plugging these in Eq. (128), and using Π(***z***^2^|***z***^1^) *π* (***z***^1^) = *f* _2→1_(***z***^1^|***z***^2^) *π*_2_ (***z***^2^), we obtain

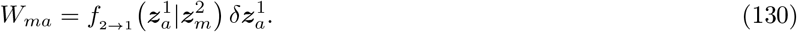

Thus a linear feedforward network with these feedforward weights correctly propagates Bayes-ratio coding to implement the exact inversion of a two-layer generative model. By induction, this result can be extended to *n*-layer generative models with Markov structure ***z***^*n*^ → … ***z***^1^ → ***s***, and their corresponding Bayes-ratio encoding recognition models furnished by *n*-layer linear feedforward nets.

### B.10 Stimulus specific adaptation for non-ideal-observer models

In this appendix, we show how discrepancies between the internal model and the external environment lead to simultaneous stimulus specific and neuron specific adaptation effects in a homeostatic DDC code. We begin with the special case of a Bayes-ratio code, which is a DDC in which the kernel functions are delta functions (as discussed in Sec. 2.8). In a Bayes-ratio code, representational curves are given by

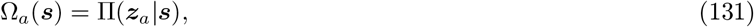

where Π is the posterior distribution under the internal generative model. We consider the case of a non-ideal-observer generative model, in which case there will be a discrepancy between the marginal stimulus distribution predicted by the model, *P*^*I*^ (***s***) and the external environment stimulus marginal *P*^*E*^(***s***). We reason as follows:

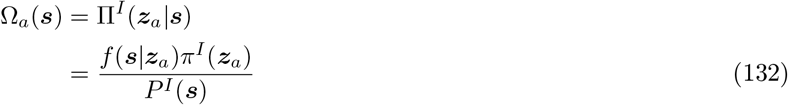

where

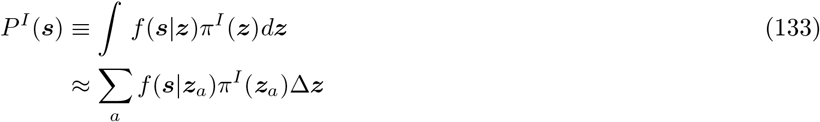

According to homeostatic coding, the tuning curve is given by

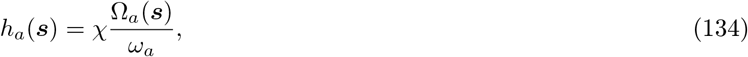

where *ω*_*a*_ is the temporal average of the representational curve, or equivalently, the expectation of Ω_*a*_(***s***) under the *true* environmental stimulus distribution. Thus

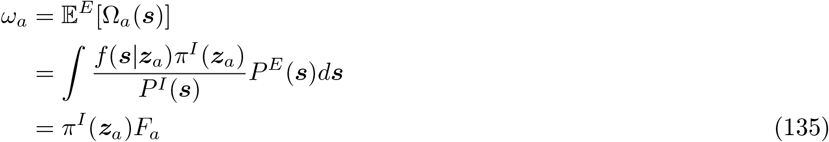

where we defined

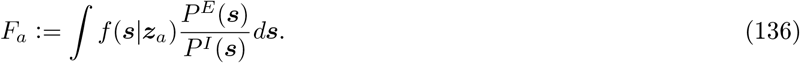

Notice that when *P*^*E*^ = *P*^*I*^, as would be the case for an ideal-observer, we will have *F*_*a*_ = 1, due to the normalization of *f* (***s***|***z***). Substituting Eq. (132) and Eq. (135) into Eq. (134) we obtain

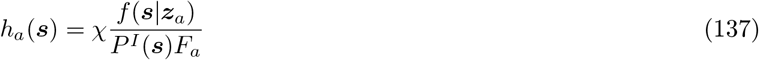

as claimed in Eq. (22).

We now consider the more general case of a DDC code. We define

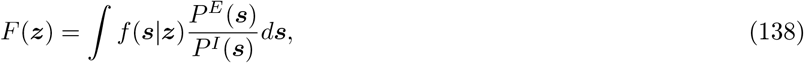

so that *F*_*a*_ in the above is equal to *F* (***z***_*a*_). An analogous derivation yields:

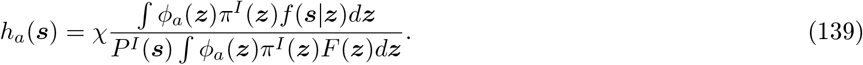

We can see that in this case there is no clean separation of neuron specific and stimulus specific factors. In particular, the generalization of Eq. (24) takes the form:

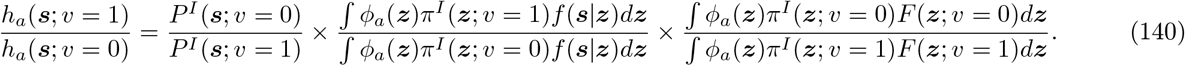

In the limit as the width of the kernels becomes small, the integrals involving kernels collapse to sampling at a single point, as occurs with the Bayes-ratio code. When this happens, we get full separation to stimulus and neuron specific effects; otherwise these effects are mixed together, with more mixing occurring the larger the width of the kernels. However, when the kernels are unimodal and sufficiently narrow, we would expect an approximate factorization of the effect of adaptation into a stimulus-specific and a neuron-specific suppression factor.

## Footnotes

1 We will use “units” instead of “neurons” to refer to the tuning curves because we will apply the theoretical framework to both cases where the units correspond to single neurons or clusters of similarly tuned neurons.

2 Note that adjusting the gains does not mathematically restrict the computations which can be performed by downstream populations, as adjustments in gains can in principle — and if needed — be compensated for by reciprocal adjustments in readout weights. More formally, if a function, *F*, of the input ***s***, can be computed from the unmodulated Ω_*a*_(***s***), say by a linear combination *F* (***s***) *≈* ∑ _*a*_ *w*_*a*_Ω_*a*_(***s***), then it can equally be represented as a linear combination of the gain-modulated tuning curves, 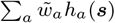, with the adjusted weights 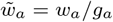.

3 Note that for Poisson-like noise scaling, *α* = 1*/*2, this matrix is actually a correlation matrix, in the sense that it is positive-definite with diagonal elements equal to 1.

4 In terms of the objective function, Eq. (9), this is mathematically equivalent to having zero stimulus-conditioned noise correlations, *i*.*e*., Σ(***s***) = 1, but with CV’s redefined not to denote the standard coefficient of variation, but rather 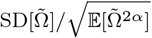 where 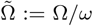; for Poisson-like scaling, *α* = 1*/*2, this reduces to the standard coefficient of variation.

5 The positivity of the noise correlation matrix, *W*, requires that 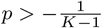. Assuming large *K*, the right side of the inequality goes to zero; hence we only simulated cases with positive *p*.

6 This is a self-consistent statement, as Δ^−1^, more precisely, encodes the high-dimensional SNR structure (which in general depends on the gains) *under the approximate homeostatic solution* Eq. (11). But since largeness of Δ^−1^ justifies that approximation, the largeness of SNR in a population with the homeostatic gains Eq. (11) self-consistently justifies that homeostatic approximation.

7 This is expected to be valid when the eigenbasis of *ρ*^−1^*W* is not aligned with the standard (cluster) basis, which, in turn, corresponds to distributed coding.

8 Further, it can be shown (see App. B.7) that *χ < βµ, i*.*e*., the mean cluster firing rate under the optimal homeostatic solution is strictly smaller than that predicted by ***g***^(0)^, and that the average spike-count of the entire population of clusters is approximately the same under this solution as it is under the first-order approximation *g*^(1)^.

9 For the case of uncorrelated power-law noise model, as well as the aligned correlated noise model, we used an analytical approximation to the optimal *χ* (see App. B.7) used in the homeostatic approximation 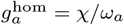, Eq. (14). This means that the performance measures in the plots of Fig. 9B and F are lower-bounds for the performance of 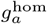 with the truly optimal *χ*.

10 We have assumed the latent variable space is partitioned into cells, one for each neuron, with *δ****z***_*b*_ denoting the volume of the cell centred on neuron *b*’s preferred latent variable ***z***_*b*_.

11 More rigorously, we ought to use a wrapped (periodic) normal distribution. However, since the standard deviations are small compared to the length of the circle, the normalisation constant is approximately 1, and we can treat the density as a normal Gaussian.

12 We can interpret ΔΩ_*i*_ as the direction of the perturbed tuning curve in excess of the cluster tuning curve Ω_*a*_.

13 This is expected to be valid when the eigenbasis of *ρ*^−1^*W* is not aligned with the standard (cluster) basis, which, in turn, corresponds to distributed coding.

14 Different normalisation factors are equivalent to different values of *χ* for different populations. Differences in the optimal choice of *χ* may arise from *e*.*g*., different noise correlation statistics or different rate coding intervals between populations.

